# Classpose drives the discovery of colorectal cancer phenotypes in clinical grade whole slide images

**DOI:** 10.64898/2025.12.18.695211

**Authors:** Soham Mandal, José Guilherme de Almeida, Konstantin Bräutigam, Nickolas Papanikolaou, Trevor A Graham

**Author notes:** Equal contribution.

## Abstract

Cell phenotyping in histopathology samples is essential for diagnostic and research workflows. However, human expert annotation requires significant time and expertise while being affected by inter-observer variability. Here, we present Classpose, an easily trainable framework for cell segmenting and phenotyping built on top of Cellpose-SAM with state-of-the-art performance across 6 distinct datasets, outperforming competing methods. We show that this requires fine-tuning the entire network, highlighting how instance segmentation is a poor objective for downstream cellular classification. We apply it to a large whole slide image (WSI) colorectal cancer (CRC) cohort (SurGen) and show that Classpose-derived cellular organisation and morphology features can be used to determine novel spatial morphological phenotypes for clinically relevant molecular conditions (MMR deficiency, *BRAF* mutations, *KRAS* mutations) and to predict these same molecular conditions. We make Classpose models available and provide a user-friendly QuPath extension for widespread use by the digital pathology community.

## 1 Main

Segmenting cells can yield significant insights through the quantitative estimation and characterisation of cell populations locally and globally. In histopathology, however, the segmentation of individual cells is not sufficient. Multiple cell types can be distinguished with simple stains such as haematoxylin and eosin (H&E), making the identification of individual cell types just as relevant. Cellular abundance, morphological features, and spatial context can reveal critical insights into tissue architecture and disease progression. For instance, clinical outcomes are associated with tumor-infiltrating lymphocytes [1,2,3], tumor-stroma ratios in cancer regions [4], and cancer-associated fibroblasts [5]. Yet, the cell phenotyping required to enable these analyses requires significant domain expertise and time. Even when assisted by state-of-the-art cell segmentation methods, researchers may have to manually annotate many individual cells for adequate cell proportion estimates.

Extensive work has been developed for automatic cell phenotyping in digital histopathology [6], and previous works have highlighted the possibilities afforded by large-scale digital histopathology [7] and foundation models [8,9]. Significant progress has been made in the development of foundation (generalist) models for cell segmentation in bioimaging: Cellpose [10] has pioneered this direction by training a convolutional neural network on large, heterogeneous microscopy image data, enabling robust instance segmentation. This framework has recently integrated the Segment Anything Model (SAM) [11], resulting in Cellpose-SAM (CP-SAM) [9]. This united Cellpose’s prediction objective — combining cell probability and bi-directional distance gradient estimation — with SAM’s pre-trained weights to develop a generalist, state-of-the-art and perturbation-invariant cell instance segmentation framework. CellSAM [12] used a two-step approach: a cell detector predicts bounding boxes for a fine-tuned SAM, which provides the final segmentation masks [12]. These foundational model-based methods excel at performing instance segmentation, but they do not predict cell types. Some efforts have begun to bridge this gap with CellViT [13] and its successor CellViT++ [8], incorporating cell phenotyping into their transformer-based architecture to simultaneously segmentation and classify cells.

In this work, we present Classpose, an expansion of CP-SAM to perform semantic segmentation. we show stateof-the-art performance across multiple datasets in both internal and external validation, with significant differences in detection performance and broadly-defined cell type clusters. We show that full model training is a necessity for optimal performance and provide a QuPath extension for easy Classpose use in digital histopathology workflows. Finally, we show the utility of Classpose by applying it to a large WSI cohort, demonstrating how the spatial and morphological characterisation of Classpose predictions can discover novel hallmarks of different molecular conditions.

## 2 Results

### 2.1 Classpose: from Cellpose to semantic segmentation

Classpose expanded the Cellpose instance cell segmentation framework to semantic cell segmentation using a diverse bag of tricks (Figure 1A). These were motivated through a literature revision which collected lessons from previous works on semantic cell segmentation. These included:

– **Balanced data sampling** — we calculated image sampling weights proportional to the sum of the ratio between the cell-specific proportions and dataset-wide proportions, increasing the likelihood of sampling images with under-represented cell types. We additionally included class weights in the classification loss function.
– **Hybrid loss for better segmentation** — rather than relying on cross-entropy alone, we used a linear combination of the cross-entropy loss with the Tversky loss. The Tversky loss was designed for segmentation, demonstrating good performance, particularly for under-represented classes [14], and leading to state-of-the-art solutions in cellular segmentation [15].
– **Adaptive weighting between losses** — determining their relative importances (i.e. their weights) is a problem common to multi-task learning. Classpose inherited the loss from Cellpose (binary cross-entropy for instance segmentation and mean squared error for flow estimation) and added two additional terms (multi-class cross-entropy and Tversky loss, both for semantic segmentation). The weights usually have to be empirically determined and can be model- or dataset-specific, requiring an extensive number of optimization trials. Solving this, we use uncertainty-driven multi-task learning [16], allowing weights to be adjusted to the linear combination of different loss values which best minimizes loss terms while preventing weights from converging towards zero.
– **Extended data augmentations** — Cellpose uses exclusively random rotations and resizing to perform data augmentations. While impactful [9], we further include random shifts in the hematoxylin-eosin, the hematoxylineosin-DAB and the hue-brightness-saturation spaces, random Gaussian blurring, and additive noise. These augmentations are associated with improved performance in digital histopathology tasks including segmentation and classification [15,17,18].
– **Sparse data support** — data annotation for semantic cell segmentation is resource-intensive, requiring both time and highly specialized professionals as annotations performed by less experienced individuals through crowd-sourcing platforms lead to worse performance [19]. This hard requirement has led to datasets with incomplete annotations (NuCLS, MoNuSAC). Addressing this, while requiring all cell instances to be annotated, we do not require all cells to be classified, enabling the use of automated cell instance segmentation (i.e. Cellpose) followed by the classification of a small fraction of cells across multiple regions-of-interest.
– **QuPath integration** — Classpose features a straightforward Python-based command-line interface for WSI inference; however, we have also integrated Classpose into QuPath [20] as an extension, making it easily and widely available for researchers with differing levels of computational expertise (more details in section S9).

**Fig. 1.**
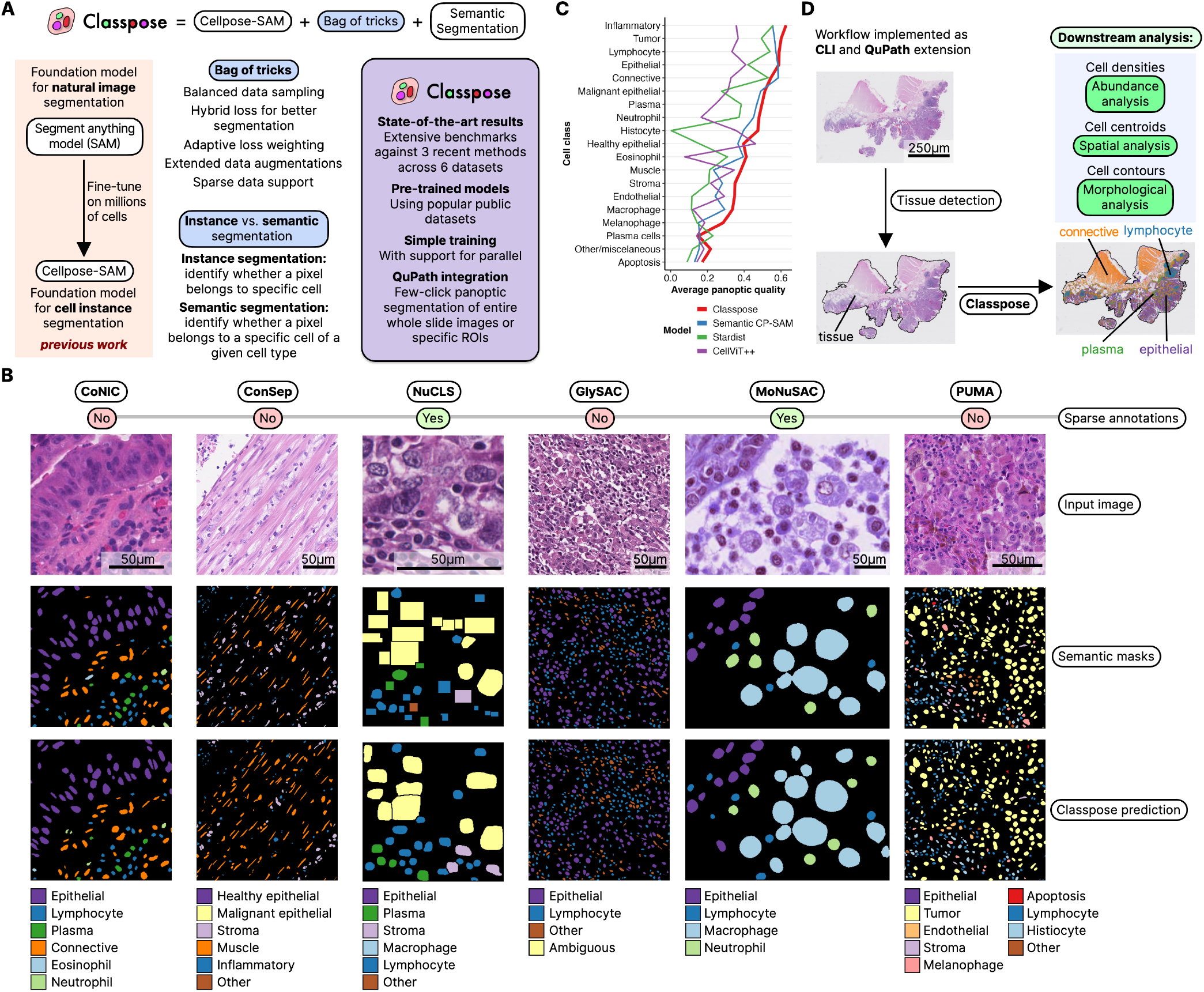
The Classpose framework. **A**. A schematic representation of the development of Classpose. Using Cellpose as a starting point, Classpose included an improved training routine and semantic classification, as well as integration with command line-based and QuPath-based inference. **B**. Examples of annotations and Classpose predictions, together with the complete list of classes used to train each dataset-specific model and whether or not annotations (training or testing) are sparse. **C**. Classpose, Stardist, and CellViT++ average (mean) performance across datasets for multiple cell types. If a cell type is present across multiple datasets, the panoptic quality is the average across datasets. Overlapping cell types (i.e. “Lymphocyte” and “Inflammatory”) are both represented as they are differently annotated across datasets. **D**. Schematic representation of the whole slide imaging prediction workflow. The tissue is detected to reduce the number of tiles requiring prediction, and also detected artefacts as part of the output. We then ran Cellpose across all tissue tiles, generating GeoJSON files by default, with optional CSV and/or SpatialData outputs to support both visualisation and downstream analyses. Tissue detection is performed using GrandQC [21]. A larger representation of the results for this slide is made available in Figure S1.

Through these changes, Classpose is easily trainable across datasets with different levels of semantic annotations, both sparse and dense (Figure 1B). These enhancements enable Classpose to outperform other state-of-the-art methods across multiple datasets and cell types (Figure 1C). Finally, to better serve the wider research community, we have made Classpose models available through HuggingFace ^6^ and offer an intuitive predictive interface for whole slide images, both for the command line and as a QuPath extension (Figure 1D, Figure S1).

### 2.2 Classpose shows state-of-the-art performance

Across multiple datasets, Classpose offers state-of-the-art panoptic quality (PQ) performance, driven largely by improved detection quality and comparable segmentation quality (Figure 2; Table S2; Table S3; Table S4). More concretely, PQ is consistently higher for Classpose when considering datasets which were not present in the Semantic CP-SAM training data. Indeed, in terms of PQ, Classpose is only outperformed by Semantic CP-SAM in CoNIC (present in Cellpose-SAM pre-training data), when classifying miscelaneous (Other) cells in ConSep, and Plasma cells, Macrophages, and miscelaneous (Other) cells in NuCLS. On average, Classpose outperforms Semantic CP-SAM, StarDist and all variations of CellViT++ across all datasets, with CoNIC being the sole exception.

**Fig. 2.**
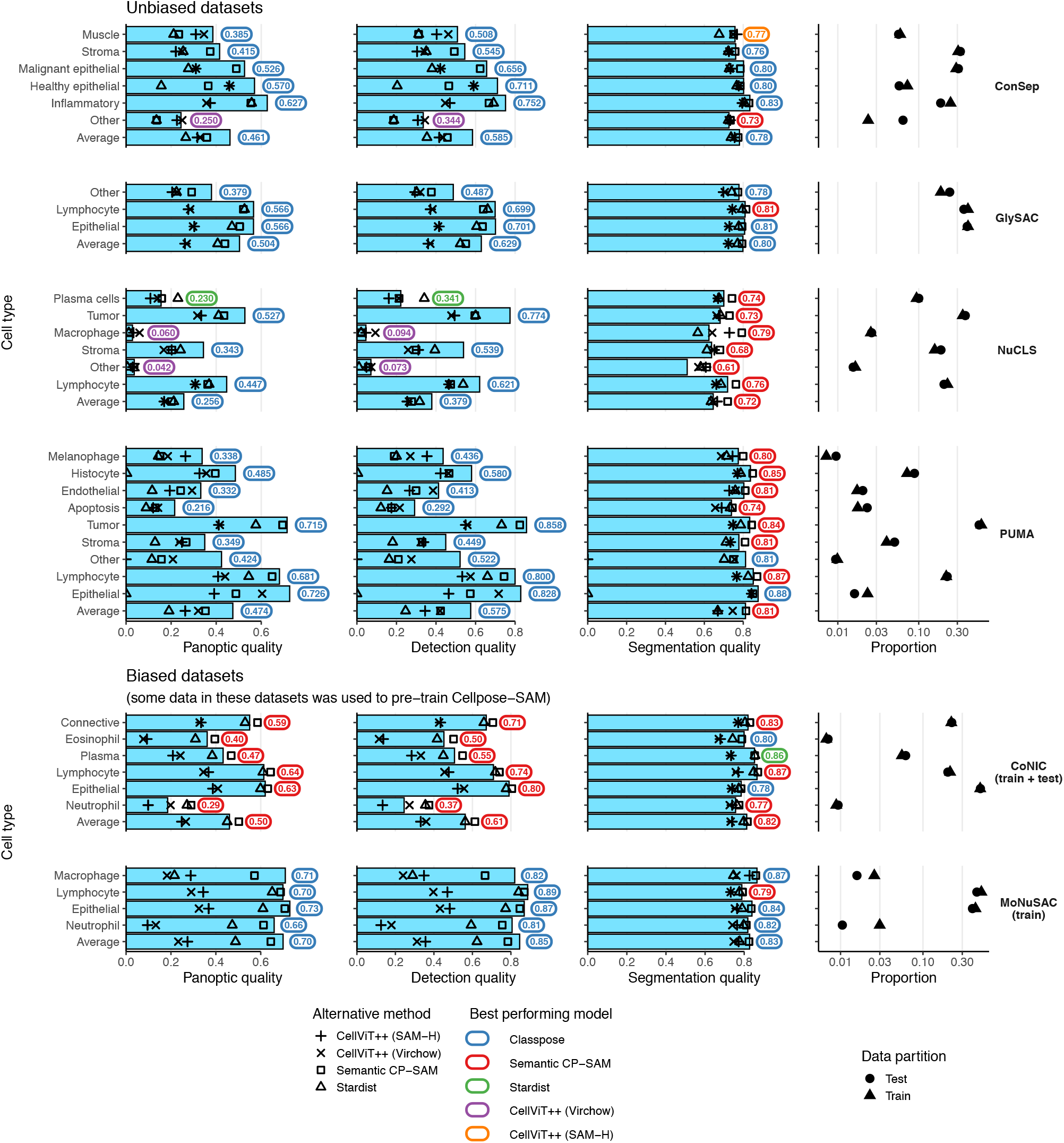
Classpose benchmark against state-of-the-art models, stratified by whether datasets were biased in favour of Classpose/Semantic Cellpose-SAM (training and/or testing data was present in Cellpose-SAM). Panoptic, detection and segmentation quality for Classpose (blue bars), Semantic Cellpose-SAM (CPSAM; squares), the two best performing CellViT++ models (crosses), Stardist (triangles). The best-performing method across each metric is annotated using different colours. The last column shows the relative proportions of each class for each dataset.

We performed a multivariate linear analysis with model (Classpose, Semantic CP-SAM, CellViT++, and Stardist), dataset (ConSep, GlySAC, NuCLS, and PUMA; we excluded CoNIC and MoNuSAC as they are used for CP-SAM pre-training), cell type cluster (Immune, Epithelial, Mesenchymal, Malignant and Other) and interaction terms between model and cell training counts and model and count-decorrelated training proportions (the residuals of the training proportions as a function of the train counts) as predictors for the PQ (Methods, Equation 5). The model coefficients for PQ are presented in Table 1; for DQ and SQ they are presented in Table S5 and Table S6, respectively.

**Table 1.** Coefficient estimates and their respective standard errors (Std. Error), t-values and statistical significance for a two-sided t-test for the linear model described in Equation 5 for PQ. CP-SAM = Cellpose-SAM.

| Variables | Estimate | Std. Error | t-value | Pr(>\|t\|) |
| --- | --- | --- | --- | --- |
| Intercept | 0.76 | 0.03 | 25.89 | < 0.0001 |
| **Model** |  |  |  |  |
| Semantic CP-SAM | -0.07 | 0.01 | -7.34 | < 0.0001 |
| Stardist | -0.13 | 0.01 | -14.15 | < 0.0001 |
| CellViT++ | -0.20 | 0.01 | -21.33 | < 0.0001 |
| **Dataset** |  |  |  |  |
| GlySAC | -0.10 | 0.03 | -3.55 | 0.00052 |
| NuCLS | -0.23 | 0.03 | -8.71 | < 0.0001 |
| PUMA | -0.11 | 0.03 | -3.95 | 0.00011 |
| **Cell clusters (vs. Malignant)** |  |  |  |  |
| Immune | 0.00 | 0.01 | 0.24 | 0.81269 |
| Malignant | -0.07 | 0.01 | -4.74 | < 0.0001 |
| Mesenchymal | -0.14 | 0.01 | -12.14 | < 0.0001 |
| Other | -0.07 | 0.01 | -6.22 | < 0.0001 |
| **Proportion effects** |  |  |  |  |
| Classpose × res<sub>p<sub>train</sub></sub> | 0.26 | 0.06 | 4.67 | < 0.0001 |
| Semantic CP-SAM × res<sub>p<sub>train</sub></sub> | 0.43 | 0.06 | 7.64 | < 0.0001 |
| Stardist × res<sub>p<sub>train</sub></sub> | 0.38 | 0.06 | 6.76 | < 0.0001 |
| CellViT++ × res<sub>p<sub>train</sub></sub> | -0.20 | 0.06 | -3.61 | 0.00031 |
| **Count effects** |  |  |  |  |
| Classpose × log(counts<sub>train</sub>) | 0.16 | 0.01 | 17.32 | < 0.0001 |
| Semantic CP-SAM × log(counts<sub>train</sub>) | 0.17 | 0.01 | 18.43 | < 0.0001 |
| Stardist × log(counts<sub>train</sub>) | 0.16 | 0.01 | 17.78 | < 0.0001 |
| CellViT++ × log(counts<sub>train</sub>) | 0.10 | 0.01 | 11.05 | < 0.0001 |

We show that, on average and in terms of PQ, Classpose outperforms Semantic CP-SAM, Stardist, and CellViT++ by 0.08 (SE = 0.006), 0.14 (SE = 0.006) and 0.16 (SE = 0.006), respectively (p<0.0001 for a t-test comparing estimated marginal means for all comparisons; Figure 3A). The average differences between datasets revealed meaningful trends: performance in NuCLS was the worst, and in ConSep it was consistently better than that of other datasets (Figure 3B). Comparing cell type cluster performances indicated that PQ for both epithelial and immune cell types were similar and better than what was observed for other cell type clusters (Figure 1B). PQ in mesenchymal cell types — a relatively varied group encompassing muscle, stromal and connective tissue cells — was lower than for all other cell type clusters.

**Fig. 3.**
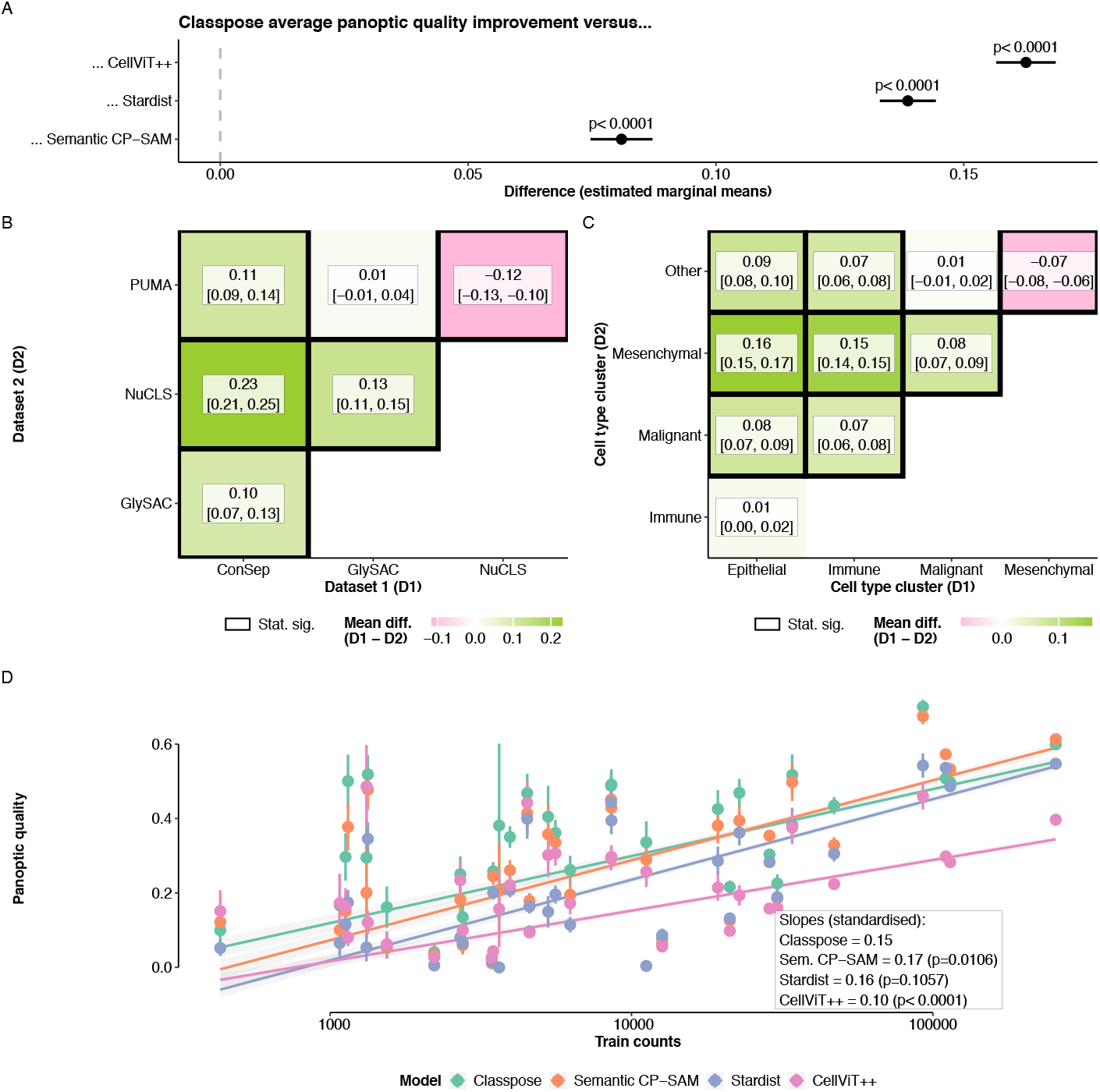
Multivariate analysis of factors underlying panoptic quality. **A**. Average difference in performance between Classpose and Semantic Cellpose-SAM (CP-SAM), Stardist and CellViT++ according to t-tests comparing the estimated marginal means. **B**. Average differences in panoptic quality between datasets according to t-tests comparing the estimated marginal means. A black box signifies whether specific dataset comparisons are statistically significant. **C**. Relationship between train proportions and the panoptic quality, stratified by model. p-values were calculated using Wald chi-squared tests comparing interaction term coefficients.

The model-decorrelated training proportion term coefficients were different from zero (Table 1), indicating that dataset-specific cell type proportions affected the performance (Figure 3C). Interestingly, this relationship was negative for CellViT++, indicating that, relative to the average, CellViT++ did not improve as much with class over-representation. Comparing the model-training count coefficients for Semantic CP-SAM, CellViT++, and Stardist with that of Classpose showed that Classpose and Stardist had comparable relationships between training proportions and PQ (p=0.15 for a Wald chi-squared test comparing coefficients). Increasing the training counts with Semantic CP-SAM led to larger PQ improvements while with CellViT++ this led to smaller performance improvements (p=0.011 and p<0.0001 for Wald chi-squared tests comparing coefficients, respectively).

We repeat these analyses for both DQ and SQ, showing that linear model coefficients follow similar patterns with some noteworthy differences (Table S5, Table S6, Figure S3, Figure S4). In DQ, Semantic CP-SAM performance does not improve more with an increase in cell counts when compared with Classpose. For SQ, i) only Stardist has a non-negative relationship between decorrelated training proportions and performance, and cell cluster performance is relatively similar, ii) there is no statistically significant difference between Classpose and CP-SAM and iii) performance for cell type clusters is similar (except for increased SQ in epithelial and other cell types when compared with mesenchymal cells).

When repeating these analyses with all datasets, including those used during pre-training (CoNIC, MoNuSAC), we including these datasets could lead to different conclusions, particularly for Classpose and CP-SAM (Figure S5, Figure S6, Figure S7, Figure S8). Both Semantic CP-SAM and Classpose would have inflated performance estimates when compared with those from CellViT++ for all metrics. When compared with Stardist, Classpose showed increased performance when not considering datasets used during pre-training, indicating that Stardist performed well on these datasets when compared to its average performance; Stardist performance also increases significantly in comparison with CellViT++ when including these datasets. Further, including datasets used during CP-SAM pretraining led to reduced differences between Classpose and Semantic CP-SAM, possibly indicating that Semantic CP-SAM benefited more from training/testing on data used during pre-training. This did not lead to Semantic CP-SAM equaling Classpose in terms of PQ or DQ.

To understand how Classpose performs in external datasets, we corresponded cell types between datasets and calculated Classpose and Semantic CP-SAM performance on these datasets after scaling test images to the training resolution. While PQ decreases when models are tested in external data (Figure 4A,B), this is cell type-specific. Inflammatory cells and lymphocytes are more easily segmented/detected across datasets. Macrophage, epithelial cells and connective/stroma cell external PQ, contrarily, were lower (Table S7). This is largely attributable to i) our broad definition of macrophages including both melanophages (macrophages which have engulfed melanin) and histiocytes (a broad category including not only macrophages but also dendritic cells) and the ii) the relatively large phenotypic variability of both epithelial and connective/stroma cells across tissues. When compared with Semantic CP-SAM, Classpose ranks first across most cell types (Figure 4C), with an estimated difference in PQ of 0.04 (p<0.0001 for a two-sided t-test comparing estimated marginal means).

**Fig. 4.**
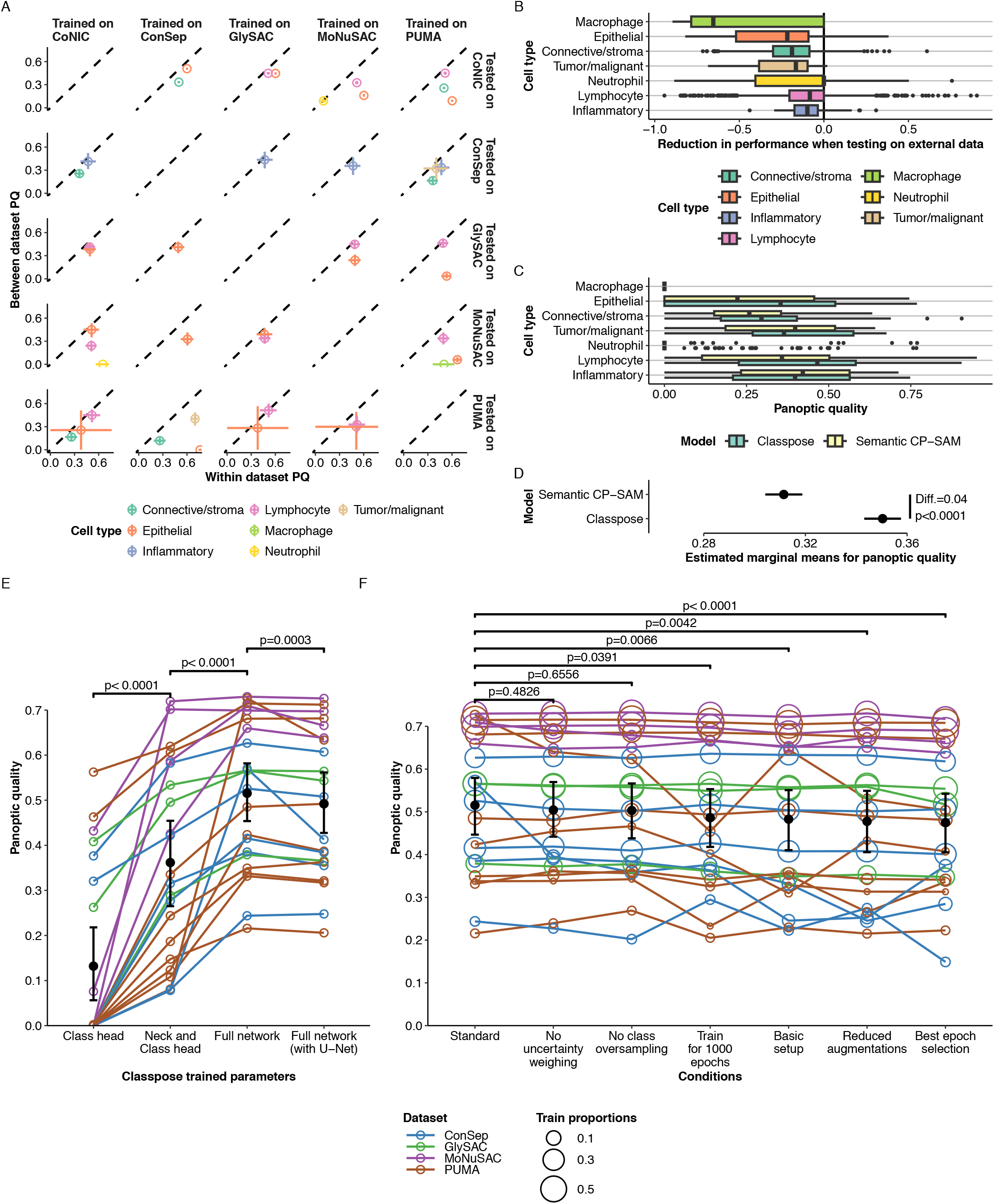
Panoptic quality estimates for external validation and ablation studies. **A**. External and internal panoptic quality comparison for different train/test dataset pairs and stratified by cell type. Each circle represents the average across all images and horizontal/vertical bars represent the 95% bootstrapped confidence interval. **B**. Distribution of image-specific differences between external and internal panoptic quality stratified by class type. **C**. Panoptic quality comparison between Classpose and Semantic CP-SAM. **D**. Estimated marginal means for external panoptic quality performance and the corresponding difference and associated p-value for a two-sided t-test comparing these estimated marginal means. **E**. Comparison of the effect of training different subsets of Classpose parameters on the panoptic quality. Each grey line and its corresponding hollowed points represents a class type. **F**. Effect on panoptic quality of training Classpose under different conditions via ablation experiments (Methods). For **E** and **F**, each line represents the same class across all training conditions. Both panels feature p-values obtained using twosided paired rank-sum tests. For both instances, points are coloured according to the dataset to which each class type belongs to, and the larger black points and vertical error bars represent the average and the bootstrapped 95% confidence interval.

### 2.3 Optimal performance through fine-tuning

The results above for Classpose were obtained by training the CP-SAM together with a classification head. To show that this is necessary for state-of-the-art performance, we additionally tested for performance differences when training i) the classification head alone, ii) the classification head and the last layer of the feature encoder (the “neck”), or iii) the entire Cellpose-SAM network and the classification head with an additional sub-network. We performed this for a mixture of older (ConSep, MoNuSAC) and newer smaller datasets (PUMA, GlySAC) made available only after the Cellpose-SAM preprint became available [9].

In Figure 4E, we show that training the entire network is necessary to achieve optimal performance (Table S8). In particular, the averaged PQ across all cell types and datasets improves from 0.134 (95%CI_boot_ = [0.061, 0.215]) to 0.362 (95%CI_boot_ = [0.280, 0.452]) when training both the class head and the neck (p<0.0001 for a two-sided paired rank-sum test). Further improvements are observed when training the full network, with the average PQ increasing to 0.515 (95%CI_boot_ = [0.451, 0.580], p<0.0001 for a two-sided paired rank-sum test). Adding the sub-network leads to a performance drop (PQ=0.492, 95%CI_boot_ = [0.426, 0.560], p=0.00036 for a two-sided paired rank-sum test).

Changes to other factors also influenced performance through ablation experiments (Figure 4F, Table S9). When compared with the average per-class PQ scores (0.515, 95%CI_boot_ = [0.451, 0.585]), removing either the uncertainty weighting or the class oversampling caused no significant performance drops (0.504, 95%CI_boot_ = [0.441, 0.562], p=0.48 and 0.503, 95%CI_boot_ = [0.440, 0.570], p=0.66). However, we observed performance drops when training for longer (1,000 instead of 500 epochs; 0.486, 95%CI_boot_ = [0.423, 0.554], p=0.039), using a basic setup (reduced augmentation panel, no class oversampling, no uncertainty weighting; 0.483, 95%CI_boot_ = [0.418, 0.551], p=0.0083), reduced augmentation panel (0.478, 95%CI_boot_ = [0.404, 0.546], p=0.0047), or training with a subset of data (20%) to select the best epoch during training (“Best epoch selection” in Figure 4F; 0.475, 95%CI_boot_ = [0.407, 0.545], p<0.0001). The effect observed for the basic setup is likely attributable to the reduced augmentation panel as the drop in performance was similar between both conditions.

We also tested whether using bf16 — a data type with wide hardware support and improved dynamic range over fp16 [22] — was comparable with fp32, used during training and regular inference. We show no performance drops when comparing the per-class metrics for Classpose (Figure S9) and improve inference times: 1.50 vs. 0.88 s/image for MoNuSAC, with n=85 test images with irregular sizes between 33 × 86 and 1771 × 1760 pixels, and 4.94 to 2.72 s/image for PUMA, with n=22 test images with 1024 × 1024 pixels. This constituted inference times up to 55% faster with no meaningful alterations in performance.

### 2.4 Classpose enables discovery at scale

We applied the CoNIC Classpose model to SurGen, a large cohort of colorectal cancer (CRC) WSIs. Classpose used 2 NVIDIA A6000 GPUs in parallel to predict 704,958,660 cells across 419 WSI, detecting large numbers of cells across 6 cell types Figure 5. This took, on average, 15:27 minutes per slide (Figure S10). Since eosinophils and neutrophils were rare in the calculated results (below 1% on average), downstream analyses were restricted to the four most abundant cell types: epithelial cells, lymphocytes, plasma cells, and connective cells. From these predictions we extracted morphological, spatial, and spatial-morphological features within pathologist annotated cancer regions.

**Fig. 5.**
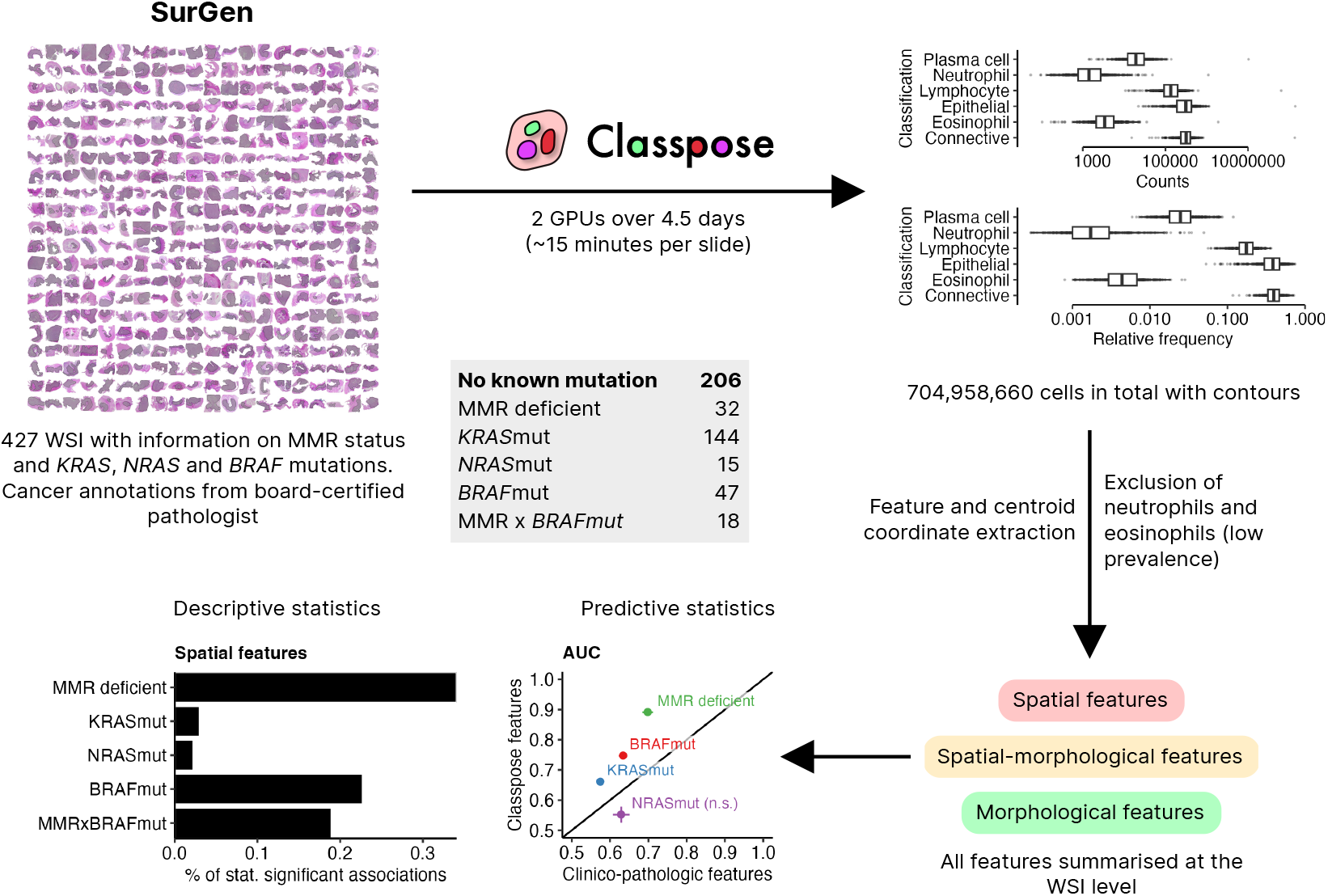
Classpose enables discovery at scale. Upon applying Classpose to SurGen, we extracted over 700,000,000 cells across 6 cell types. Upon exclusion of cell types whose cell prevalence was, on average, below 1% (neutrophils and eosinophils), we extracted spatial, spatial-morphological, and morphological features for each WSI. We used these features to predict different molecular conditions (MMR deficiency and mutations in *KRAS, BRAF*, and *NRAS*), and to characterise the spatial organization of cells.

We first determined associations between these features and five molecular conditions: MMR deficiency (dMMR); *KRAS*^MUT^, *BRAF* ^MUT^, and *NRAS*^MUT^; and the dMMR-*BRAF* ^MUT^ interaction (dMMRx*BRAF* ^MUT^ ) derived from the SurGen clinical and genomic annotations. The interaction term models the frequent *BRAF* ^MUT^/dMMR co-occurrence [23] and avoids confounding condition-specific effects. Most significant associations are accounted for by dMMR, *BRAF* ^MUT^, and their interaction, whereas *KRAS*^MUT^ and *NRAS*^MUT^ show a smaller set confined mainly to connective-cell morphological gradients (Figure 6, Figure S11; given the breadth of this analysis, we provide a more verbose version in section S10). Classpose-derived morphological and spatial features also improve dMMR, *KRAS*^MUT^, and *BRAF* ^MUT^ prediction beyond clinico-pathological variables alone, indicating that cell-level spatial phenotyping recovers biologically interpretable and clinically relevant signal from routine H&E.

**Fig. 6.**
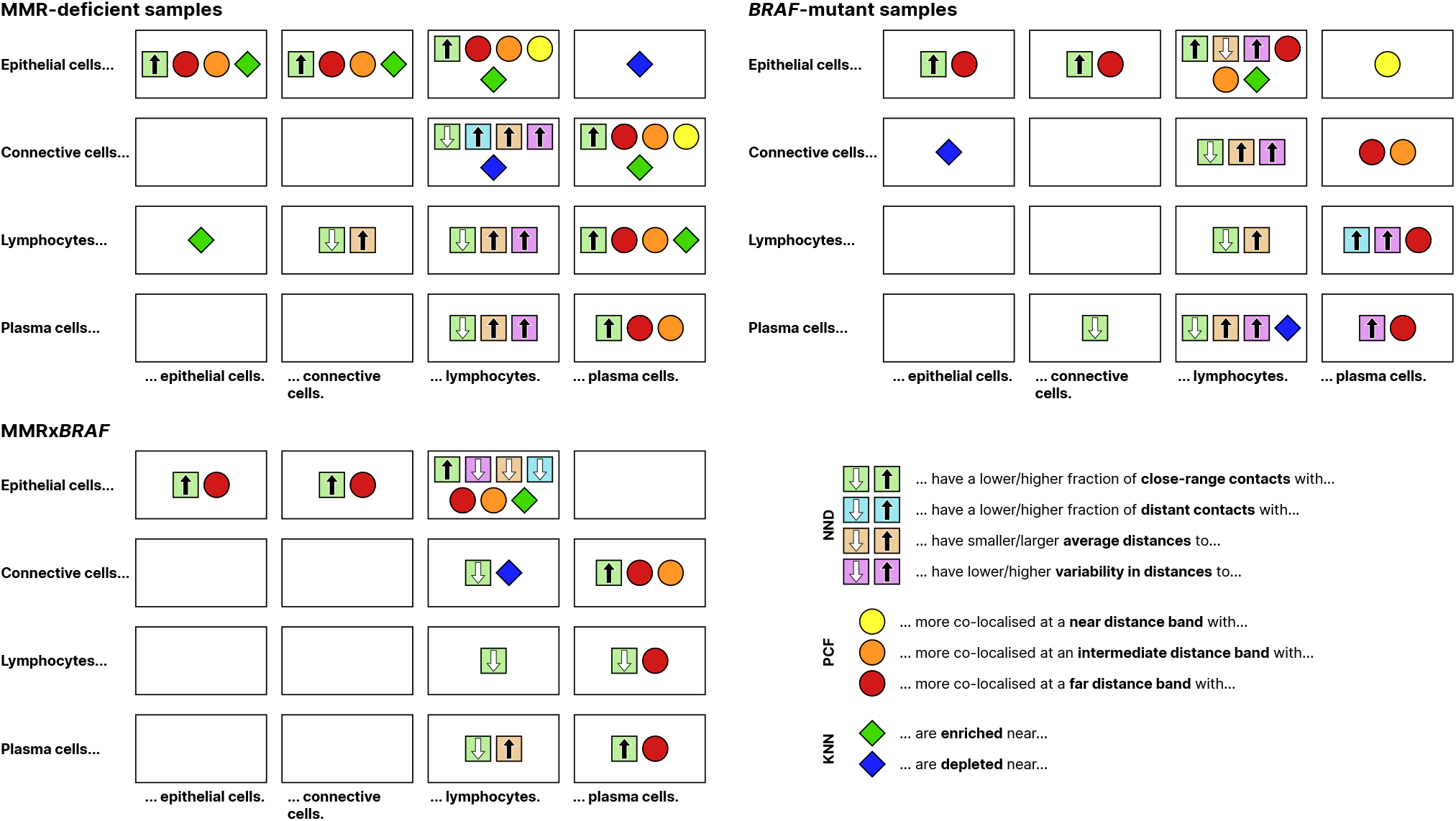
Spatial hallmarks driven by Classpose predictions. Summary of the associations between spatial features and the molecular conditions with the most frequent statistical associations: MMR-deficient samples, *BRAF* -mutant samples and samples which are simultaenously MMR deficient and *BRAF* mutant. Only statistically significant correlations are shown.

#### MMR deficiency: immune enrichment, connective dispersal, and elevated niche diversity

Through the quantification of spatial with Classpose, we resolve three quantitative spatial correlates characteristic of dMMR CRC (Figure 6).

Immune cells (lymphocytes, plasma cells) are found further from connective cells, but closer to epithelial cells. Given that H&E offers no way of discriminating lymphocyte subtypes, part of the signal detected using Classpose is likely attributable to T cells. Epithelial niche diversity — the Shannon entropy of the 10-nearest-neighbour label distribution — is correspondingly elevated, highlighting the overall increase in neighbourhood heterogeneity (Figure S13). In dMMR CRC, connective cells are further apart from one another and are closer to and more co-localised with epithelial cells (Figure S14, Figure S15).

Finally, we observe differential morphological gradients at the cancer border. In particular, the nuclear size (perimeter) of epithelial cells, plasma cells and lymphocytes decreases more steeply when approaching the cancer boundary in dMMR samples (Figure S12), while plasma-cell convexity (solidity) declines more slowly near the cancer border in dMMR samples.

#### *BRAF* ^MUT^: immune enrichment near cancer cells, connective-cell dispersal, and tighter epithelial–connective proximity

In our analysis, *BRAF* ^MUT^ CRC exhibits a highly diverse epithelial niche characterized by close spatial proximity between epithelial cells, immune cells (lymphocytes and plasma cells), and connective cells. While connective tissue cells are broadly dispersed from one another, they maintain an unusually tight association with the epithelial boundary. Because BRAF mutations frequently co-occur with mismatch repair deficiency (dMMR), many of these immune-enriched spatial features are shared between the two conditions and cannot be completely disentangled. However, the exceptionally close and uniform distance between epithelial and connective cells remains a distinct, independent spatial signature unique to the BRAF mutation itself, as it is absent in tumors with dMMR alone.

#### *KRAS*^MUT^ and *NRAS*^MUT^ are marked by few associations

*KRAS*^MUT^ samples exhibit predominantly differential connective-cell morphological gradients (Figure S12): as they approach the cancer border, the rate at which connective cell nuclei become more elongated, larger and less convex increases, indicative of longer, more irregular boundaries rather than simple enlargement. Additionally, a smaller fraction of epithelial cells lies close to connective cells in *KRAS*^MUT^ samples.

*NRAS*^MUT^ are uncommon in CRC (≈ 4%) [23] and show the fewest associations in our data. No significant associations were recovered in the spatial feature classes. The sole detectable signal is confined to morphological gradients at the border: connective cell nuclei have more elongated and less convex outlines near the boundary, mirroring the pattern seen in *KRAS*^MUT^, while plasma cell nuclei elongate at a higher rate. These observations should be treated as exploratory given the small number of *KRAS*^MUT^ samples (*n* = 15).

#### dMMRx*BRAF* ^MUT^ : distinct synergistic associations

The dMMRx*BRAF* ^MUT^ interaction comprises a large core of features shared with dMMR or *BRAF* ^MUT^ and a small set of significant changes.

*Shared features*. Given their known co-occurrence [23], it is expected that *dMMRxBRAF*^*MUT*^ cases share features from dMMR/*BRAF* ^MUT^ cases: intra-epithelial immune enrichment, elevated immune fraction, connective-cell dispersal, and lymphocyte perimeter gradients are all present in dMMRx*BRAF* ^MUT^ . Notably, the bulk immune-connective dispersal metrics (median and IQR of lymphocyte- and plasma cell-to-connective distances), significant in both parent conditions, do not reach significance in dMMRx*BRAF* ^MUT^, suggesting the interaction signal is concentrated at closer ranges rather than across the full distribution. Other features are inherited from dMMR (reduced connective cell self-enrichment, more lymphocytes and plasma cells near epithelial cells) or from *BRAF* ^MUT^ cases (epithelial cells closer to connective cells, with lower variability).

*Emergent features*. Only three features characterise dMMRx*BRAF* ^MUT^ but not parent conditions (Figure 6). The fraction of epithelial cells far from their nearest connective cell neighbour is reduced. Together with elevated closecontact fraction from both parent conditions, the reduced epithelial-to-connective distance distribution toward shorter ranges suggests increased epithelial-connective compartment intermingling. Second and third, niche diversity is independently elevated for lymphocytes and plasma cells, indicating that these cells occupy more heterogeneous local microenvironments.

#### Classpose-derived features improve prediction of MMR status and selected oncogenic mutations

Using the Classpose-derived features, we used reguralised logistic regression models to predict four different molecular conditions: dMMR and *KRAS*^MUT^, *BRAF* ^MUT^ and *NRAS*^MUT^. As shown in Figure 7A-D, we show that features derived from Classpose outperform the clinico-pathologic baseline for 3 molecular conditions: dMMR, *KRAS*^MUT^ and *BRAF* ^MUT^ (p=0.02, p=0.002 and p=0.01, respectively, for two-sided Mann-Whitney tests). For *NRAS*^MUT^ prediction, we show that using Classpose morphological features alone is worse than the baseline (p=0.02), while other Classpose models are comparable (p>0.05). Importantly, we note that using cellular density alone is insufficient to improve on baseline performance, highlighting that relative cellular prevalence is not sufficient for prediction. Furthermore, different feature sets are more predictive for specific tasks: while morphological features are especially important for dMMR prediction (p=0.02), both spatial and morphological features improve on the baseline in *BRAF* ^MUT^ prediction. However, improvement over the baseline for *KRAS*^MUT^ prediction (p=0.002) is observed only when all Classpose-enabled WSI features are combined. Using linear models allows us to easily estimate feature importance using standardised coefficients (Figure 7E-H; Figure S16). For instance, in dMMR and *BRAF* ^MUT^ prediction, it is evident that, among Classpose-enabled WSI features, variability in the epithelial nuclei perimeter (epithelial perimeter (std.)) drives predictive performance. For *KRAS*^MUT^, where only the combination of all WSI features outperforms the baseline, the WSI feature signature is considerably more complex: with morphological features quantifying variation driving prediction, and spatial features also contributing significantly towards this. Importantly, the factor contributing the most in absolute terms is the number of proximal contacts between epithelial and connective cells: this feature is negatively associated with *KRAS*^MUT^ samples, similar to what was shown above using descriptive statistics.

**Fig. 7.**
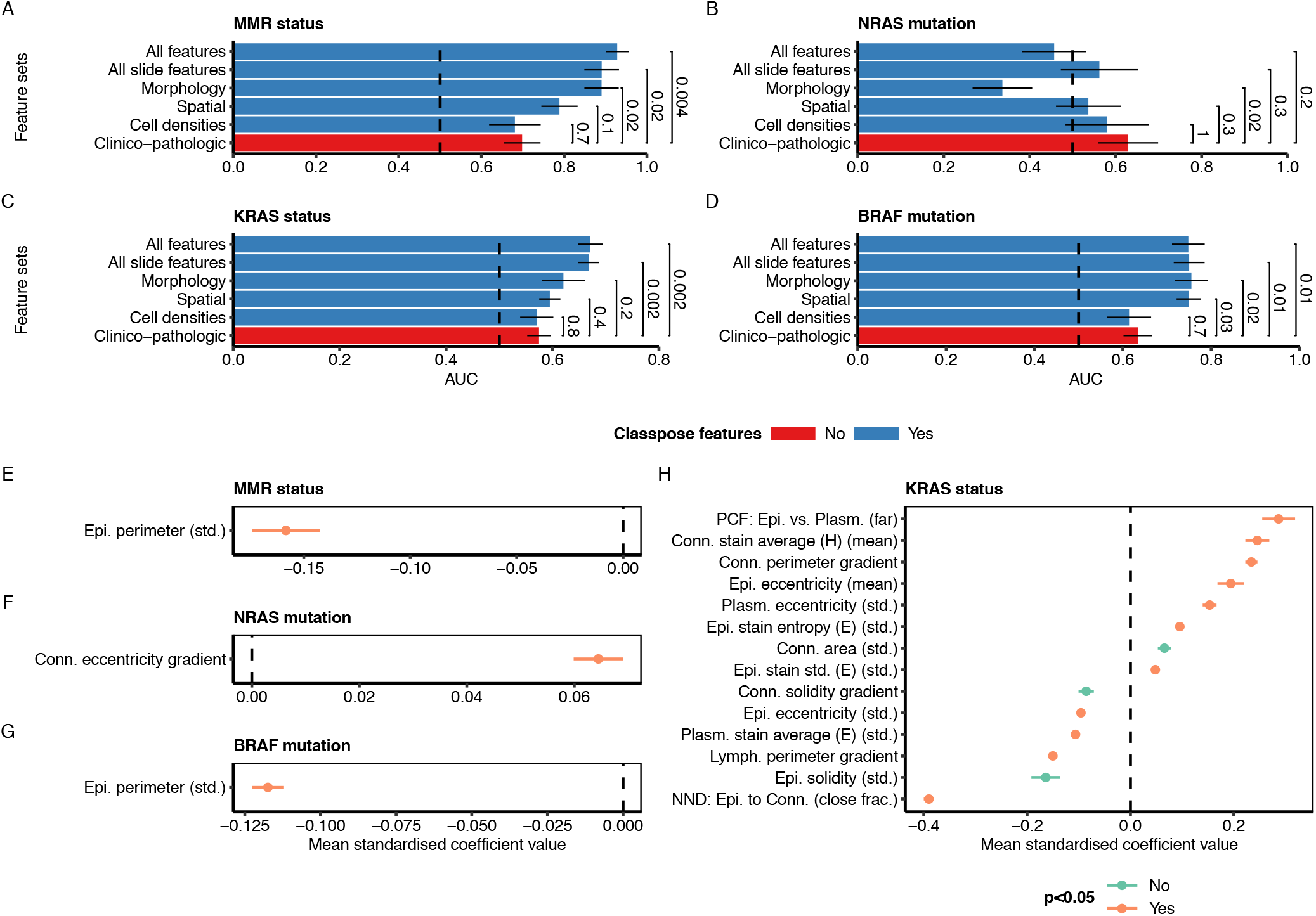
Classpose-derived features improve molecular condition prediction. **A-D**. AUC distribution for 10-fold cross-validation (CV) for different feature sets for the prediction of four distinct molecular conditions: MMR status and *KRAS, BRAF* and *NRAS* mutations. **E-H**. 10-fold CV feature importance (standardised coefficients) distribution for features where more than half of the folds had standardised coefficients different from zero for the model using all morphological features. For A-D, the vertical bars represent the average AUC while the horizontal bars represent the standard error around the mean, and the dashed vertical line represents prediction at random (AUC=0.5). Colours represent whether the models incorporate Classpose-enabled WSI features. For E-H, the points represent the mean while the horizontal bars represent the standard error around the mean. Colours represent whether the coefficient distribution is different from zero according to a two-sided Wilcoxon test.

## 3 Discussion

We introduced Classpose, a robust framework for cell phenotyping in H&E images built on Cellpose-SAM. Classpose consistently outperformed state-of-the-art baselines (Semantic CP-SAM, Stardist, CellViT++), demonstrating a statistically significant average increase of ≥ 8% in Panoptic Quality (PQ). Our architectural ablation studies revealed that restricting training to linear probing or high-level features (e.g., classification head/neck) yields sub-optimal results, confirming that full-network fine-tuning is required for complex cellular semantic segmentation [24]. To facilitate clinical translation, we integrated Classpose into a QuPath extension featuring automated tissue and artifact detection. Finally, we explored how Classpose-derived features can be used to discover novel biomarkers for molecular conditions using a large CRC WSI cohort.

To address potential data-leakage biases from datasets used during Cellpose-SAM pre-training (CoNIC and MoNuSAC), we focused our technical evaluation on unbiased datasets (ConSep, GlySAC, NuCLS, and PUMA); when analysing both the unbiased or all datasets, our findings show that Classpose outperforms other methods. The single exception was CoNIC, where Semantic CP-SAM performed best, likely attributable to the latter benifting the most from a training/testing data overlap. Across all models, systemic performance drops on NuCLS were attributed to its use of bounding boxes rather than true boundaries, validating that NuCLS benefits from a detection rather than segmentation objective [25].

On external validation, Classpose exhibited superior generalizability over Semantic CP-SAM, despite shared performance drops in highly variable cell lineages (macrophages, epithelial, and stromal cells). Grouping melanophages and histiocytes into a single “macrophage” category introduced expected complexity, as differentiating melanophages from tumor cells remains a known diagnostic challenge even for expert pathologists [26]. Furthermore, tissue-specific morphological plasticity in epithelial [27,28,29] and stromal cells explains cross-dataset drops, given that most training repositories originate from single-tissue sources.

Classpose-driven analysis of SurGen, a large CRC WSI cohort [30], revealed associations with different molecular conditions and recapitulated known associations which required more complex experimental setups. For dMMR samples, we show reduced lymphocyte-epithelial distances, recapitulating previous findings using immunohistochemistry and spatially resolved single-cell transcriptomics to characterise T cell infiltration in dMMR CRC [31,32,33], and a disruption in connective cell organisation, in line with previous findings [32]. Additionally, we show that epithelial cells are depleated near connective cells in *KRAS*^MUT^ samples, consistent with reports that *KRAS* signalling in MSS locally advanced rectal cancer cells modulate the activity of surrounding fibroblasts and depletes the extracellular matrix [34]. The discovery of cell type-resolved spatial signatures directly from routine H&E illustrates the potential of Classpose for hypothesis generation and biological discovery at WSI scale. While prior work has quantified immune-cell density as a function of the distance to the cancer border [35] and gradients in protein marker expression in CRC tissue [36], the measurement of morphological gradients has not been reported.

Finally, Classpose-derived features predict molecular conditions (dMMR, *KRAS*^MUT^, and *BRAF* ^MUT^) directly from cell type-specific spatial and morphological features and discovers novel morphological associations. While the prediction of these mutations from H&E is not novel [37,38], linking the predictive power with simple features is: for dMMR and *BRAF* ^MUT^, we show that epithelial anisonucleosis (i.e. perimeter variability) is essential for the prediction of either. We also show how *KRAS*^MUT^ prediction is driven by a complex spatial-morphological signature, in line with previous findings [38].

While extensive in its analysis, our work has limitations. While data splitting led to ROIs/WSIs overlaps, a more extensive external validation beyond cell type matching is required to further validate Classpose performance, particularly for datasets where data contamination may have benefited Classpose/Semantic Cellpose-SAM (CoNIC, MoNuSAC). Our discovery findings pertaining to lymphocytes or connective cells require further investigation to highlight which cellular subtypes drives morphological/spatial changes, particularly as “connective” is a broad morphological/functional category. Additionally, our predictive modelling predicts a single mutation, potentially shadowing the effects of epistasis, and the contributions from dMMR and *KRAS*^MUT^ cannot be fully disentangled. While we unconver emergent features for dMMRx*BRAF* ^MUT^, they can be reflective not of genuine epistatic effects, but of a statistical effect in which cases carrying both alterations are at the extreme end of distributions that do not individually reach significance in either parent condition alone. Resolving this would require experimental data beyond the scope of this study.

## 4 Methods

We start here by denoting two separate tasks in cell segmentation: **instance segmentation**, which aims to determine whether a pixel belongs to a unique cell, and **semantic segmentation**, which aims to determine to which cell type a pixel belongs to.

### 4.1 Data

We made use of six distinct datasets to train Classpose — **CoNIC** [39], **ConSep, NuCLS** [25], **MoNuSAC** [40], **GlySAC** [41] and **PUMA** [42]. **CoNIC** and **ConSep** are both derived from colorectal histopathology slides, and feature diverse and over-lapping sets of patches annotated using different pipelines and classes. While part of **ConSep** has been integrated into **CoNIC**, they differ in terms of resolution, annotated cell types and annotation protocol. **NuCLS**, on the other hand, was derived from breast cancer histopathology slides from The Cancer Genome Atlas (TCGA) project [43]. **MoNuSAC** contained organ segmentations from multiple organs for WSI derived from multiple regions and cancers in TCGA [43]. **GlySAC** and **PUMA** contained annotated nuclei instances for gastric cancer and melanoma, respectively.

All datasets without author-provided splits were split into training and testing images and organized into arrays containing images of multiple sizes. The number of cell type instances for all datasets is described in Table 2.

**Table 2.** Cell type composition for all datasets. Cases where cells not labeled (Unlabeled) were ignored during training and not considered during testing. Stars (“*”) indicate datasets which are used during Cellpose-SAM pre-training; for MoNuSAC this is only the case for its training data, while for ConSep this is only the case as part of it is included in CoNIC. Only a subset of CoNIC data is used for Cellpose-SAM pre-training (n=3,863 of images with at least one nuclei annotation). However the authors did not specify which images were used during pre-training.

| Dataset | Cell type index | Cell type | Train count | Test count |
| --- | --- | --- | --- | --- |
| **CoNIC\*** | 1 | Neutrophil | 4,587 | 495 |
|  | 2 | Epithelial | 255,684 | 26,398 |
|  | 3 | Lymphocyte | 110,295 | 10,638 |
|  | 4 | Plasma | 28,696 | 3,269 |
|  | 5 | Eosinophil | 3,475 | 374 |
|  | 6 | Connective | 114,101 | 11,849 |
| **ConSep** | 1 | Other | 371 | 561 |
|  | 2 | Inflammatory | 3,941 | 1,638 |
|  | 3 | Healthy epithelial | 1,117 | 503 |
|  | 4 | Malignant epithelial | 4,420 | 2,711 |
|  | 5 | Stroma | 4,805 | 2,869 |
|  | 6 | Muscle | 901 | 495 |
| **GlySAC** | 1 | Other | 3,386 | 3,121 |
|  | 2 | Lymphocyte | 7,409 | 4,672 |
|  | 3 | Epithelial | 7,154 | 5,133 |
| **MoNuSAC\*** | 1 | Epithelial | 14,537 | 6,470 |
|  | 2 | Lymphocyte | 15,654 | 6,751 |
|  | 3 | Macrophage | 587 | 260 |
|  | 4 | Neutrophil | 634 | 147 |
|  | - | Unlabeled | 0 | 154 |
| **NuCLS** | 1 | Tumor | 19,266 | 2,013 |
|  | 2 | Stroma | 8,120 | 1,007 |
|  | 3 | Lymphocyte | 12,463 | 1,100 |
|  | 4 | Plasma cell | 5,016 | 536 |
|  | 5 | Macrophage | 1,211 | 140 |
|  | 6 | Other | 908 | 83 |
|  | - | Unlabeled | 7,036 | 473 |
| **PUMA** | 1 | Apoptosis | 1,576 | 228 |
|  | 2 | Tumor | 51,722 | 5,511 |
|  | 3 | Endothelial | 1,495 | 201 |
|  | 4 | Stroma | 3,359 | 497 |
|  | 5 | Lymphocyte | 19,421 | 2,220 |
|  | 6 | Histiocyte | 6,292 | 875 |
|  | 7 | Epithelial | 2,052 | 159 |
|  | 8 | Melanophage | 601 | 94 |
|  | 9 | Other | 793 | 93 |

**CoNIC** ^7^ was split using region-of-interest (ROI) of origin stratification (i.e. ROIs were present in only the training or test split). The 4,981 patches of size 256 × 256 were used as provided by the dataset authors and split into training (90%) and testing (10%) sets (n=4,450 and n=531 for training and testing, respectively). 6 cell classes are available for CoNIC: “epithelial”, “lymphocyte”, “plasma”, “neutrophil”, “eosinophil”, “connective”.

**ConSep** was accessed through the Kaggle website ^8^. We used the split provided by the dataset authors in the Kaggle webpage: 27 1, 000 × 1, 000 pixel images for training and 14 1, 000 × 1, 000 pixel images for testing. 7 cell classes are available for ConSep: “other”, “inflammatory”, “healthy epithelial”, “dysplastic/malignant epithelial”, “fibroblast”, “muscle” and “endothelial”. We group “fibroblast” and “endothelial” as the latter has very few instances into a general “stroma” category. Unlike Horst and others [8], we do not group “healthy epithelial” and “dysplastic/malignant epithelial”.

**NuCLS** ^9^ was split at the slide level using their TCGA nomenclature. 1,744 images of variable pixel sizes were split into testing and training slides (n=1,589 and n=155 for training and testing, respectively). For compliance with training, training images smaller than 256 × 256 were padded to 256 × 256. For testing, images are kept whole and no patching is performed. NuCLS features 12 distinct classes: “Tumor cells”, “Fibroblast”, “Lymphocyte”, “Mitotic figure”, “Vascular endothelium”, “Plasma cell”, “Apoptotic”, “Macrophage”, “Neutrophil”, “Myoepithelial”, “Eosinophil”. Given that some of these annotations had very low abundance (“Mitotic figure”, “Vascular endothelium”, “Apoptotic”, “Neutrophil”, “Myoepithelial”, and “Eosinophil” have fewer than 1,000 instances each across nearly 60,000 instances), we merged “Mitotic figure” with “Tumor” into a generic “Tumor” classification, “Fibroblast” and “Vascular endothelium” into a generic “Stromal” classification, and merged the “Myoepithelium”, “Apoptotic body”, “Neutrophil”, “Ductal epithelium” and “Eosinophil” classes into a generic “Other cells” class. Finally, nearly 17,000 instances have no classification annotations, and a relatively large number of cells are annotated only as bounding boxes, making this dataset particularly challenging.

**MoNuSAC** ^10^ was used as provided by the original challenge organizers with 209 train and 85 test images. The 4 original cell classes (epithelial, lymphocyte, macrophage, neutrophil) were considered. In particular, we utilized the approach from [9] to convert annotations from XML to tiff format ^11^. In the testing data, over 2,000 instances have no classification annotations and are, as such, ignored for semantic evaluation purposes. Additionally, some images have regions annotated as “ambiguous” (i.e. there is no way of clearly determining the class of a given instance) and cells with an overlap with ambiguous regions were also ignored for semantic evaluation purposes.

**GlySAC** ^12^ was used with the three available cell classes (lymphocytes, epithelial (normal and tumor) and other cell types) provided by the authors of GlySAC using their notation and train/test splits [41]. GlySAC contained 59 images derived from 8 WSI; these were divided by the authors into training (n=34) and testing (n=25) sets. The heuristic categorization of each cell into a broad cell type (such as normal and tumor epithelial cells or different types of lymphocytes) was performed.

**PUMA** ^13^ contained 206 1024 × 1024 pixel ROIs (103 from primary sites and 103 from metastatic sites). We divided these ROIs into training (n=183) and testing (n=22) sets. PUMA was used after merging two cell types with relatively low proportions (plasma cells and neutrophils) into an “Other” cell type class. All other cells annotations — apoptotic, tumor, endothelium, epithelial, and stromal cells, lymphocytes, histiocytes and melanophages — were used as provided [42].

To display Classpose results in H&E whole slide images (WSI) in Figure 1 and Figure S1, we used a WSI from an in-house cohort of FFPE colorectal tissues, scanned using a Hamamatsu scanner at 40x magnification with resolution of 0.2261 microns per pixel.

#### 4.1.1 Train/test splits

Dataset splits were used if provided by the original dataset authors (**ConSep, MoNuSAC** and **GlySAC**). For all other cases, data was split at the ROI level (**CoNIC** and **PUMA**) or at the WSI level (**NuCLS**). This reduced the possibility of shortcut learning while also making our test considerably more robust. For **CoNIC, PUMA** and **NuCLS**, ROI/WSI splitting was performed greedily: after shuffling all ROI/WSI, each was iteratively assigned to either train or test based on the current train and test cell class counts at slide index *i* (*N*_train_ and *N*_test_, respectively). If the ratio between *N*_test_ and *N*_train_ + *N*_test_ contained more than two cell types where this ratio was smaller than the testing fraction (10%), the ROI/WSI was assigned to the testing set; otherwise the ROI/WSI was assigned to the training set. This process was repeated 250 times and the best split (the one whose training and testing proportions were the most similar) was kept.

### 4.2 Classpose: extending Cellpose for semantic segmentation

To extend Cellpose for semantic segmentation (i.e. to develop Classpose), we adapted their latest iteration (CPSAM) [9] to be easily adjustable to perform this task and make this public at https://github.com/sohmandal/classpose. We clustered these enhancements into three broad categories: **pre-processing, model architecture** and **training strategies**.

#### 4.2.1 Pre-processing

Biological datasets are frequently unbalanced, with some cell types being frequently under-represented in H&E data; this can lead to sub-par performance for rare cell types. To address this, we relied primarily on different data augmentations while also over-sampling under-represented cell types.

##### Augmentations

Data augmentation and preprocessing pipelines were parallelised to improve computational efficiency. In addition to the random rotation and resizing augmentations employed in Cellpose, we incorporated several augmentation strategies from other works [15], including H&E colour space perturbations, Gaussian noise injection, random blurring, and hue-brightness-saturation (HBS) adjustments. Furthermore, we introduced HaematoxylinEosin-DAB (HED) colour space augmentations [45], with a 50% probability (during training, either HED or H&E staining augmentations were picked at random). The augmentations are applied in a structured sequence: colour → quality → geometrical transformations. This comprehensive multi-step strategy provided robustness to complex variations in staining, illumination, and overall image quality.

##### Oversampling under-represented classes

To address class imbalance through oversampling, we first computed the relative frequency of each cell class *c* ∈ *C* in the dataset, noted as *F*_*c*_. Class weights were then calculated as *W*_*c*_ = 1*/F*_*c*_. For each image *k*, we calculated the count of cells belonging to class *c*, denoted *a*_*k,c*_. Finally, image-specific sampling weights were computed using Equation 1, where *τ* controls the degree of bias towards under-represented classes. These image-specific weights are then used to randomly sample *n* images for training at the beginning of each epoch.

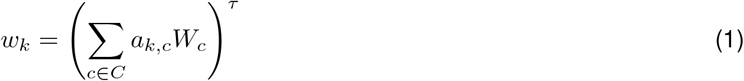

#### 4.2.2 Model architecture

While the original authors provide an example implementation of semantic segmentation, CP-SAM was developed to perform instance segmentation alone. We thus expand its architecture to simultaneously predict class labels for the segmented instances. Our model design choice resolves the target task in multi-task learning fashion i.e. learning both segmentation and classification at the same time. To this end, we detached instance from semantic segmentation. This allowed us to carry out ablation experiments such as fine-tuning only parts of Classpose (described below).

#### 4.2.3 Training strategies

##### Training loss design

Since Classpose learns both segmentation and classification tasks in parallel, we supervised the segmentation and classification heads simultaneously. For the segmentation task, we retained the same loss formulation as in CP-SAM: a combination of binary cross-entropy loss to classify between the foreground and background, and a mean squared error loss for predicting the flow field. To supervise the per-pixel class labelling, we combined multi-class cross-entropy and focal Tversky losses [14]. This combination leveraged their complementary strengths [46]: cross-entropy provides stable optimisation over predicted class probabilities across all pixels, while the focal Tversky loss emphasises spatial overlap and boundary accuracy. Further, we assigned class weights *w*_*c*_ to the classification losses by aggregating pixel-level class counts across all training labels and applying inverse frequency weighting normalised by the median class count with square root smoothing. This approach assigned higher weights to rarer classes and was inspired by StarDist’s CoNIC challenge solution [15]. The total loss per sample is defined as a weighted combination of the three component losses:

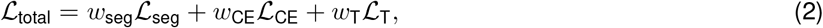

where ℒ_seg_ is the segmentation loss as defined in CP-SAM, ℒ_CE_ is the multi-class cross-entropy loss, and ℒ_T_ is the Tversky loss. The weights *w*_seg_, *w*_CE_, and *w*_T_ control the relative contribution of each loss term during training.

##### Uncertainty-based loss weighting for multi-task learning

In a multi-task learning setting like in Classpose, the individual loss terms often differ significantly in scale and convergence behaviour. Fixed scalar weighting of losses tend to be difficult to tune manually, which could also lead to suboptimal trade-offs between tasks. Therefore, to dynamically balance the contributions generated from the different loss components in 2, we adopted the uncertainty-aware weighting scheme proposed by [16], which treats each task’s loss as being corrupted by task-specific homoscedastic noise, where the model learns not only the task predictions but also the relative confidence (or uncertainty) for each task. As shown in 2, the contribution from each task-specific loss is scaled by the inverse of the variance (1*/σ*^2^) of a learnable parameter *σ*, which encodes the homoscedastic uncertainty associated with that task.

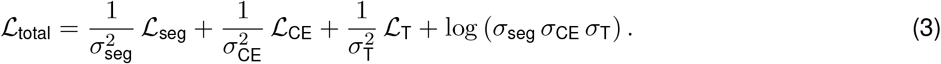

Here *σ*_seg_, *σ*_CE_, and *σ*_T_ are learnable parameters that represent the model’s homoscedastic uncertainty for each subtask. *log*(*σ*_seg_*σ*_CE_*σ*_T_) is a regularisation term used to penalise the model from becoming overconfident, which would otherwise dominate the overall training and suppress performance of other tasks. Throughout Classpose training *σ*_seg_, *σ*_CE_, and *σ*_T_ are learned end-to-end.

##### Sparsity-compliant training

To handle datasets with sparse semantic annotations, such as NuCLS where many cells have instance segmentation masks but lack class labels, we introduced a masking mechanism that excluded unlabelled pixels from the calculation of the classification losses. Let M denote the set of pixel indices that have valid semantic annotations. We defined **1**_M_(*i*) such that

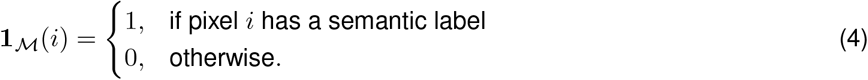

To make the classification losses sparsity-compliant, we multiplied all pixel-level terms in ℒ_CE_ and ℒ_T_ by 1_M_(*i*). This ensured that only pixels with valid semantic annotations contributed to the classification losses, while the segmentation loss ℒ_seg_ remained unaffected and continued to learn from all available instance masks.

### 4.3 Benchmarks

To better ground our performance analysis of Classpose, we compared it with three state-of-the-art methods: **StarDist** [47], **CellViT++** [8] and the CP-SAM adaptation for semantic classification provided by the authors (henceforth **Semantic CP-SAM**) [9]. These were compared with the standard fine-tuned Classpose.

For each datasets, we trained StarDist for 100 epochs and by following the exact model config as described in the conic-2022 branch ^14^ of StarDist’s Github repository. Thus, the number of rays used to describe a star-convex polygon was set to 64, the learning rate was 0.0003. We used patch sizes of 256×256 and a batch size of 4.

We trained CellViT++ linear classifier heads using their standard model fitting approach with frozen weights for three distinct backbones: the Virchow backbone [48], the SAM-H backbone from the original SAM preprint [11], and the HIPT backbone, one of the earliest foundation models for histopathology [49] as made available by the CellViT++ authors ^15^. These represent a considerable amount of the experiments performed in the original preprint, and all linear classifiers were trained on all datasets using a single validation fold. We ran 100 trials of Bayesian hyperparameter optimization and selected the best fold and backbone for each dataset based on the validation area under the receiver operating characteristic curve (AUROC). We optimize the number of neurons in the classifier (between 128, 256, and 512), the learning rate (between 0.00001 and 0.01), the weight decay (between 0.00001 and 0.01) and the learning rate schedule (between constant and exponential). Other parameters were used as described in the original CellViT++ publication: batch size of 256, dropout rate of 0.1 and *β*_1_ = 0.85 and *β*_2_ = 0.90 for the AdamW optimiser. Each model was trained for a maximum of 50 epochs with early stopping using a patience of 20 epochs. This process was performed for all six datasets using the cell types described in Table 2. To adjust CellViT++ to predictions across tiles with diverse sizes (as is the case in MoNuSAC or NuCLS), we implemented a simple tile-based prediction framework: whenever an image was larger than 256 × 256 (the size used to train our CellViT++ adaptations), the image was tiled into overlapping squares and individual cells and their respective coordinates were extracted. We then filtered out overlapping cells by systematically keeping the largest cell, thus preventing the selection of cells whose prediction was incomplete due to tiling artifacts.

Semantic CP-SAM and Classpose were trained for the same number of epochs: 500 epochs for all ConSep, GlySAC, MoNuSAC and PUMA, and 200 epochs for the two largest datasets, CoNIC and NuCLS. We adapted the training routine made available by the authors^16^ to allow us to use different datasets for training and validation. We also reduced the batch size from 16 to 4 due to GPU memory constraints.

### 4.4 Ablation experiments

#### Optimised parameters

To better understand to which extent transfer learning (i.e. training or retraining only a few layers), as opposed to full model finetuning, could lead to performance improvements, we considered four optimisation workflows: **i)** optimising only the semantic segmentation layer, **ii)** optimising the semantic segmentation layer, instance segmentation layer, and the SAM neck (which produces the features which are used directly by the semantic and instance segmentation layers), **iii)** optimising the entire network, and **iv)** optimising the entire network together with a an additional semantic classification sub-network (Supplimentary Methods). For clarity, the semantic classification layer is always optimised and initialised randomly, whereas the other layers may be frozen or trained but always initialised from CP-SAM weights. Across these experiments, we trained Classpose for 500 epochs and with a batch size of 4 on **ConSep, GlySAC, MoNuSAC** and **PUMA**. During each epoch, Classpose processes as many samples as are present in the entire training set. We selected these datasets as they are sufficiently small and in large part were not used during Cellpose-SAM pre-training.

#### Additional ablation experiments

We additionally assessed the impact of:

– **Having a stripped-down version of Classpose**. This experiment acted as the baseline for Classpose — in it, we excluded the uncertainty weighting and extensive augmentation strategy, and performed no oversampling of under-represented cell classes.
– **Increasing the number of training epochs**. We additionally tested how the number of epochs (500 vs. 1,000) affected the performance, allowing us to understand whether 500 epochs are sufficient and whether there might be detrimental effects associated with overfitting due to longer training times.
– **Using uncertainty weighting**. To estimate the impact of uncertainty weighting, we removed it from training runs.
– **Using a validation set**. To understand the usage of a validation set to monitor model training and select the best epoch (the one with the lowest validation loss), we used a subset of 20% of the training data for validation.
– **Using an extensive augmentation strategy**. We used a wide array of augmentations to artificially increase the sample size of our training data. To understand how our augmentation strategy might impact model training, we removed all augmentations except for HED augmentations.
– **Oversampling under-represented classes**. Finally, we performed under-represented class oversampling to guide training. To understand whether this might have a deleterious impact on training, we removed it.

All experiments were performed on the same four datasets (**ConSep, GlySAC, MoNuSAC** and **PUMA**) and over 500 epochs (excluding the epoch increase experiment, where models were trained over 1,000 epochs). Unless otherwise noted, the remaining training hyper-parameters (such as learning rate and weight decay) are identical to those from the default CP-SAM [9].

### 4.5 Validation and statistical analysis

As described we trained Classpose models for all datasets and compare them with three state-of-the-art methods. As evaluation metrics, we made use of three distinct per-class quantities:

– Average instance intersection over the union (IoU, or “segmentation quality”) — this metric is calculated as the mean Intersection over Union (IoU) across all cells in a given class. For a set of cells belonging to class *C*, 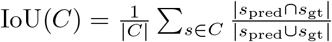, where *s*_gt_ represents the ground truth pixels for a given cell and *s*_pred_ represents the predicted pixels for that cell. To allocate different cell predictions to their corresponding ground truth cell, we considered a prediction to correspond to a ground truth whenever their IoU *> α*, where *α* is a pre-defined non-negative threshold. To avoid having multiple predictions for each ground truth cell, only the cell with the largest IoU is kept.
– F1-score at a given threshold (or “detection quality”) — the F1-score is calculated as 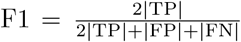, where |TP|, |FP| and |FN| correspond to the true positives, false positives, and false negatives, respectively. A ground truth instance is counted as a true positive if there exists a predicted instance with which it has IoU *> α*, otherwise, it is counted as a false negative. A predicted instance is labeled a false positive if it is not matched to any ground truth instance at the IoU threshold *α*. Although we used *α* = 0.5 for all datasets with mask-based ground truths, we set a lower IoU threshold of *α* = 0.25 for NuCLS, as a subset of its annotations are provided as bounding boxes, which typically yield lower IoU scores even for correctly detected instances.
– Panoptic quality — this metric [50] is the product of the average segmentation quality and the detection quality.

Prior to the calculation of per-class metrics, the binary cell instance map is multiplied by the cell class maps. Whenever cell classes were unavailable (i.e. cells were unlabeled but had an instance annotation) in the testing data, we excluded these cells and their corresponding predictions. To do so, we followed a simple heuristic: whenever a cell was unlabeled, any prediction which intersected with these cells was excluded.

Statistical analysis focused on comparing the different metrics of Classpose with those of Semantic CP-SAM, Stardist and CellViT++ while controlling for dataset, training proportions and training counts and cell type cluster for each cell type. To do so, and given that training counts and training proportions are co-linear, we start by calculating the residuals of p_train_ ∼ counts_train_, 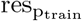, as the training count-uncorrelated effect of p_train_. Then, we fit a mixed-effects multivariate linear model (Equation 5) to the data. This linear model considers each image as an individual sample, and quantifies PQ, DQ and SQ for each class type. Missing values (if no predicted or ground truth instances of a given cell type are present) are excluded. Given that the entire CoNIC dataset was used during CP-SAM pre-training, we excluded it to avoid data leakage. We further excluded MoNuSAC as its training set was used during CP-SAM pre-training, giving both Classpose and Semantic CP-SAM an unfair advantage over other methods. We nonetheless fitted an additional mixed-effects multivariate linear model including both CoNIC and MoNuSAC to understand how our assessments would change in the presence of data leakage. Given that we used class metrics for individual images as samples and that this may bias results towards datasets with more images, we weighed each sample by its relative proportion within the dataset such that *w*_*i,c,D*_ = (*n*_*i,c,D*_ + 1)*/N*_*D*_. Here, *w*_*i,c,D*_ and *n*_*i,c,D*_ are the weight and counts for cells of class *c* on image *i* of dataset *D*, respectively, and *N*_*D*_ is the total number of cells in dataset *D*.

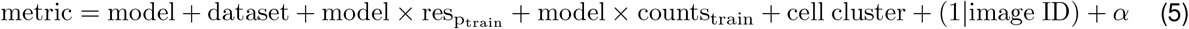

For Equation 5:

– metric is the relevant metric (while we focus on PQ, we perform the same assessments for DQ and SQ);
– dataset is a factor representing each dataset (CoNIC, ConSep, GlySAC, MoNuSAC, NuCLS and PUMA);
– model is a factor representing each model (Classpose, CP-SAM Semantic, StarDist and CellViT++). To reduce the number of models we focused on the CellViT++ model with the best average performance across all datasets;
– model × 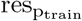 and model × p_train_ are the interaction term between the model and 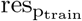 and p_train_, respectively;
– cell cluster is a loosely defined set of cell function clusters: **Immune** (Neutrophil, Eosinophil, Lymphocyte, Macrophage, Histiocyte, Melanophage, Plasma, Plasma cells, Inflammatory), **epithelial** (Epithelial, Healthy epithelial), **Mesenchymal** (Stroma, Connective, Endothelial, Muscle), **Malignant** (Tumor, Malignant epithelial), and **Other** (Other, Apoptosis). This allowed us to identify which broadly defined cell types are more easily identifiable in segmentation tasks;
– (1 | image ID) is a random effect accounting for image identifier;
– *α* is an intercept term.

To compare models, datasets and cell clusters we used the expected marginal means framework (also known as least-square means) using the emmeans package [51] for the R programming language. This framework allowed us to estimate the expected differences between different factors within the same categorical variable and their standard errors while controlling for the remaining variables using mixed effects linear models. The estimated difference *d* was then tested against the null hypothesis (*d* = 0) using a standard t-test, and Tukey p-value correction was applied to the p-values. Comparisons of training size-model interaction terms were performed using Wald chi-squared tests implemented via the linearHypothesis function (R package car) which evaluates whether interaction term coefficients are equal. We considered the interaction between training proportions and model to assess how increasing the availability of under-represented cell types affected the performance of each model to better understand how scaling dataset sizes could improve performance.

Finally, we performed cross-dataset comparisons by i) finding correspondences between cells types across datasets and ii) scaling test images to match the resolution of the training images for each model before prediction (Table S1). We excluded NuCLS from these data as masks in this dataset are a mixture of bounding boxes and detailed cell boundaries, making the evaluation more complicated. The complete mapping used for each dataset pair is available in the Supplementary Methods, but the broad rules are as follows:

– **Immune/inflammatory cells** — whenever the model was trained using discrete immune cell types (neutrophils, eosinophils, etc.) as in CoNIC or MoNuSAC and the test data had a single inflammatory/immune cell class (i.e. ConSep), the immune cell type predictions were aggregated into a single immune/inflammatory cell type;
– **Epithelial** — whenever the model was trained to predict more than one epithelial cell type (i.e. ConSep with healthy and malignant epithelial cell types) and the testing data had a single “epithelial” cell type, the predicted epithelial cell types were aggregated into a single “epithelial” cell type;
– **Macrophages** — whenever the model was trained to predict more than one type of macrophage (i.e. PUMA with melanophages and histiocytes) and the testing data had a single “macrophage” cell type, the predicted macrophage cell types were aggregated into a single “macrophage” cell type;
– **Connective/stroma** — the “connective” and the “stroma” cell types were used interchangeably;
– **Tumor** — the “tumor” and the “malignant epithelial” cell types were used interchangeably;
– **Background** — all cell types which did not have a clear match or belonged to “other” or “ambiguous” cell types were considered background and ignored for metric calculation purposes.

For this, we compared the external panoptic quality performance of Classpose with that of Semantic CP-SAM with a linear mixed effects model similar to that introduced above in Equation 5 and described in Equation 6. Here, we controlled only for train and test dataset and for cell class, with a random effect for image identifier. We also used sample weights proportional to the number of images of each class in each image. Our downstream analysis focused exclusively on comparing both methods.

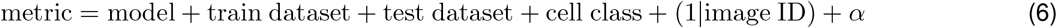

### 4.6 Whole slide image inference

To facilitate the use of Classpose across the wider community, we developed a Python-based entrypoint that performs inference on whole slide images (WSI) and produces several interoperable outputs. In addition to two QuPath-compatible GeoJSON files: containing cell contours (enabling cell composition and morphometric analyses) and cell centroids (supporting spatial analysis), Classpose also outputs a CSV file with cellular density statistics. This CSV reports, for each combination of region and cell class, the cell count and density in cells per mm^2^. Regions correspond to user-provided regions-of-interest (ROIs), such as tumor or stroma, otherwise, a single default region (tissue) represents the entire tissue area. Densities are normalised by the effective tissue area, with area converted to mm^2^ using the slide’s microns-per-pixel (MPP) values. Furthermore, Classpose generates a unified SpatialData [52] Zarr store that consolidates all tabular and spatial results. This includes vectorised shapes (cell, tissue, artefact, and ROI contours), point annotations (cell centroids with class labels), a table of cellular densities by region and cell type, and rich metadata (such as the slide’s MPP and Classpose model configuration). Together, these outputs support flexible downstream analysis, visualisation, and integration with established spatial biology analysis toolboxes, including MuSpAn [53] and Squidpy [54]. The workflow is as follows:

1. GrandQC, a tool for tissue and artefact detection [21], is used to detect all tissue and artefacts. The tissue annotations are used to sample tiles which contain any amount of tissue, while the artefacts are exported for user inspection.
2. A slide tiler and reader loads tiles into memory sequentially after determining whether or not they overlap with the tissue contours.
3. The inference pipeline runs on each tile using Cellpose functions which automatically tile images to a compliant size. This was adapted to enable multi-GPU inference using torch.multiprocessing. After each prediction (cell instances and cell type predictions), a polygon characterizing each cell type is extracted, together with the coordinates for its centroid, class and area.
4. To eliminate duplicates associated with overlaps, we developed a simple duplicate removal algorithm which constructs a k-d tree, ii) detects cells whose centroids are within 7.5 pixels of one another (i.e. cells with duplicates) and iii) removes all cells in a given duplicate group except the one with the largest area.

### 4.7 Discovery experiment

#### 4.7.1 Dataset and data processing

To demonstrate the utility of Classpose outputs, we utilised SR386 from the SurGen dataset [30]. SR386 contains 427 WSI obtained from colorectal cancer patients and has information on mismatch repair (dMMR) status and on *KRAS*^MUT^, *BRAF* ^MUT^, and *NRAS*^MUT^, as well as age, sex, staging, differentiation and whether tumours were mucinous. We codified MMR status and mutations as binary variables (0=absent, 1=present), staging and differentiation as a categorical variables (I/II/III/IV and Poor/Moderate (Mid)/Well, respectively), age as a continuous variable and sex as a binary variable (male/female). We first converted WSI data from CZI format to TIFF using VIPS [55] and BioFormats [56]. We then ran the Classpose model trained on CoNIC for all WSI across two NVIDIA RTX A6000 GPU cards. Regions of invasive colorectal cancer were annotated by a board-certified pathologist (KB), which we define as regions-of-interst (ROI). Adjacent colorectal mucosa and any grade of dysplasia were excluded from annotation. Non-tumour tissues (fat, smooth muscle, necrosis) were excluded wherever possible. Technical artefacts were also excluded when not infiltrated by tumour. We then calculated a set of morphological, spatial, and spatial-morphological features at the ROI level for each sample.

#### 4.7.2 Morphological features

We made use of a relatively concise panel of morphological features, namely: area, perimeter, convex hull area, and the major/minor axes length. We also calculated three derived features — eccentricity (as the ratio between major and minor axes), solidity (ratio between area and convex hull area), and form factor 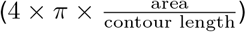. On top of these, we quantified colourimetric changes by quantifying the average, standard deviation, and statistical entropy (assuming 256 bins) of the RGB channels and of the H&E channels (using RGB-to-HED deconvolution method described in [57]).

We then summarised all features at the sample and cell type levels using the mean and standard deviation.

#### 4.7.3 Spatial features

We computed three spatial feature families that capture different but complementary aspects of cell-type organisation-the pair correlation function (PCF) characterises whether two cell populations are spatially associated across a range of distances, the nearest-neighbour distance (NND) quantifies how close each individual cell gets to its nearest neighbour of another type, and *k*-nearest-neighbour (*k*-NN) neighbourhood composition describes the mixture of cell types immediately surrounding each cell. Together they distinguish between population-level spatial patterning, pairwise proximity at the individual cell level, and local microenvironmental composition. All three were computed within each connected component of the annotated region and aggregated to the sample level by weighting each component by its area (see subsection S9.5 for full mathematical definitions). The PCF is a second-order summary statistic that quantifies whether cells of one type co-localise with, or avoid, cells of another type relative to complete spatial randomness (CSR), as a function of distance. We evaluated the PCF over annuli of width 0.010 mm to a maximum distance of 0.100 mm, and summarised each cell-type pair by the signed area under the PCF curve (relative to the CSR baseline of 1) in three pre-specified distance bands: short (0–0.025 mm), mid (0.025–0.050 mm), and long (0.050–0.100 mm). Positive values indicate clustering above CSR, while negative values indicate exclusion. The NND captures the most immediate contact scale. For each focal cell of type *A*, we measured the distance to its nearest neighbour of type *B*, and summarised the resulting distribution by its median, interquartile range, the fraction of focal cells within 0.020 mm of a *B* cell (close-contact proxy), and the fraction beyond 0.100 mm (exclusion proxy). *k*-NN neighbourhood composition characterises the local microenvironment of each cell using its *k* = 10 nearest neighbours. For each focal cell type *A*, we computed the mean fraction of *B* neighbours across all *A* cells, standardised against the global prevalence of *B* to calculated an enrichment score. We further computed the Shannon entropy of the 10-neighbour label distribution as a summary of niche diversity, and separately quantified the immune fraction (lymphocytes, plasma cells, eosinophils, and neutrophils) in the neighbourhood of each epithelial cell.

#### 4.7.4 Spatial-morphological features

To quantify spatial variations of feature values, we developed a statistical framework to characterise how features change as they approach the ROI border: **pericancerous morphological gradients**.

##### Pericancerous morphological gradients

We quantify the rate of change of four features (perimeter, solidity, eccentricity, and haematoxylin entropy) as they approach the cancer ROI border. To do so, we fit, for each WSI and for each feature, the linear model

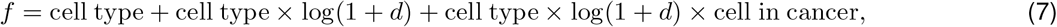

where *f* is the standardised feature value, *d* is the distance to the tumour border, cell type is a factor for each cell type, cell type × log(1 + *d*) is the cell type-specific rate of change for the entire WSI, and cell type × log(1 + *d*) × cell in cancer is the incremental cell type-specific rate of change which happens specifically within the cancer ROI. For downstream analysis, we focus on the last term and call it “pericancerous morphological gradients”.

#### 4.7.5 Statistical analysis

##### Descriptive statistics using spatial features

To determine whether there are statistically significant changes between molecular conditions and different spatial features, we make use of a general independence test using the permutation-based conditional inference framework implemented in the coin package for R [58]. We chose this to provide a robust test of the null hypothesis of independence without assuming a specific distribution for the underlying data. For cases where the omnibus test was significant (*p <* 0.05), post-hoc comparisons were performed by extracting the standardized Pearson residuals (Z-statistics) and using that to calculate p-values as 2 × *P* (*Z >* |*z*| ) (this ensures consistent numeric precision, essential for downstream multiple testing correction).

We assume that the relationship between spatial features and molecular conditions is of the form *f* = MMR + *KRAS* + *NRAS* + *BRAF* + MMR × *BRAF*, where each of the four independent variables is a binary variable accounting for the presence of specific mutations while MMR × *BRAF* accounts for the frequent dMMR/*BRAF* ^MUT^ co-occurrence [59]. We apply this to all spatial and spatial-morphological features, accounting for a total of 132 omnibus tests (at most, 660 post-hoc tests). We correct for multiple testing using the Benjamini-Hochberg methods.

##### Predictive statistics

To determine whether it is possible to predict different molecular conditions (dMMR, *KRAS*^MUT^, *NRAS*^MUT^ and *BRAF* ^MUT^) using features extracted from Classpose predictions, we make use of elastic net regularised classification models [60]. We use a panel of human expert-derived clinico-pathologic features (age, sex, stage, differentiation, whether the tumour is mucinous) as a baseline, and compare different models to this baseline: a model using only cell densities (I), a model using only spatial and spatial-morphological features (II), a model using only morphological features (III), a model using all WSI-derived features (I + II + III), and a final model which combines all WSI-derived features with the clinico-pathologic baseline features.

For each model we calculate the AUC distribution using nested cross-validation with 10 outer folds and 10 inner folds. The inner folds are used to determine the ideal value of *λ* (regularisation parameter), and the outer fold is used to determine the validation AUC. We compare the AUC distribution of each model with that of the baseline model using two-sided pairwise rank-sum tests.

## 4.8 Code and code availability

Classpose was developed using Python version 3.13 and PyTorch version 2.7.0. It is available at https://github.com/sohmandal/classpose. Adaptations introduced to CellViT++ are available and described in https://github.com/josegcpa/CellViT-plus-plus. Plotting and statistical analysis were performed using the R programming language [61] version 4.2. Model weights for each dataset are hosted in Huggingface in https://huggingface.co/classpose/classpose.

## 5 Acknowledgements

This work used the DiRAC Extreme Scaling service (Tursa) which is part of the STFC DiRAC HPC Facility (www.dirac.ac.uk). The service is hosted by the Edinburgh Parallel Computing Centre at the University of Edinburgh. Access to DiRAC resources was granted through a Director’s Discretionary Time allocation in 2024/25. We also thank Mariana Monteiro for valuable discussion.

## 6 Ethics declarations

TAG is named as a coinventor on patent applications that describe a method for TCR sequencing (GB2305655.9), a method to measure evolutionary dynamics in cancers using DNA methylation (GB2317139.0), and a method to infer drug resistance mechanisms from barcoding data (GB2501439.0). TAG has received honorarium from Genentech and consultancy fees from DAiNA therapeutics. TAG acknowledges funding from Cancer Research UK (DRCNPGMay21_100001) and the CRUK Convergence Science Centre (CTRQQR-2021\100009). KB was funded by the Swiss National Science Foundation (P500PM_217647/1).

## S7 Supplementary Figures

**Fig. S1.**
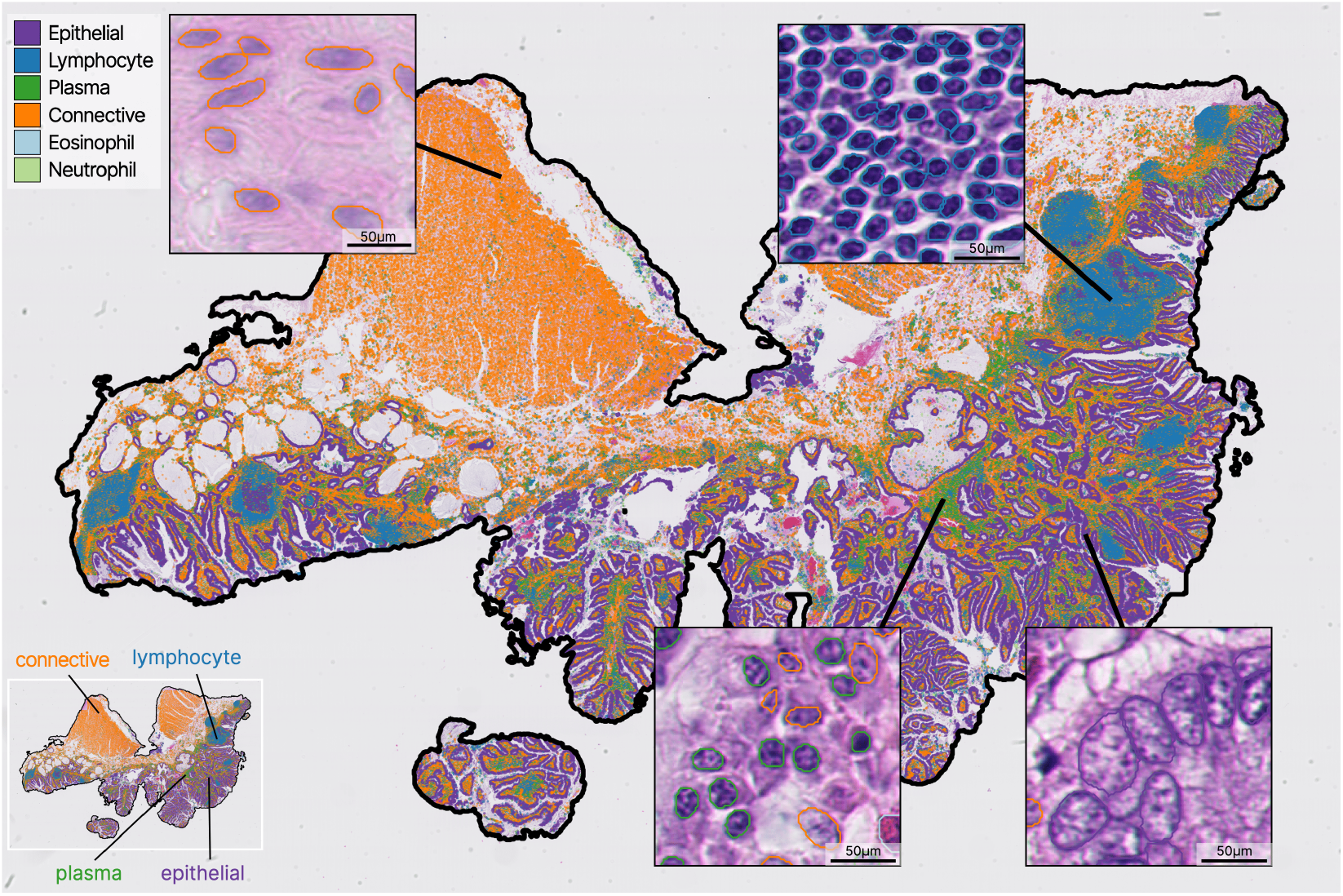
Classpose results on a WSI. Classpose results for a FFPE H&E human colorectal tissue sample from an in-house cohort, which has been scanned with a Hamamatsu scanner at 40x magnification. From Classpose’s result, we can easily detect the presence of different microenvironments throughout the WSI.

**Fig. S2.**
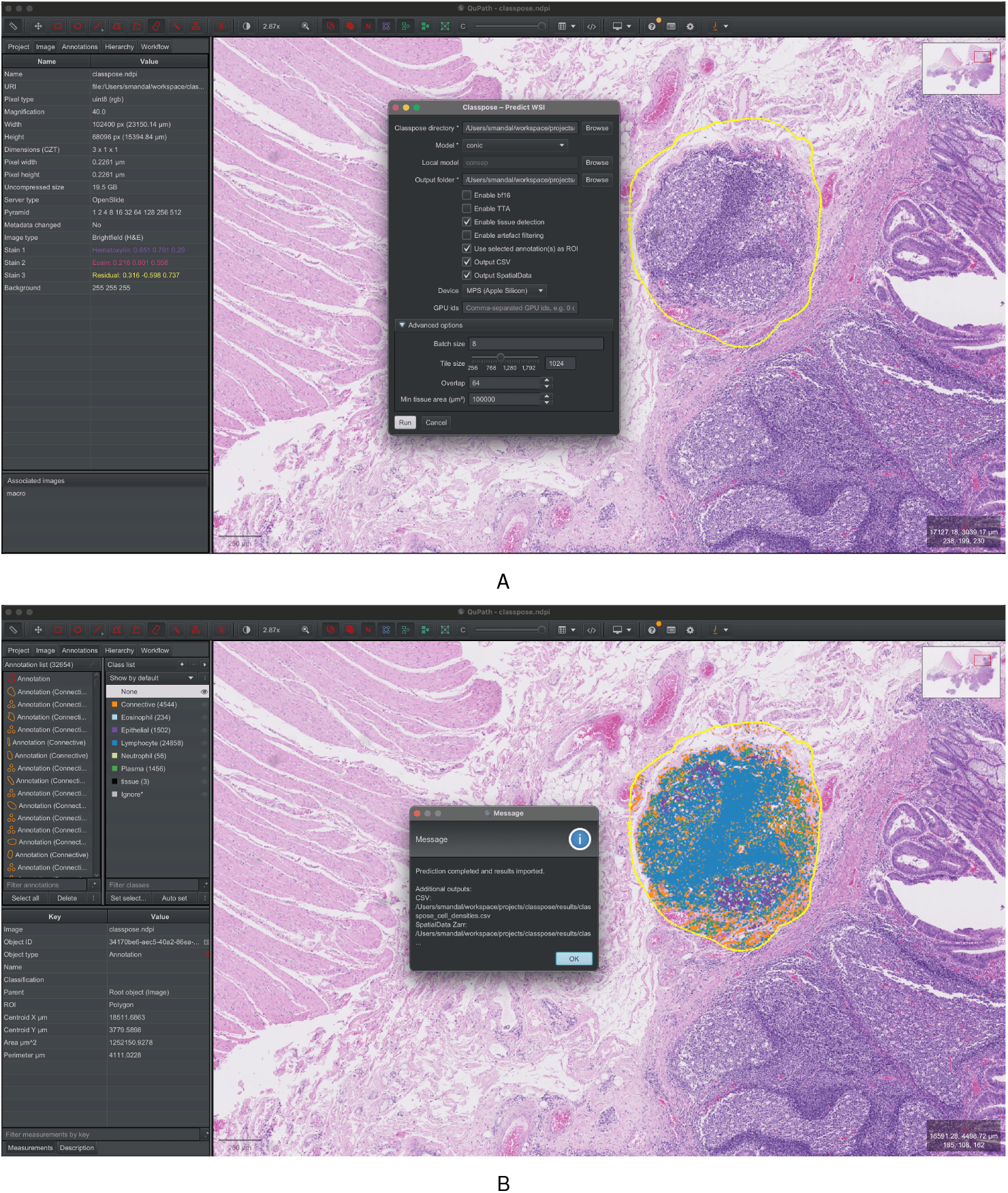
User interface and workflow of the Classpose QuPath extension. **A**. The ‘Classpose – Predict WSI’ dialog box is shown overlaid on a whole slide image (WSI) within QuPath. Key parameters are visible, including model selection, output folder, device configuration, and advanced options like tile size and batch size. A user-defined region of interest (ROI), outlined in yellow, is selected for targeted inference. **B**. The post-inference state, displaying the results imported into QuPath. The predicted cell classifications are visualised as colored dots within the ROI, with the corresponding class list (e.g., Lymphocyte, Epithelial, Plasma) shown in the sidebar. A message box confirms successful prediction and lists the generated output files (CSV, SpatialData).

**Fig. S3.**
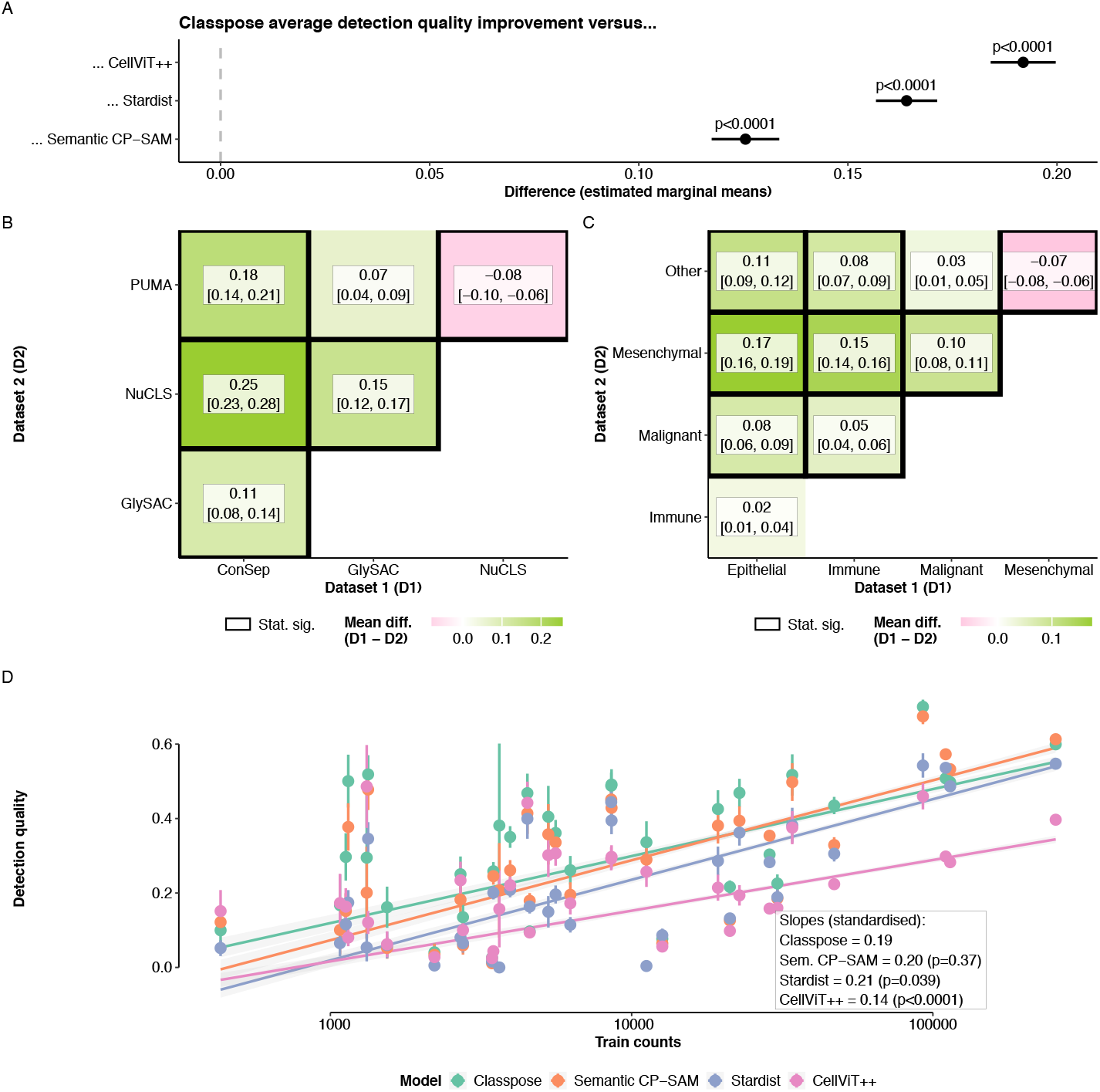
Multivariate analysis of factors underlying detection quality. **A**. Average difference in performance between Classpose and Semantic Cellpose-SAM, Stardist and CellViT++ according to t-tests comparing the estimated marginal means. **B**. Average differences in detection quality between datasets according to t-tests comparing the estimated marginal means. A black box signifies whether specific dataset comparisons are statistically significant. **C**. Relationship between train proportions and the detection quality, stratified by model. p-values were calculated using Wald chi-squared tests comparing interaction term coefficients.

**Fig. S4.**
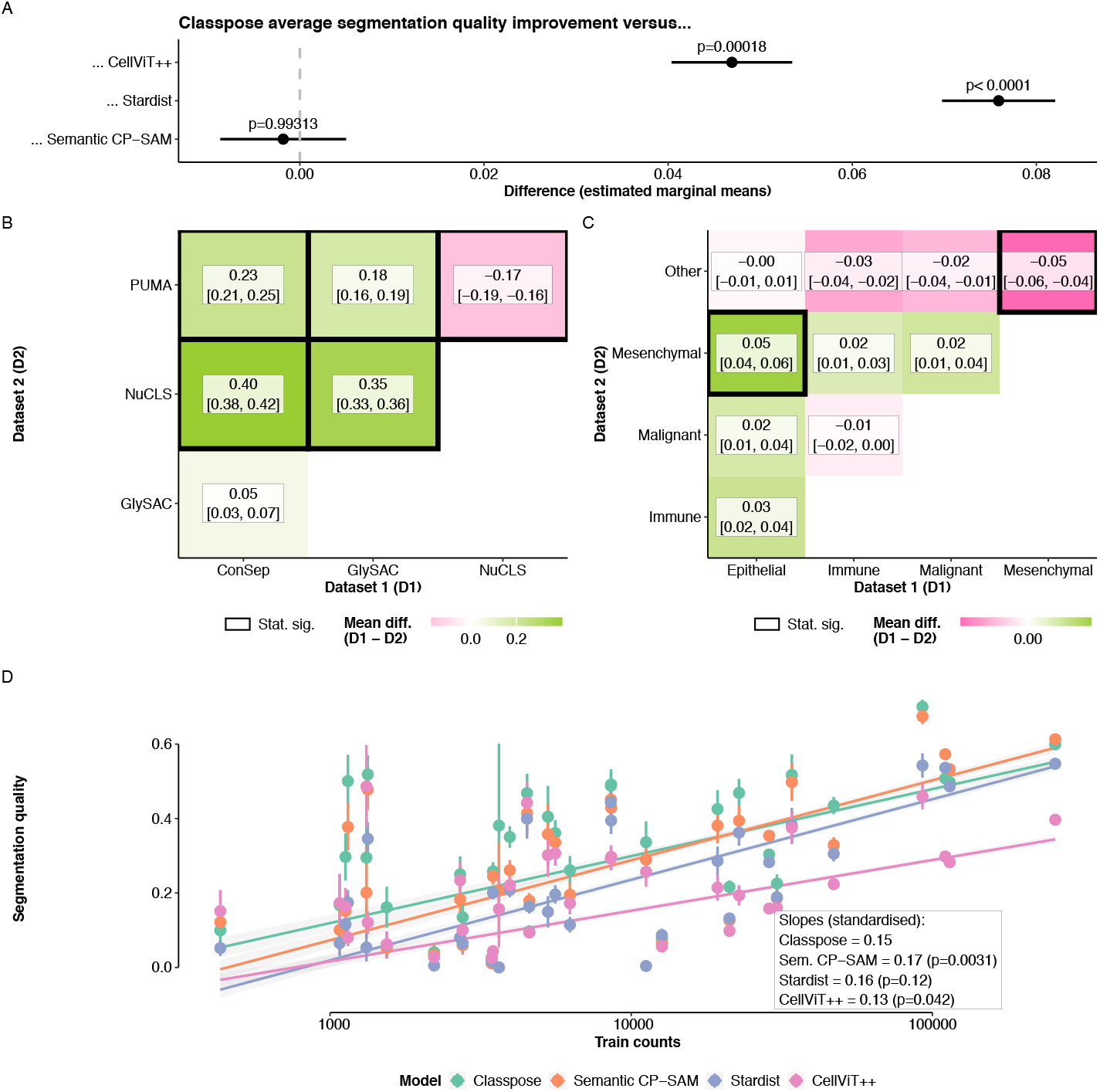
Multivariate analysis of factors underlying segmentation quality. **A**. Average difference in performance between Classpose and Semantic Cellpose-SAM, Stardist and CellViT++ according to t-tests comparing the estimated marginal means. **B**. Average differences in segmentation quality between datasets according to t-tests comparing the estimated marginal means. A black box signifies whether specific dataset comparisons are statistically significant. **C**. Relationship between train proportions and the segmentation quality, stratified by model. p-values were calculated using Wald chi-squared tests comparing interaction term coefficients.

**Fig. S5.**
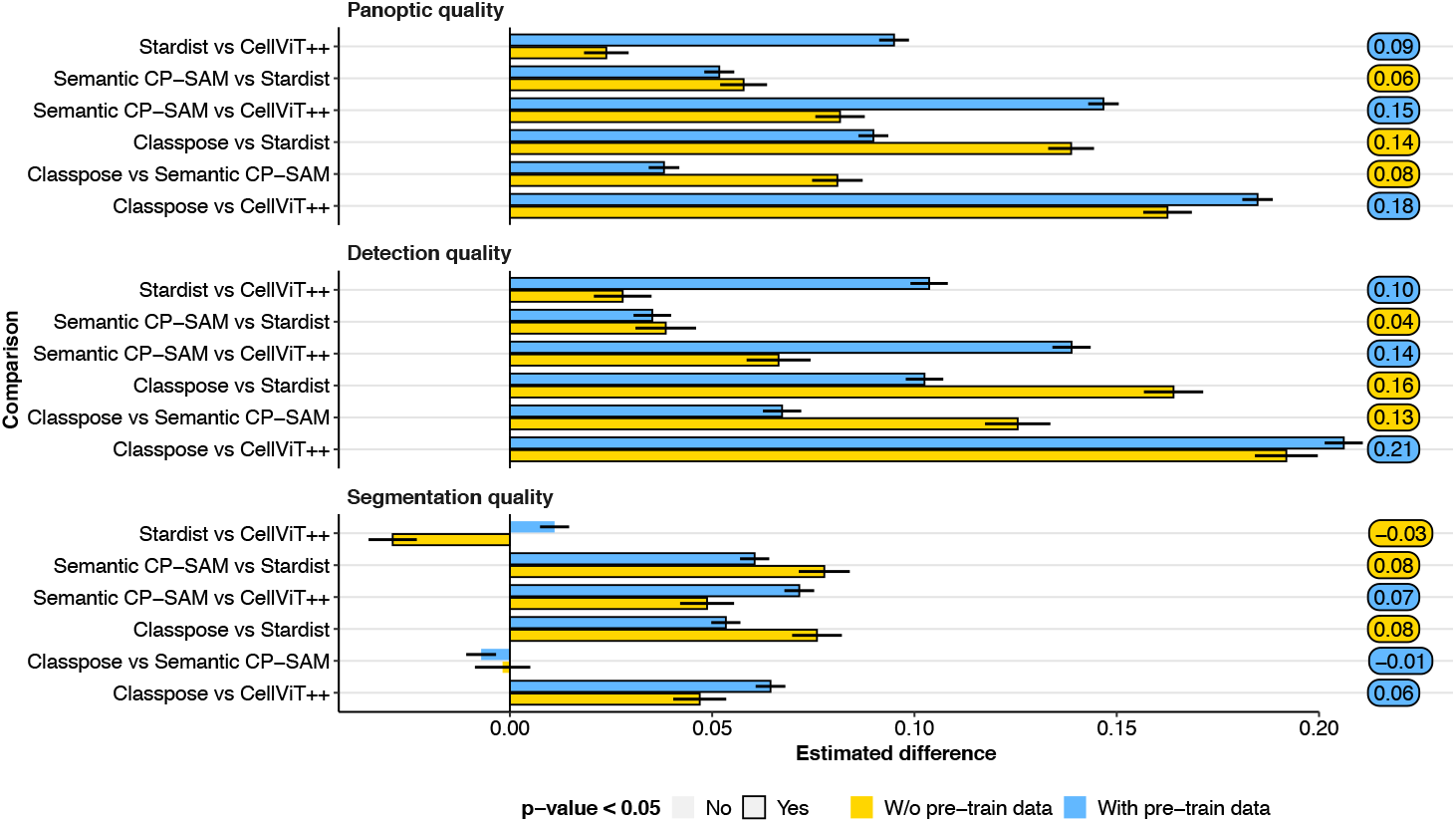
Comparison of pairwise comparisons between estimated marginal means (for Equation 5) for different models using all data (including datasets contained in the Cellpose-SAM pre-train data; “With pre-train data”) and using only datasets not used during Cellpose-SAM pre-training (“W/o pre-train data”). Black outlines indicate cases where the coefficients are statistically significant. The colours for the rounded rectangles indicate the set of datasets with the largest absolute coefficient value, and the value in the rounded rectangles indicates that coefficient value.

**Fig. S6.**
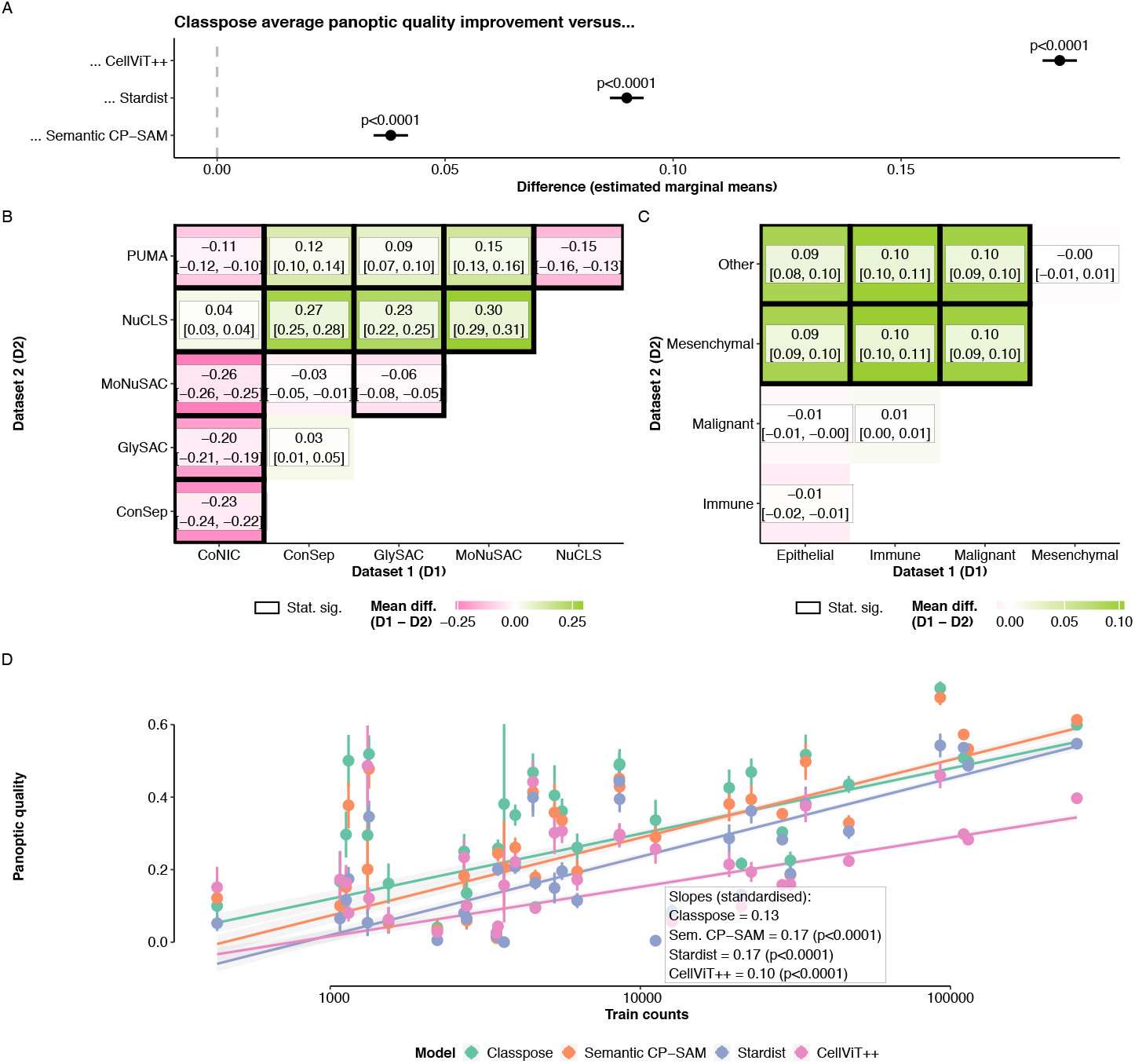
Multivariate analysis of factors underlying panoptic quality. **A**. Average difference in performance between Classpose and Semantic Cellpose-SAM, Stardist and CellViT++ according to t-tests comparing the estimated marginal means for all datasets including CoNIC and MoNuSAC. **B**. Average differences in panoptic quality between datasets according to t-tests comparing the estimated marginal means. A black box signifies whether specific dataset comparisons are statistically significant. **C**. Relationship between train proportions and the panoptic quality, stratified by model. p-values were calculated using Wald chi-squared tests comparing interaction term coefficients.

**Fig. S7.**
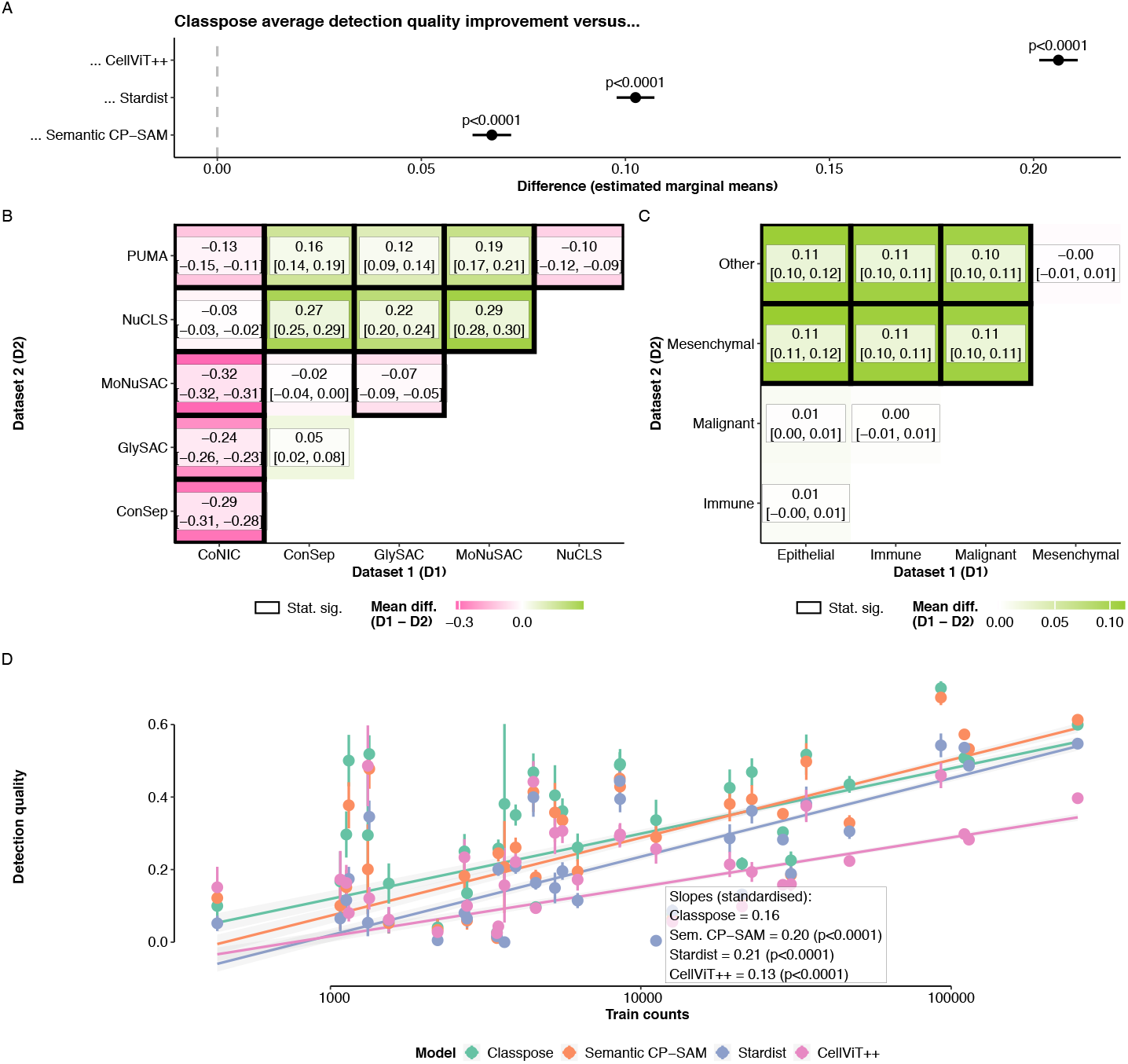
Multivariate analysis of factors underlying detection quality. **A**. Average difference in performance between Classpose and Semantic Cellpose-SAM, Stardist and CellViT++ according to t-tests comparing the estimated marginal means for all datasets including CoNIC and MoNuSAC. **B**. Average differences in detection quality between datasets according to t-tests comparing the estimated marginal means. A black box signifies whether specific dataset comparisons are statistically significant. **C**. Relationship between train proportions and the detection quality, stratified by model. p-values were calculated using Wald chi-squared tests comparing interaction term coefficients.

**Fig. S8.**
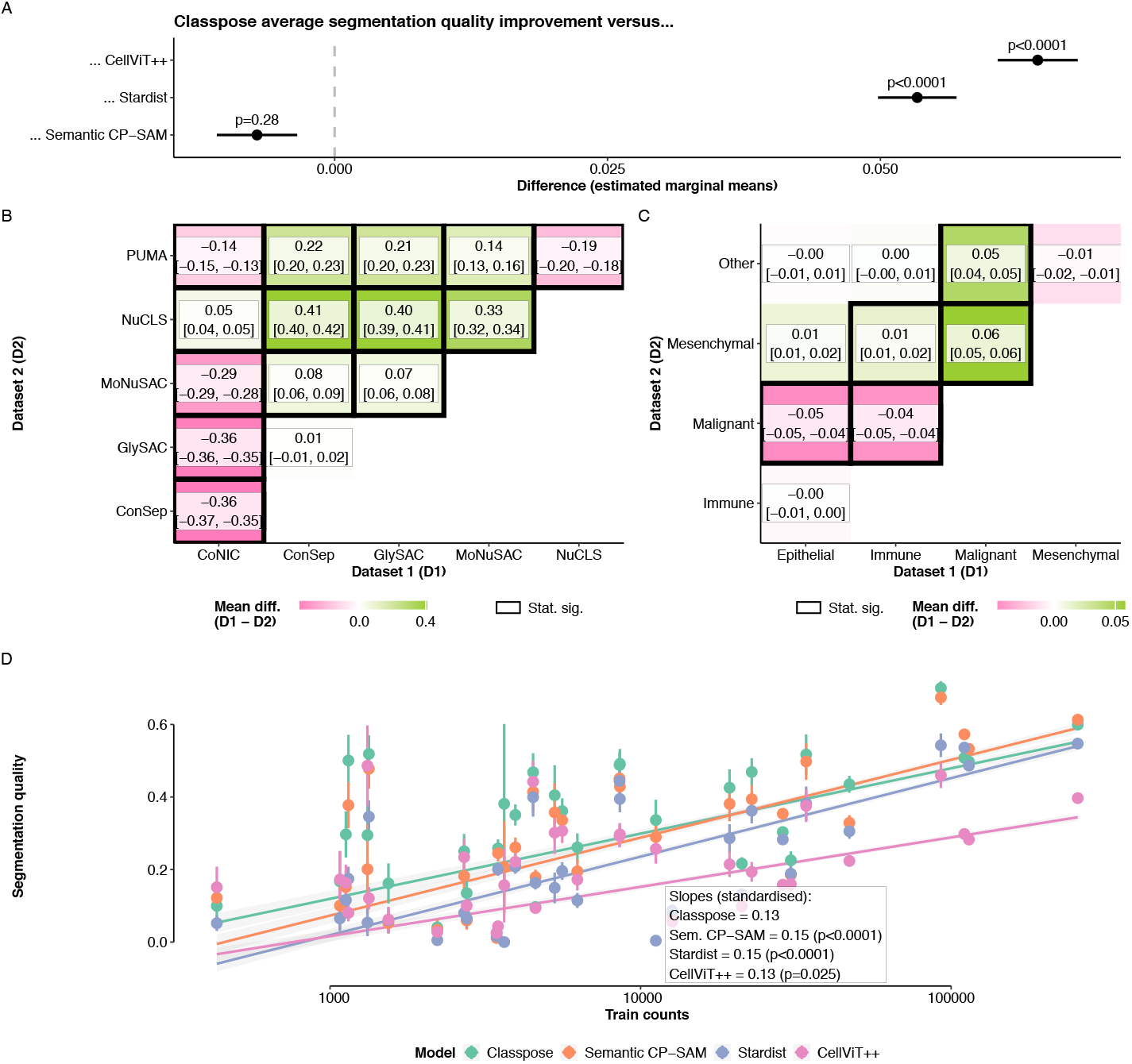
Multivariate analysis of factors underlying segmentation quality. **A**. Average difference in performance between Classpose and Semantic Cellpose-SAM, Stardist and CellViT++ according to t-tests comparing the estimated marginal means for all datasets including CoNIC and MoNuSAC. **B**. Average differences in segmentation quality between datasets according to t-tests comparing the estimated marginal means. A black box signifies whether specific dataset comparisons are statistically significant. **C**. Relationship between train proportions and the segmentation quality, stratified by model. p-values were calculated using Wald chi-squared tests comparing interaction term coefficients.

**Fig. S9.**
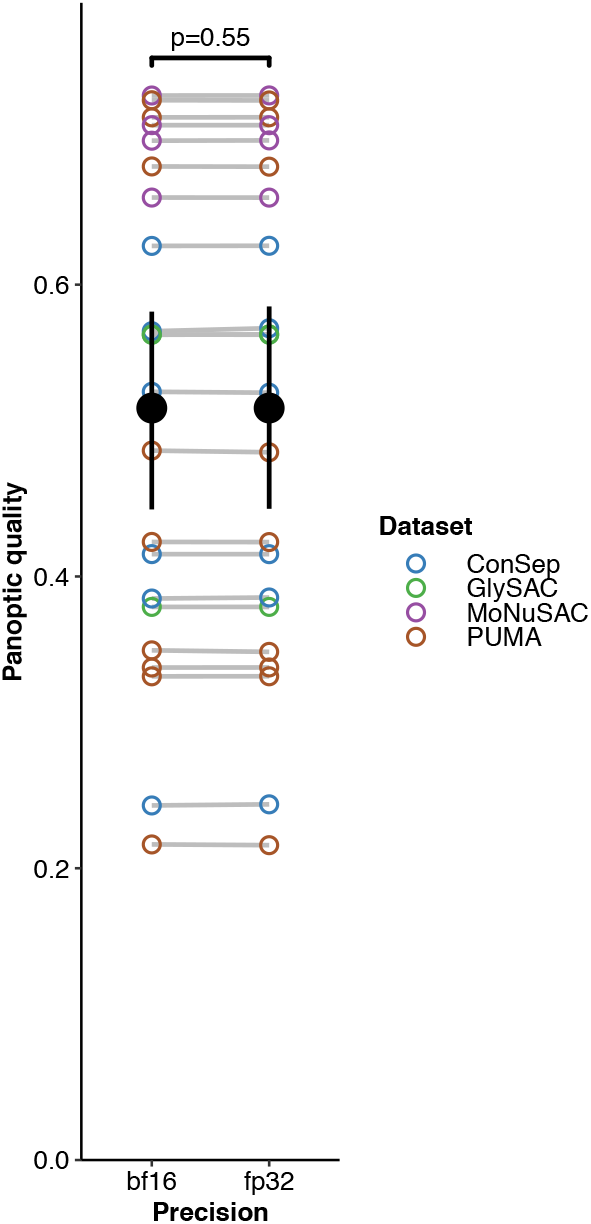
Comparison of per-class PQ for all cell class types.

**Fig. S10.**
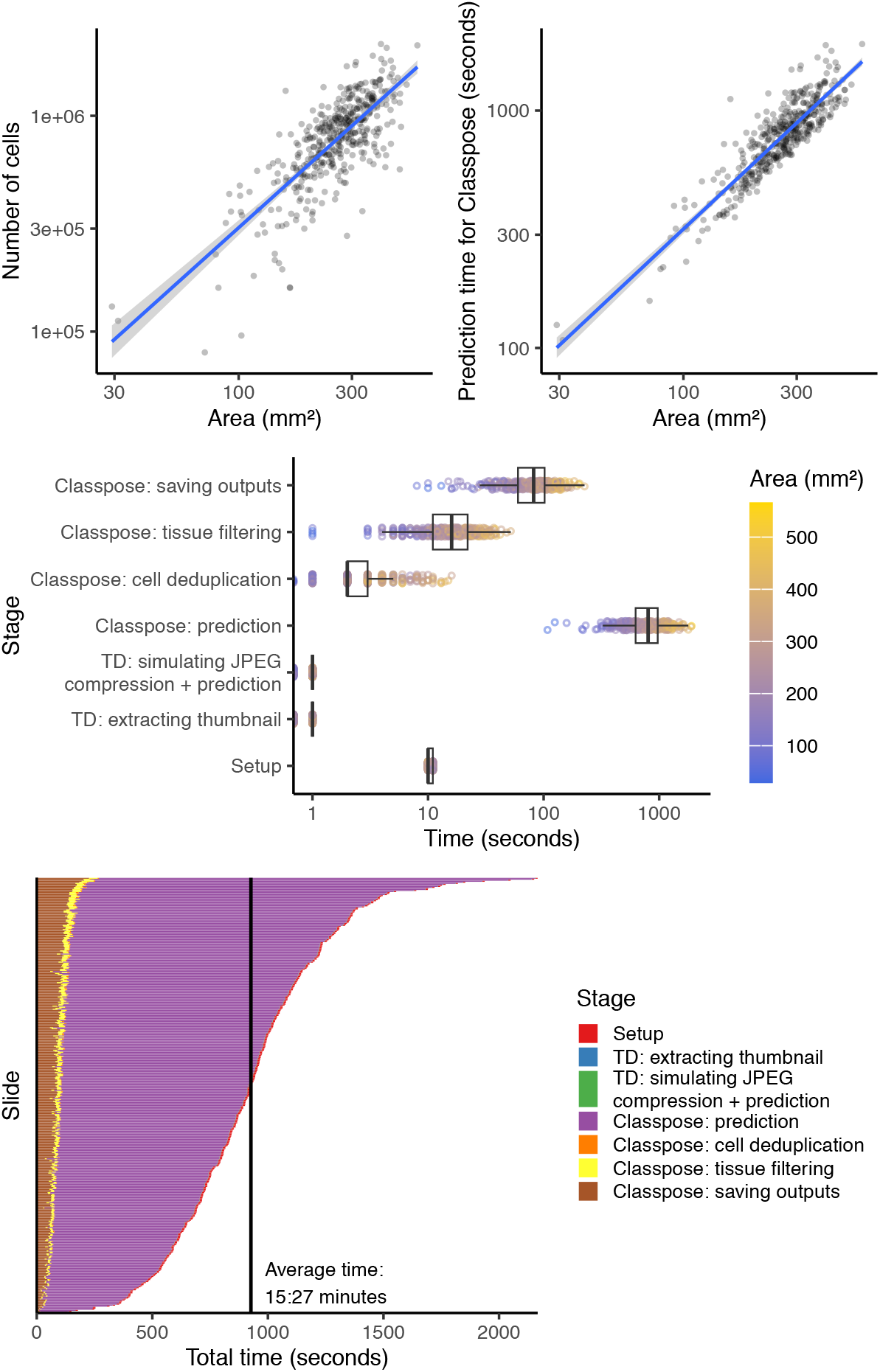
Performance benchmarking of SurGen predictions. **A**. Relationship between area and detected number of cells. **B** Relationship between area and prediction time. **C** Time allocation for the different stages of Classpose. **D** Time allocation for the different stages of Classpose stratified by slide.

**Fig. S11.**
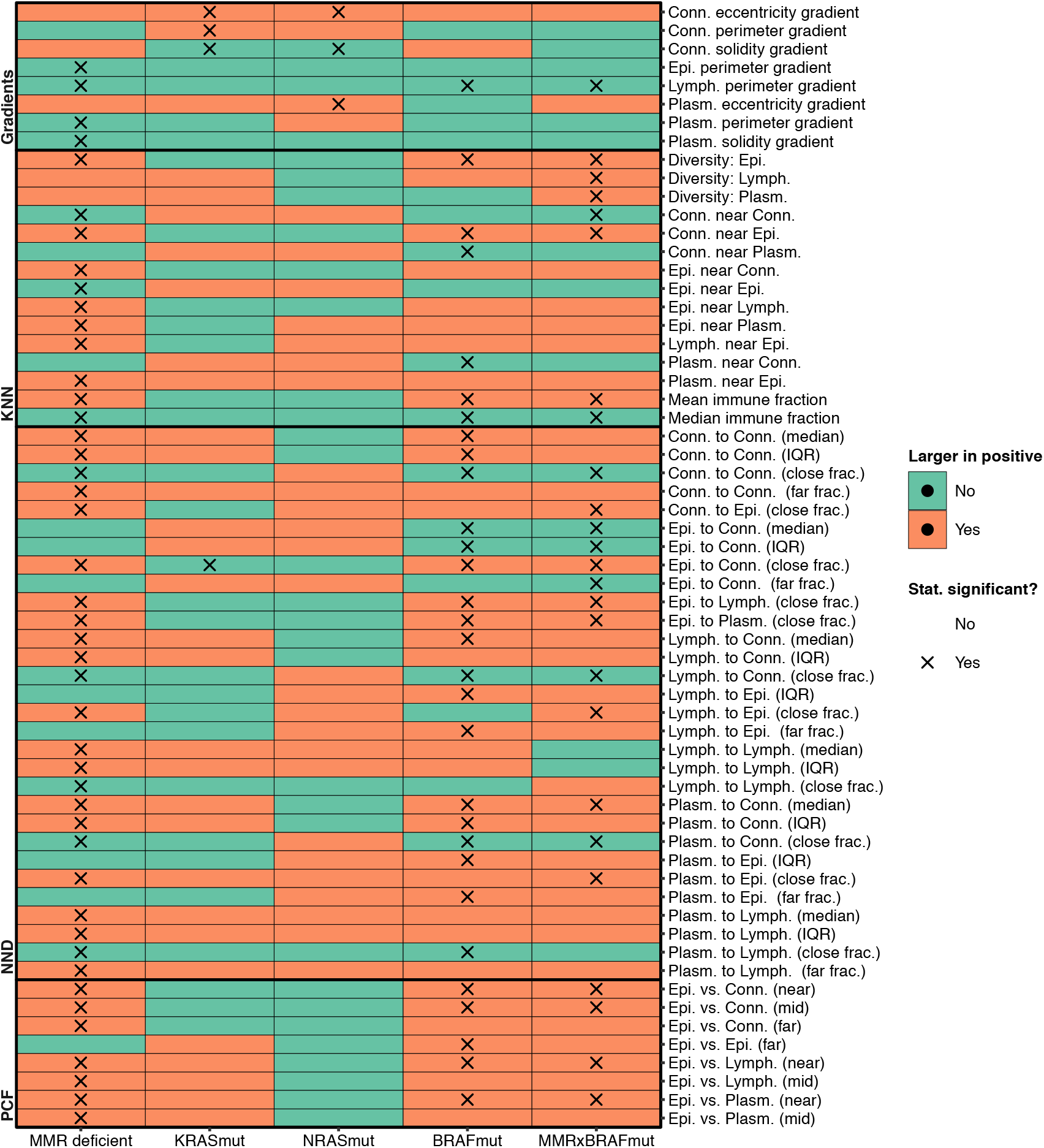
Summary of the associations between spatial and spatial-morphometric features. Only statistically significant correlations are shown.

**Fig. S12.**
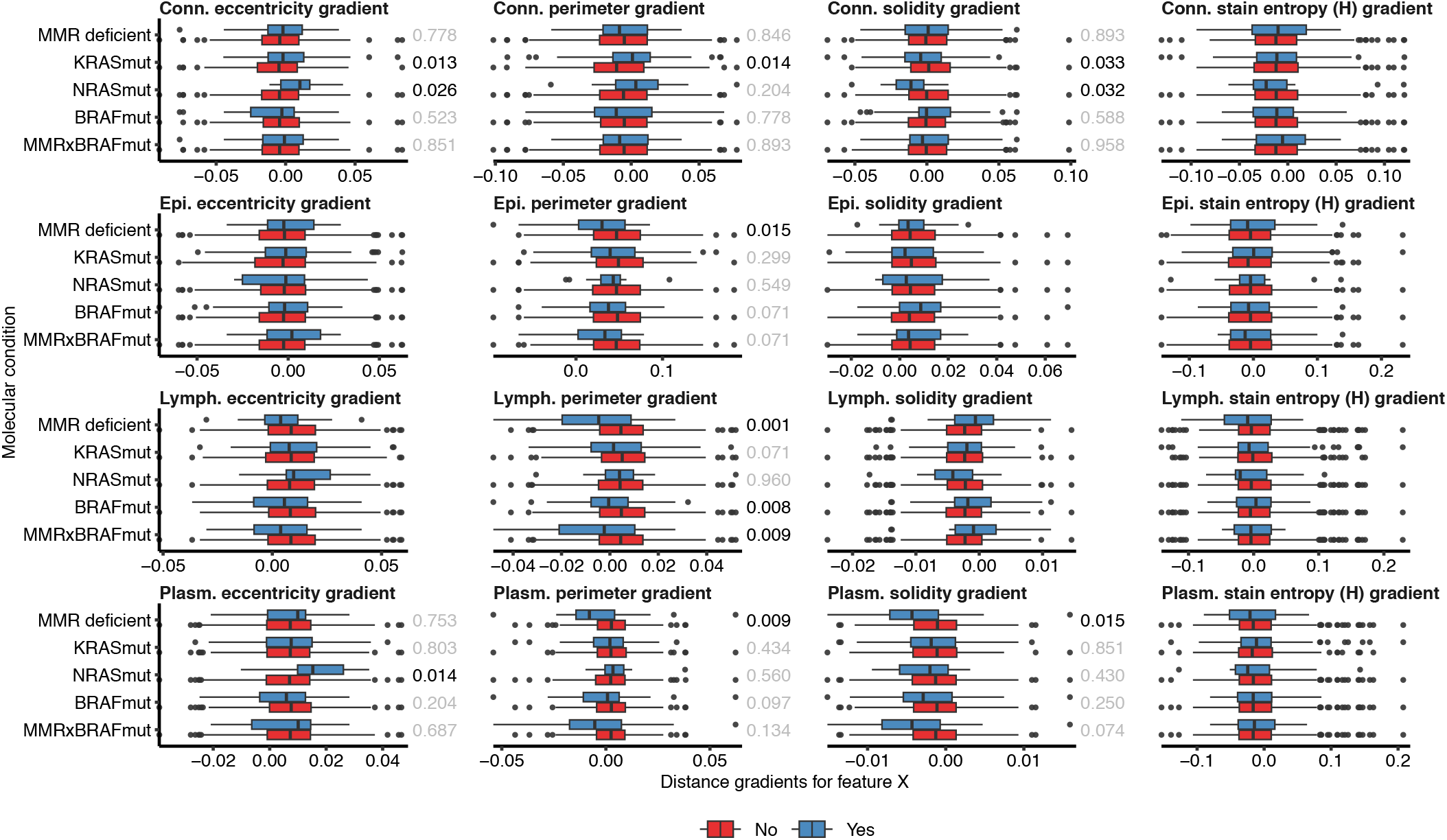
Association between pericancerous morphometric gradients and mutations/MMR status. For cases where the omnibus test was significant (*p <* 0.05), post-hoc comparisons were performed and the corresponding p-value is shown. The colour of the p-value is indicative of statistical significance (black implies statistical significance, whereas grey implies an absence of statistical significance).

**Fig. S13.**
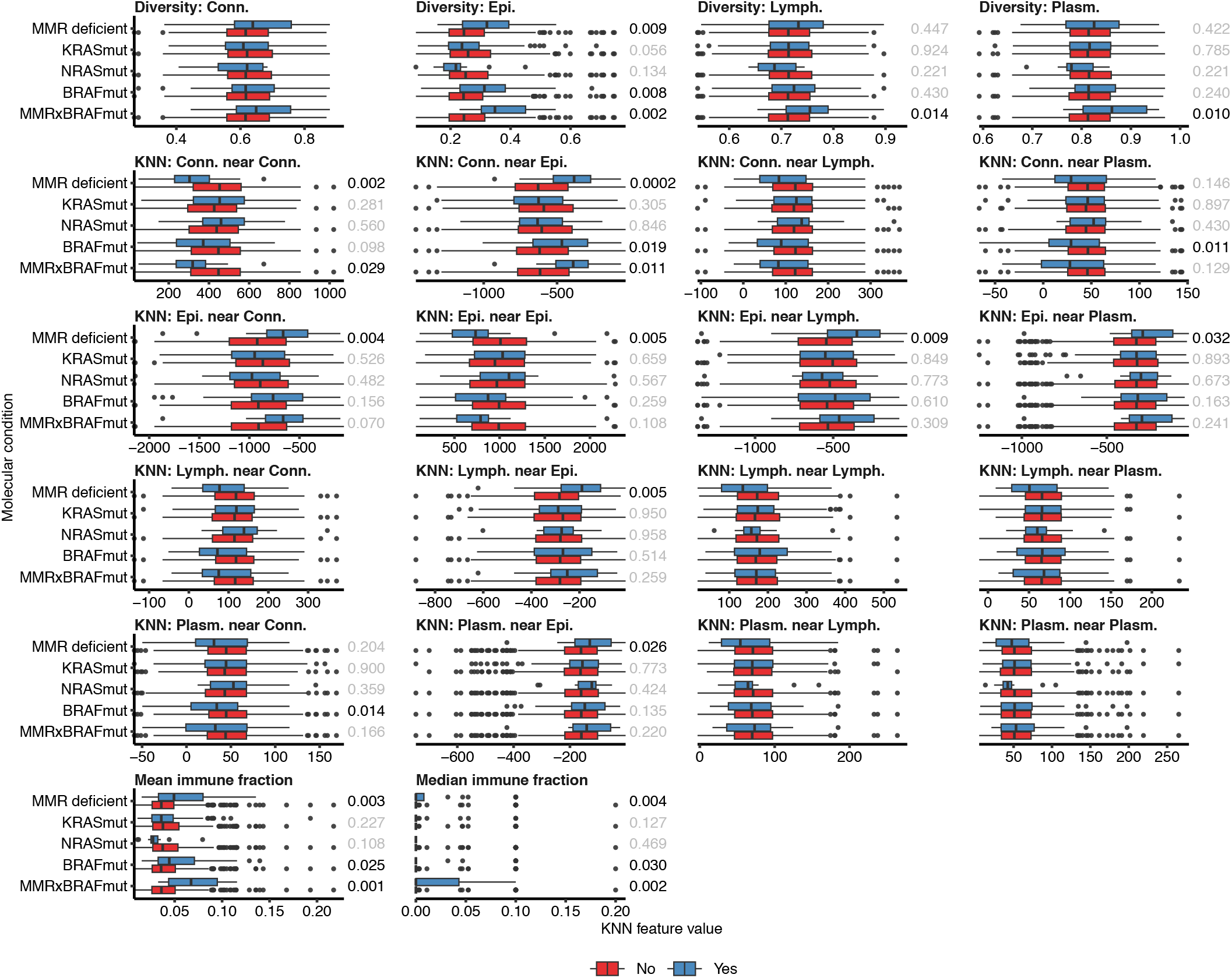
Association between KNN-based features and mutations/MMR status. For cases where the omnibus test was significant (*p <* 0.05), post-hoc comparisons were performed and the corresponding p-value is shown. The colour of the p-value is indicative of statistical significance (black implies statistical significance, whereas grey implies an absence of statistical significance).

**Fig. S14.**
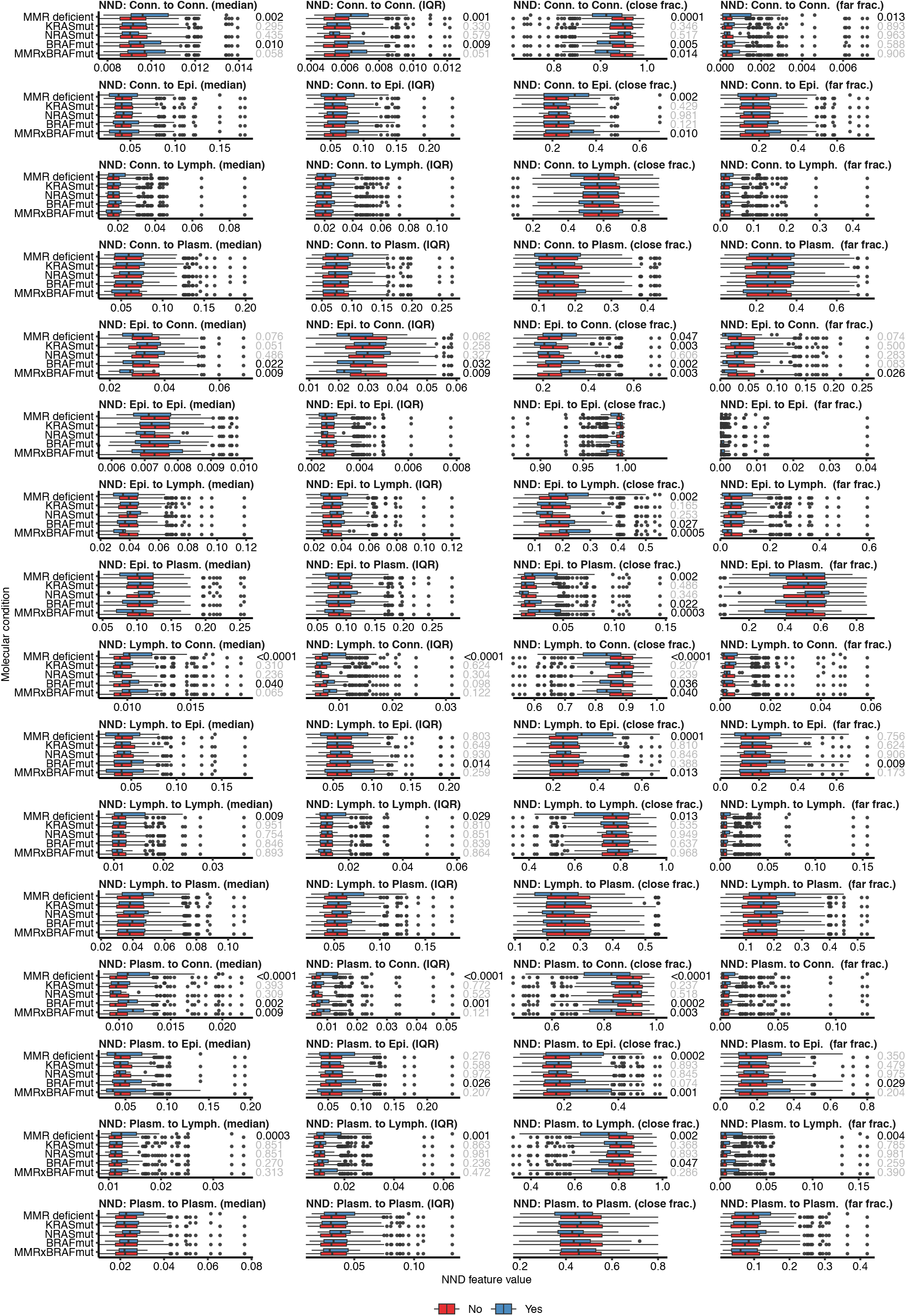
Association between NND-based features and mutations/MMR status. For cases where the omnibus test was significant (*p <* 0.05), post-hoc comparisons were performed and the corresponding p-value is shown. The colour of the p-value is indicative of statistical significance (black implies statistical significance, whereas grey implies an absence of statistical significance).

**Fig. S15.**
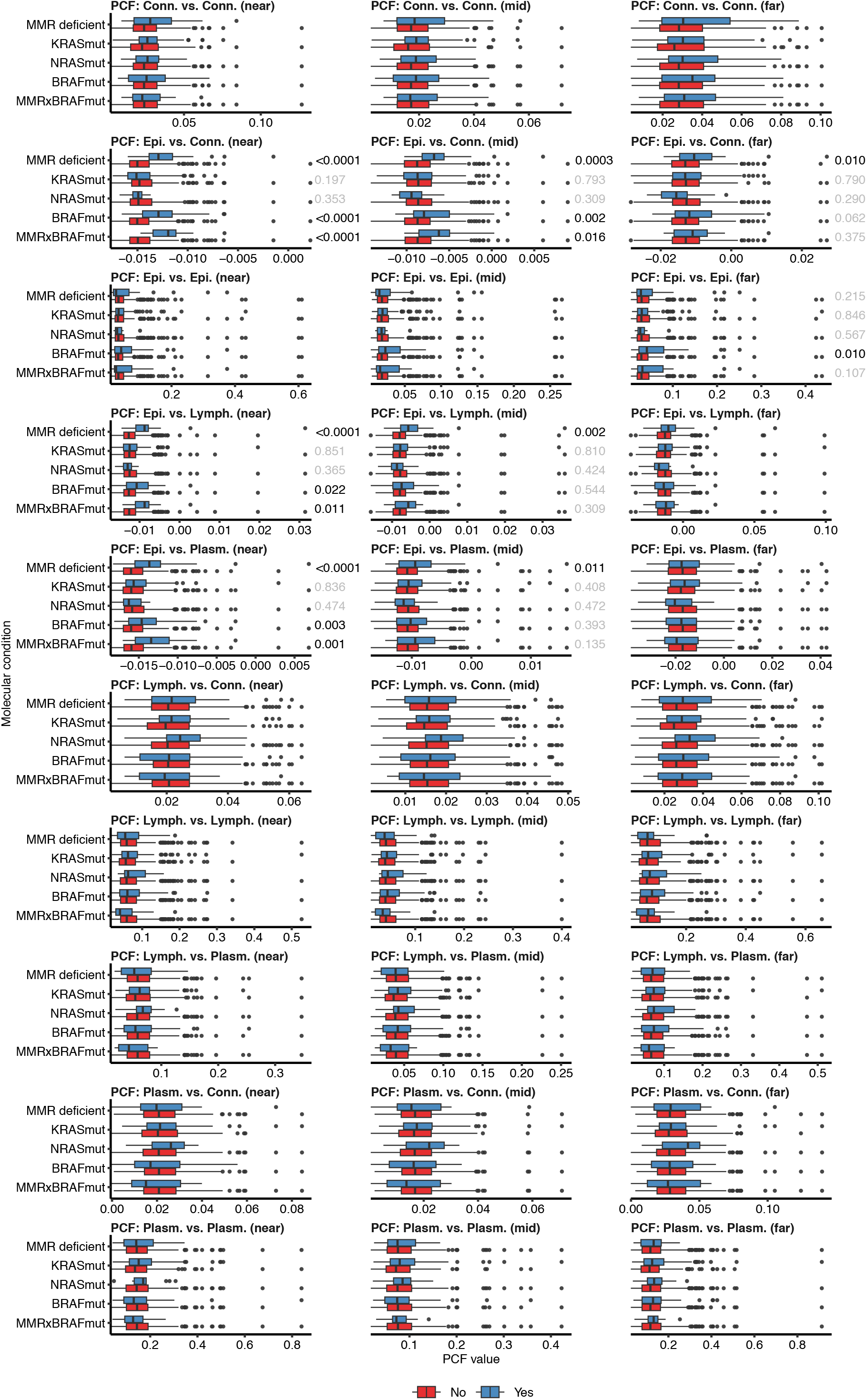
Association between PCF values and mutations/MMR status. For cases where the omnibus test was significant (*p <* 0.05), post-hoc comparisons were performed and the corresponding p-value is shown. The colour of the p-value is indicative of statistical significance (black implies statistical significance, whereas grey implies an absence of statistical significance).

**Fig. S16.**
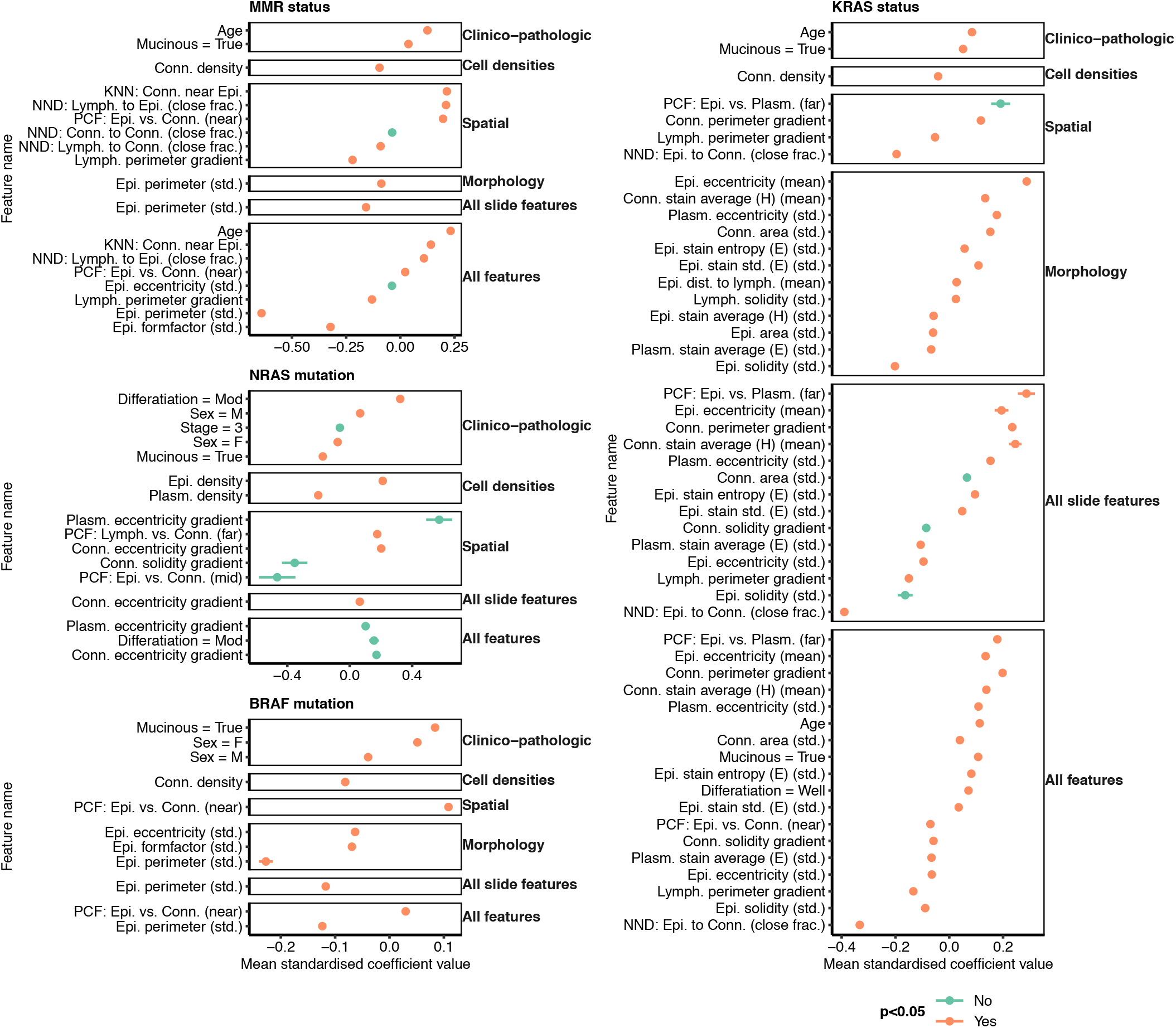
10-fold CV feature importance (standardised coefficients) distribution for features where more than half of the folds had standardised coefficients different from zero for the model using all morphological features. For E-H, the points represent the mean while the horizontal bars represent the standard error around the mean. Colours represent whether the coefficient distribution is different from zero according to a two-sided Wilcoxon test.

## S8 Supplementary Tables

**Table S1.**
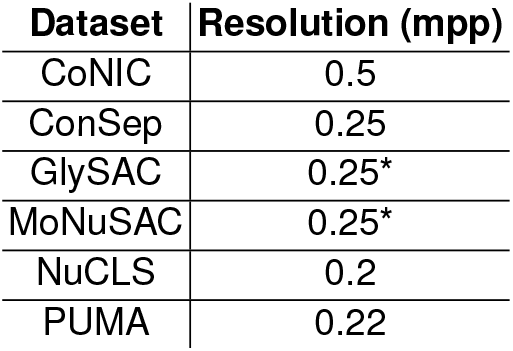
Data resolution for each dataset in microns per pixel (mpp). Values with a star (“*”) correspond to values where mpp information was not made available and correpond to data acquired at 40x magnification.

| Dataset | Resolution (mpp) |
| --- | --- |
| CoNIC | 0.5 |
| ConSep | 0.25 |
| GlySAC | 0.25* |
| MoNuSAC | 0.25* |
| NuCLS | 0.2 |
| PUMA | 0.22 |

**Table S2.** PQ results for Classpose and other methods used for benchmarking. Bold indicates the best performance for individual cell classes; the “Average” cell class is the average performance across all classes. CP-SAM = Cellpose-SAM.

| Cell class | CellViT++<br>(HIPT256) | CellViT++<br>(SAM-H) | CellViT++<br>(Virchow) | Stardist | Semantic<br>CP-SAM | Classpose |
| --- | --- | --- | --- | --- | --- | --- |
| **Unbiased datasets** |  |  |  |  |  |  |
| **ConSep** |  |  |  |  |  |  |
| Muscle | 0.251 | 0.31 | 0.346 | 0.211 | 0.234 | **0.386** |
| Stroma | 0.169 | 0.221 | 0.249 | 0.254 | 0.373 | **0.415** |
| Malignant epithelial | 0.268 | 0.31 | 0.31 | 0.277 | 0.489 | **0.526** |
| Healthy epithelial | 0.378 | 0.458 | 0.462 | 0.156 | 0.365 | **0.57** |
| Inflammatory | 0.343 | 0.374 | 0.357 | 0.557 | 0.554 | **0.627** |
| Other | 0.195 | 0.226 | **0.25** | 0.135 | 0.136 | 0.244 |
| Average | 0.267 | 0.316 | 0.329 | 0.265 | 0.359 | **0.461** |
| **GlySAC** |  |  |  |  |  |  |
| Other | 0.174 | 0.205 | 0.226 | 0.224 | 0.291 | **0.379** |
| Lymphocyte | 0.264 | 0.276 | 0.285 | 0.526 | 0.521 | **0.566** |
| Epithelial | 0.281 | 0.303 | 0.297 | 0.47 | 0.505 | **0.566** |
| Average | 0.24 | 0.261 | 0.269 | 0.407 | 0.439 | **0.504** |
| **NuCLS** |  |  |  |  |  |  |
| Plasma cells | 0.065 | 0.108 | 0.141 | **0.23** | 0.158 | 0.155 |
| Tumor | 0.32 | 0.334 | 0.317 | 0.411 | 0.435 | **0.527** |
| Macrophage | 0.037 | 0.033 | **0.06** | 0.012 | 0.017 | 0.028 |
| Stroma | 0.186 | 0.203 | 0.167 | 0.242 | 0.204 | **0.343** |
| Other | 0.019 | 0.029 | **0.042** | 0.006 | 0.035 | 0.036 |
| Lymphocyte | 0.281 | 0.307 | 0.307 | 0.369 | 0.362 | **0.447** |
| Average | 0.151 | 0.169 | 0.172 | 0.212 | 0.202 | **0.256** |
| **PUMA** |  |  |  |  |  |  |
| Melanophage | 0.106 | 0.264 | 0.185 | 0.143 | 0.151 | **0.338** |
| Histiocyte | 0.292 | 0.326 | 0.357 | 0.004 | 0.396 | **0.485** |
| Endothelial | 0.167 | 0.194 | 0.293 | 0.116 | 0.242 | **0.332** |
| Apoptosis | 0.088 | 0.12 | 0.142 | 0.089 | 0.13 | **0.216** |
| Tumor | 0.41 | 0.411 | 0.416 | 0.576 | 0.695 | **0.715** |
| Stroma | 0.208 | 0.248 | 0.236 | 0.129 | 0.267 | **0.348** |
| Other | 0.132 | 0 | 0.208 | 0.114 | 0.157 | **0.424** |
| Lymphocyte | 0.396 | 0.408 | 0.44 | 0.545 | 0.648 | **0.681** |
| Epithelial | 0.513 | 0.391 | 0.604 | 0 | 0.488 | **0.726** |
| Average | 0.257 | 0.262 | 0.32 | 0.191 | 0.353 | **0.474** |
| **Biased datasets** |  |  |  |  |  |  |
| **CoNIC** |  |  |  |  |  |  |
| Connective | 0.274 | 0.334 | 0.328 | 0.53 | **0.586** | 0.551 |
| Eosinophil | 0.1 | 0.093 | 0.077 | 0.309 | **0.395** | 0.361 |
| Plasma | 0.153 | 0.208 | 0.242 | 0.384 | **0.468** | 0.432 |
| Lymphocyte | 0.31 | 0.367 | 0.344 | 0.61 | **0.644** | 0.614 |
| Epithelial | 0.336 | 0.387 | 0.409 | 0.599 | **0.631** | 0.62 |
| Neutrophil | 0.058 | 0.098 | 0.198 | 0.271 | **0.29** | 0.185 |
| Average | 0.205 | 0.248 | 0.267 | 0.45 | **0.502** | 0.461 |
| **MoNuSAC** |  |  |  |  |  |  |
| Macrophage | 0.086 | 0.288 | 0.182 | 0.216 | 0.572 | **0.709** |
| Lymphocyte | 0.335 | 0.344 | 0.29 | 0.651 | 0.688 | **0.699** |
| Epithelial | 0.19 | 0.368 | 0.325 | 0.61 | 0.707 | **0.73** |
| Neutrophil | 0.069 | 0.096 | 0.133 | 0.473 | 0.612 | **0.66** |
| Average | 0.17 | 0.274 | 0.232 | 0.487 | 0.645 | **0.699** |

**Table S3.** Detection quality results for Classpose and other methods used for benchmarking. Bold indicates the best performance for individual cell classes; the “Average” cell class is the average performance across all classes.

| Cell class | CellViT++<br>(HIPT256) | CellViT++<br>(SAM-H) | CellViT++<br>(Virchow) | Stardist | Semantic<br>CP-SAM | Classpose |
| --- | --- | --- | --- | --- | --- | --- |
| **CoNIC** |  |  |  |  |  |  |
| Connective | 0.355 | 0.434 | 0.427 | 0.66 | **0.706** | 0.672 |
| Eosinophil | 0.145 | 0.137 | 0.115 | 0.417 | **0.502** | 0.452 |
| Plasma | 0.213 | 0.284 | 0.332 | 0.449 | **0.549** | 0.507 |
| Lymphocyte | 0.411 | 0.477 | 0.455 | 0.719 | **0.74** | 0.709 |
| Epithelial | 0.461 | 0.523 | 0.557 | 0.776 | **0.805** | 0.79 |
| Neutrophil | 0.08 | 0.133 | 0.273 | 0.356 | **0.375** | 0.245 |
| Average | 0.278 | 0.332 | 0.36 | 0.563 | **0.613** | 0.563 |
| **ConSep** |  |  |  |  |  |  |
| Muscle | 0.349 | 0.403 | 0.462 | 0.312 | 0.312 | **0.509** |
| Stroma | 0.241 | 0.304 | 0.342 | 0.351 | 0.492 | **0.545** |
| Malignant epithelial | 0.373 | 0.422 | 0.425 | 0.38 | 0.623 | **0.656** |
| Healthy epithelial | 0.497 | 0.59 | 0.592 | 0.204 | 0.465 | **0.711** |
| Inflammatory | 0.43 | 0.47 | 0.446 | 0.691 | 0.668 | **0.752** |
| Other | 0.269 | 0.308 | **0.344** | 0.187 | 0.185 | 0.335 |
| Average | 0.36 | 0.416 | 0.435 | 0.354 | 0.457 | **0.585** |
| **GlySAC** |  |  |  |  |  |  |
| Other | 0.25 | 0.293 | 0.321 | 0.302 | 0.377 | **0.487** |
| Lymphocyte | 0.356 | 0.371 | 0.382 | 0.661 | 0.639 | **0.699** |
| Epithelial | 0.385 | 0.416 | 0.411 | 0.605 | 0.636 | **0.701** |
| Average | 0.33 | 0.36 | 0.371 | 0.523 | 0.55 | **0.629** |
| **MoNuSAC** |  |  |  |  |  |  |
| Macrophage | 0.107 | 0.348 | 0.24 | 0.289 | 0.665 | **0.82** |
| Lymphocyte | 0.454 | 0.47 | 0.397 | 0.84 | 0.869 | **0.887** |
| Epithelial | 0.25 | 0.482 | 0.431 | 0.772 | 0.846 | **0.869** |
| Neutrophil | 0.089 | 0.125 | 0.179 | 0.593 | 0.755 | **0.806** |
| Average | 0.225 | 0.356 | 0.312 | 0.624 | 0.784 | **0.845** |
| **NuCLS** |  |  |  |  |  |  |
| Plasma cells | 0.1 | 0.161 | 0.212 | **0.341** | 0.213 | 0.222 |
| Tumor | 0.474 | 0.494 | 0.478 | 0.6 | 0.596 | **0.774** |
| Macrophage | 0.057 | 0.046 | **0.094** | 0.021 | 0.022 | 0.045 |
| Stroma | 0.283 | 0.311 | 0.259 | 0.394 | 0.299 | **0.539** |
| Other | 0.028 | 0.048 | **0.073** | 0.011 | 0.058 | 0.071 |
| Lymphocyte | 0.416 | 0.465 | 0.466 | 0.537 | 0.475 | **0.621** |
| Average | 0.227 | 0.254 | 0.263 | 0.317 | 0.277 | **0.379** |
| **PUMA** |  |  |  |  |  |  |
| Melanophage | 0.145 | 0.354 | 0.27 | 0.2 | 0.189 | **0.436** |
| Histocyte | 0.379 | 0.423 | 0.463 | 0.004 | 0.467 | **0.58** |
| Endothelial | 0.228 | 0.266 | 0.385 | 0.153 | 0.299 | **0.413** |
| Apoptosis | 0.127 | 0.175 | 0.216 | 0.121 | 0.175 | **0.292** |
| Tumor | 0.551 | 0.55 | 0.559 | 0.732 | 0.824 | **0.858** |
| Stroma | 0.279 | 0.337 | 0.323 | 0.181 | 0.33 | **0.448** |
| Other | 0.174 | 0 | 0.275 | 0.163 | 0.21 | **0.522** |
| Lymphocyte | 0.519 | 0.532 | 0.575 | 0.66 | 0.744 | **0.8** |
| Epithelial | 0.617 | 0.464 | 0.717 | 0 | 0.573 | **0.828** |
| Average | 0.335 | 0.345 | 0.42 | 0.246 | 0.423 | **0.575** |

**Table S4.** Segmentation quality results for Classpose and other methods used for benchmarking. Bold indicates the best performance for individual cell classes; the “Average” cell class is the average performance across all classes.

| Cell class | CellViT++<br>(HIPT256) | CellViT++<br>(SAM-H) | CellViT++<br>(Virchow) | Stardist | Semantic<br>CP-SAM | Classpose |
| --- | --- | --- | --- | --- | --- | --- |
| **CoNIC** |  |  |  |  |  |  |
| Connective | 0.773 | 0.769 | 0.768 | 0.804 | **0.829** | 0.82 |
| Eosinophil | 0.691 | 0.678 | 0.67 | 0.741 | 0.788 | **0.799** |
| Plasma | 0.72 | 0.733 | 0.729 | **0.855** | 0.854 | 0.853 |
| Lymphocyte | 0.755 | 0.77 | 0.757 | 0.848 | **0.87** | 0.866 |
| Epithelial | 0.73 | 0.739 | 0.735 | 0.772 | **0.784** | **0.784** |
| Neutrophil | 0.721 | 0.738 | 0.726 | 0.762 | **0.774** | 0.756 |
| Average | 0.732 | 0.738 | 0.731 | 0.797 | **0.817** | 0.813 |
| **ConSep** |  |  |  |  |  |  |
| Muscle | 0.72 | **0.768** | 0.748 | 0.676 | 0.751 | 0.758 |
| Stroma | 0.7 | 0.726 | 0.728 | 0.723 | 0.758 | **0.761** |
| Malignant epithelial | 0.72 | 0.735 | 0.73 | 0.728 | 0.785 | **0.802** |
| Healthy epithelial | 0.761 | 0.777 | 0.78 | 0.767 | 0.785 | **0.802** |
| Inflammatory | 0.798 | 0.795 | 0.799 | 0.807 | 0.83 | **0.834** |
| Other | 0.727 | 0.733 | 0.726 | 0.723 | **0.734** | 0.727 |
| Average | 0.738 | 0.756 | 0.752 | 0.737 | 0.774 | **0.781** |
| **GlySAC** |  |  |  |  |  |  |
| Other | 0.695 | 0.698 | 0.705 | 0.741 | 0.773 | **0.778** |
| Lymphocyte | 0.741 | 0.744 | 0.745 | 0.795 | **0.815** | 0.81 |
| Epithelial | 0.73 | 0.729 | 0.724 | 0.777 | 0.794 | **0.807** |
| Average | 0.722 | 0.724 | 0.725 | 0.771 | 0.794 | **0.798** |
| **MoNuSAC** |  |  |  |  |  |  |
| Macrophage | 0.803 | 0.827 | 0.76 | 0.745 | 0.859 | **0.865** |
| Lymphocyte | 0.738 | 0.731 | 0.731 | 0.775 | **0.792** | 0.788 |
| Epithelial | 0.763 | 0.764 | 0.752 | 0.79 | 0.836 | **0.84** |
| Neutrophil | 0.773 | 0.764 | 0.739 | 0.797 | 0.811 | **0.818** |
| Average | 0.769 | 0.771 | 0.746 | 0.777 | 0.825 | **0.828** |
| **NuCLS** |  |  |  |  |  |  |
| Plasma cells | 0.651 | 0.671 | 0.666 | 0.675 | **0.742** | 0.7 |
| Tumor | 0.674 | 0.675 | 0.662 | 0.684 | **0.73** | 0.681 |
| Macrophage | 0.652 | 0.729 | 0.64 | 0.566 | **0.793** | 0.624 |
| Stroma | 0.657 | 0.651 | 0.647 | 0.613 | **0.681** | 0.637 |
| Other | **0.668** | 0.608 | 0.569 | 0.581 | 0.612 | 0.512 |
| Lymphocyte | 0.676 | 0.662 | 0.658 | 0.686 | **0.762** | 0.72 |
| Average | 0.663 | 0.666 | 0.64 | 0.634 | **0.72** | 0.646 |
| **PUMA** |  |  |  |  |  |  |
| Melanophage | 0.729 | 0.745 | 0.686 | 0.714 | **0.799** | 0.775 |
| Histocyte | 0.771 | 0.771 | 0.77 | 0.787 | **0.849** | 0.837 |
| Endothelial | 0.731 | 0.728 | 0.761 | 0.759 | **0.81** | 0.803 |
| Apoptosis | 0.695 | 0.689 | 0.656 | 0.733 | **0.744** | 0.739 |
| Tumor | 0.745 | 0.748 | 0.745 | 0.787 | **0.843** | 0.833 |
| Stroma | 0.745 | 0.734 | 0.732 | 0.711 | **0.809** | 0.778 |
| Other | 0.756 | 0 | 0.755 | 0.702 | 0.749 | **0.812** |
| Lymphocyte | 0.763 | 0.767 | 0.766 | 0.825 | **0.871** | 0.852 |
| Epithelial | 0.831 | 0.843 | 0.842 | 0 | 0.85 | **0.877** |
| Average | 0.752 | 0.669 | 0.746 | 0.669 | **0.814** | 0.812 |

**Table S5.**
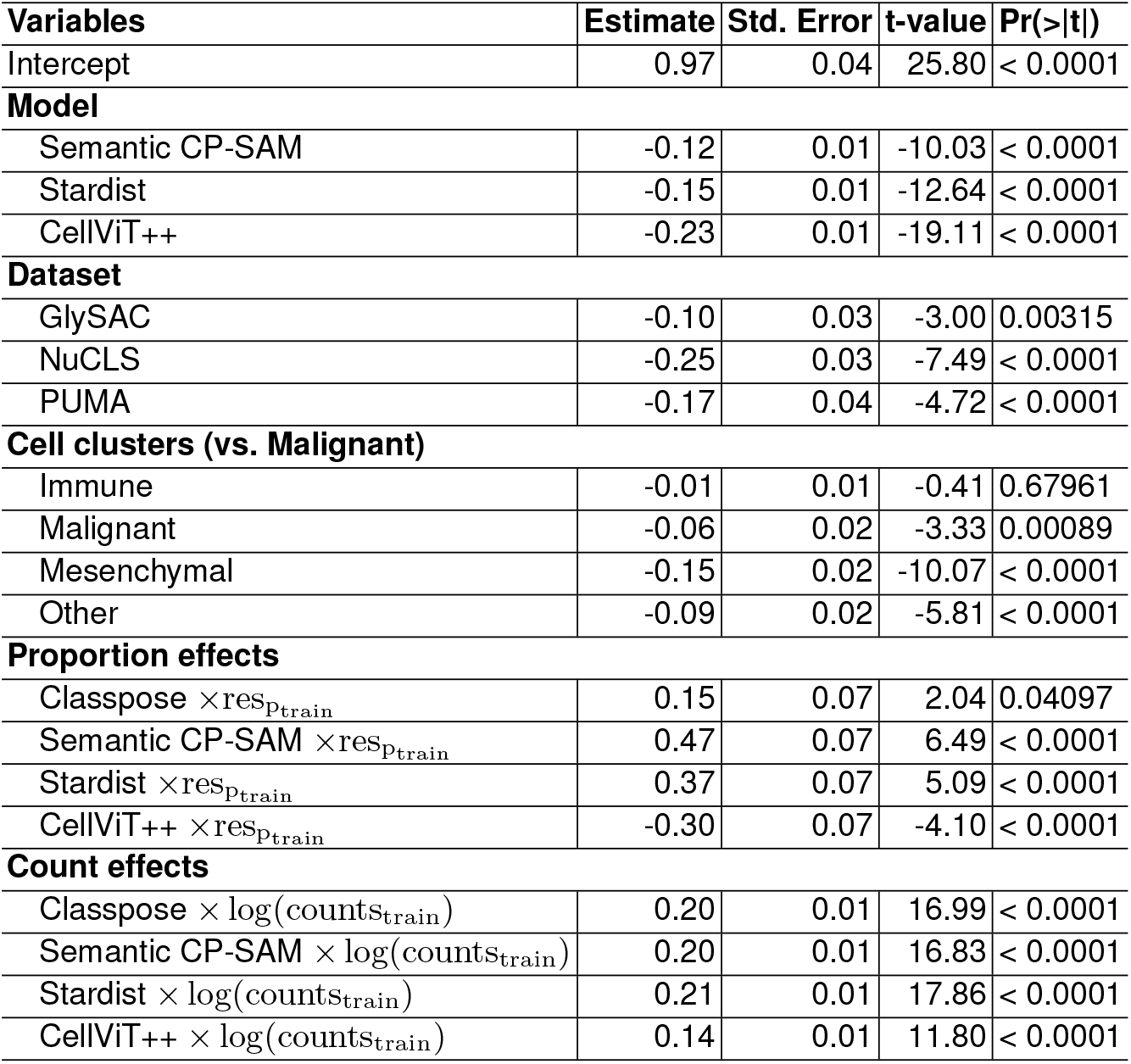
Coefficient estimates and their respective standard errors (Std. Error), t-values and statistical significance for a two-sided t-test for the linear model described in Equation 5 for DQ.

| Variables | Estimate | Std. Error | t-value | Pr(>\|t\|) |
| --- | --- | --- | --- | --- |
| Intercept | 0.97 | 0.04 | 25.80 | < 0.0001 |
| **Model** |  |  |  |  |
| Semantic CP-SAM | -0.12 | 0.01 | -10.03 | < 0.0001 |
| Stardist | -0.15 | 0.01 | -12.64 | < 0.0001 |
| CellViT++ | -0.23 | 0.01 | -19.11 | < 0.0001 |
| **Dataset** |  |  |  |  |
| GlySAC | -0.10 | 0.03 | -3.00 | 0.00315 |
| NuCLS | -0.25 | 0.03 | -7.49 | < 0.0001 |
| PUMA | -0.17 | 0.04 | -4.72 | < 0.0001 |
| **Cell clusters (vs. Malignant)** |  |  |  |  |
| Immune | -0.01 | 0.01 | -0.41 | 0.67961 |
| Malignant | -0.06 | 0.02 | -3.33 | 0.00089 |
| Mesenchymal | -0.15 | 0.02 | -10.07 | < 0.0001 |
| Other | -0.09 | 0.02 | -5.81 | < 0.0001 |
| **Proportion effects** |  |  |  |  |
| Classpose × res<sub>p<sub>train</sub></sub> | 0.15 | 0.07 | 2.04 | 0.04097 |
| Semantic CP-SAM × res<sub>p<sub>train</sub></sub> | 0.47 | 0.07 | 6.49 | < 0.0001 |
| Stardist × res<sub>p<sub>train</sub></sub> | 0.37 | 0.07 | 5.09 | < 0.0001 |
| CellViT++ × res<sub>p<sub>train</sub></sub> | -0.30 | 0.07 | -4.10 | < 0.0001 |
| **Count effects** |  |  |  |  |
| Classpose × log(counts<sub>train</sub>) | 0.20 | 0.01 | 16.99 | < 0.0001 |
| Semantic CP-SAM × log(counts<sub>train</sub>) | 0.20 | 0.01 | 16.83 | < 0.0001 |
| Stardist × log(counts<sub>train</sub>) | 0.21 | 0.01 | 17.86 | < 0.0001 |
| CellViT++ × log(counts<sub>train</sub>) | 0.14 | 0.01 | 11.80 | < 0.0001 |

**Table S6.** Coefficient estimates and their respective standard errors (Std. Error), t-values and statistical significance for a two-sided t-test for the linear model described in Equation 5 for SQ.

| Variables | Estimate | Std. Error | t-value | Pr(>\|t\|) |
| --- | --- | --- | --- | --- |
| Intercept | 1.06 | 0.03 | 38.29 | < 0.0001 |
| **Model** |  |  |  |  |
| Semantic CP-SAM | 0.02 | 0.01 | 1.89 | 0.05860 |
| Stardist | -0.07 | 0.01 | -6.54 | < 0.0001 |
| CellViT++ | -0.06 | 0.01 | -5.69 | < 0.0001 |
| **Dataset** |  |  |  |  |
| GlySAC | -0.05 | 0.02 | -2.23 | 0.02746 |
| NuCLS | -0.40 | 0.02 | -16.99 | < 0.0001 |
| PUMA | -0.23 | 0.02 | -9.39 | < 0.0001 |
| **Cell clusters (vs. Malignant)** |  |  |  |  |
| Immune | -0.03 | 0.01 | -2.50 | 0.01241 |
| Malignant | -0.02 | 0.02 | -1.38 | 0.16846 |
| Mesenchymal | -0.05 | 0.01 | -3.61 | 0.00031 |
| Other | 0.00 | 0.01 | 0.24 | 0.80838 |
| **Proportion effects** |  |  |  |  |
| Classpose × res<sub>p<sub>train</sub></sub> | -0.26 | 0.06 | -4.29 | < 0.0001 |
| Semantic CP-SAM × res<sub>p<sub>train</sub></sub> | -0.29 | 0.06 | -4.79 | < 0.0001 |
| Stardist × res<sub>p<sub>train</sub></sub> | -0.06 | 0.06 | -1.02 | 0.31006 |
| CellViT++ × res<sub>p<sub>train</sub></sub> | -0.44 | 0.06 | -7.17 | < 0.0001 |
| **Count effects** |  |  |  |  |
| Classpose × log(counts<sub>train</sub>) | 0.15 | 0.01 | 14.73 | < 0.0001 |
| Semantic CP-SAM × log(counts<sub>train</sub>) | 0.17 | 0.01 | 16.96 | < 0.0001 |
| Stardist × log(counts<sub>train</sub>) | 0.16 | 0.01 | 15.92 | < 0.0001 |
| CellViT++ × log(counts<sub>train</sub>) | 0.13 | 0.01 | 12.94 | < 0.0001 |

**Table S7.** Between-cell type estimated differences in panoptic quality (PQ) using expected marginal means. The p-value corresponds to a two-sided t-test for the linear model described in Equation 6. Standard errors (SE) were not possible to estimate for differences between neutrophils and other cell types.

| Dataset (*A*) | Dataset (*B*) | Estimate ($PQ_A - PQ_B$) | SE | t-ratio | p-value |
| --- | --- | --- | --- | --- | --- |
| Connective/stroma | Epithelial | -0.034 | 0.007 | -4.625 | 0.002 |
|  | Inflammatory | -0.274 | 0.010 | -28.029 | <0.0001 |
|  | Lymphocyte | -0.237 | 0.008 | -30.588 | <0.0001 |
|  | Macrophage | 0.133 | 0.040 | 3.339 | 0.03 |
|  | Neutrophil | 0.243 |  |  |  |
|  | Tumor/malignant | -0.120 | 0.010 | -11.494 | <0.0001 |
| Epithelial | Inflammatory | -0.241 | 0.012 | -20.439 | <0.0001 |
|  | Lymphocyte | -0.203 | 0.005 | -39.055 | <0.0001 |
|  | Macrophage | 0.167 | 0.039 | 4.232 | 0.004 |
|  | Neutrophil | 0.276 |  |  |  |
|  | Tumor/malignant | -0.087 | 0.011 | -7.798 | <0.0001 |
| Inflammatory | Lymphocyte | 0.037 | 0.012 | 3.093 | 0.03 |
|  | Macrophage | 0.408 | 0.041 | 9.929 | <0.0001 |
|  | Neutrophil | 0.517 |  |  |  |
|  | Tumor/malignant | 0.154 | 0.013 | 11.424 | <0.0001 |
| Lymphocyte | Macrophage | 0.370 | 0.039 | 9.385 | <0.0001 |
|  | Neutrophil | 0.480 |  |  |  |
|  | Tumor/malignant | 0.117 | 0.011 | 10.622 | <0.0001 |
| Macrophage | Neutrophil | 0.109 |  |  |  |
|  | Tumor/malignant | -0.254 | 0.041 | -6.229 | <0.0001 |
| Neutrophil | Tumor/malignant | -0.363 |  |  |  |

**Table S8.** Panoptic quality values for different cell types for the trained parameters ablation experiments (Methods).

| Dataset | Cell class | Class head | Neck and class head | Full network | Full network + U-Net |
| --- | --- | --- | --- | --- | --- |
| ConSep | Other | 0.000 | 0.080 | 0.244 | 0.248 |
|  | Inflammatory | 0.377 | 0.583 | 0.627 | 0.607 |
|  | Healthy epithelial | 0.000 | 0.078 | 0.570 | 0.413 |
|  | Malignant epithelial | 0.320 | 0.422 | 0.526 | 0.508 |
|  | Stroma | 0.000 | 0.277 | 0.415 | 0.384 |
|  | Muscle | 0.000 | 0.316 | 0.386 | 0.355 |
| GlySAC | Epithelial | 0.262 | 0.496 | 0.566 | 0.543 |
|  | Lymphocyte | 0.408 | 0.534 | 0.566 | 0.564 |
|  | Other | 0.000 | 0.289 | 0.379 | 0.366 |
| MoNuSAC | Neutrophil | 0.000 | 0.422 | 0.660 | 0.638 |
|  | Epithelial | 0.000 | 0.717 | 0.730 | 0.726 |
|  | Lymphocyte | 0.432 | 0.700 | 0.699 | 0.697 |
|  | Macrophage | 0.125 | 0.609 | 0.709 | 0.666 |
| PUMA | Epithelial | 0.000 | 0.081 | 0.726 | 0.634 |
|  | Lymphocyte | 0.463 | 0.606 | 0.681 | 0.682 |
|  | Other | 0.000 | 0.123 | 0.424 | 0.387 |
|  | Stroma | 0.002 | 0.244 | 0.348 | 0.363 |
|  | Tumor | 0.563 | 0.620 | 0.715 | 0.712 |
|  | Apoptosis | 0.000 | 0.148 | 0.216 | 0.206 |
|  | Endothelial | 0.000 | 0.108 | 0.332 | 0.317 |
|  | Histocyte | 0.000 | 0.335 | 0.485 | 0.492 |
|  | Melanophage | 0.000 | 0.187 | 0.338 | 0.321 |

**Table S9.** Per-class panoptic performance results for ablation experiments.

| Dataset | Cell class | Standard | No uncertainty weighing | No class oversampling | Train for 1000 epochs | Basic setup | Reduced augmentations | Best epoch selection |
| --- | --- | --- | --- | --- | --- | --- | --- | --- |
| ConSep | Other | 0.244 | 0.227 | 0.202 | 0.295 | 0.223 | 0.275 | 0.149 |
|  | Inflammatory | 0.627 | 0.629 | 0.626 | 0.636 | 0.633 | 0.632 | 0.618 |
|  | Healthy epithelial | 0.570 | 0.396 | 0.359 | 0.376 | 0.332 | 0.245 | 0.285 |
|  | Malignant epithelial | 0.526 | 0.507 | 0.502 | 0.517 | 0.504 | 0.501 | 0.506 |
|  | Stroma | 0.415 | 0.419 | 0.410 | 0.427 | 0.408 | 0.408 | 0.401 |
|  | Muscle | 0.385 | 0.391 | 0.383 | 0.363 | 0.246 | 0.253 | 0.374 |
| GlySAC | Epithelial | 0.566 | 0.562 | 0.558 | 0.549 | 0.554 | 0.559 | 0.518 |
|  | Lymphocyte | 0.566 | 0.560 | 0.562 | 0.565 | 0.557 | 0.563 | 0.554 |
|  | Other | 0.379 | 0.372 | 0.378 | 0.361 | 0.349 | 0.353 | 0.348 |
| MoNuSAC | Neutrophil | 0.660 | 0.648 | 0.651 | 0.666 | 0.653 | 0.651 | 0.638 |
|  | Epithelial | 0.730 | 0.731 | 0.733 | 0.728 | 0.722 | 0.730 | 0.717 |
|  | Lymphocyte | 0.699 | 0.701 | 0.702 | 0.697 | 0.682 | 0.692 | 0.690 |
|  | Macrophage | 0.709 | 0.690 | 0.679 | 0.668 | 0.651 | 0.674 | 0.663 |
| PUMA | Epithelial | 0.726 | 0.641 | 0.624 | 0.454 | 0.646 | 0.530 | 0.504 |
|  | Lymphocyte | 0.681 | 0.684 | 0.685 | 0.685 | 0.681 | 0.675 | 0.671 |
|  | Other | 0.424 | 0.454 | 0.466 | 0.402 | 0.310 | 0.433 | 0.408 |
|  | Stroma | 0.349 | 0.352 | 0.362 | 0.342 | 0.357 | 0.343 | 0.341 |
|  | Tumor | 0.715 | 0.716 | 0.715 | 0.710 | 0.705 | 0.709 | 0.708 |
|  | Apoptosis | 0.216 | 0.240 | 0.269 | 0.205 | 0.230 | 0.215 | 0.223 |
|  | Endothelial | 0.332 | 0.361 | 0.356 | 0.325 | 0.352 | 0.266 | 0.336 |
|  | Histocyte | 0.485 | 0.481 | 0.504 | 0.494 | 0.504 | 0.490 | 0.481 |
|  | Melanophage | 0.338 | 0.338 | 0.342 | 0.233 | 0.329 | 0.313 | 0.313 |

## S9 Supplementary Methods

### S9.1 Classpose for QuPath: GUI-assisted semantic segmentation

Classpose was developed featuring a simple and straightforward Python-based command-line interface for whole slide images. However, and while popular, programming and coding are not necessarily a part of the wider life sciences curriculum. As such, we developed an extension for the QuPath software program [20], a widely used software for digital histopathology with nearly 8,000 citations.

Our QuPath extension acts as a wrapper for the Python-based command-line interface. This tool allows for the specification of several different parameters:

– **Model** — models are selected from 5 default options, each trained on the datasets used throughout this work (CoNIC, ConSep, GlySAC, MoNuSAC, and PUMA; NuCLS was excluded due to its comparatively poorer performance). These models are hosted in HuggingFace and are automatically downloaded if locally unavailable.
– **Local models** — users might want to train their own Classpose models. For this, we allowed the user to specify a local model as a path to a YAML configuration file with the following fields: i) path, indicating the model path, url, indicating the model URL in case it is not locally available, mpp, the resolution in microns per pixel (MPP), and cell_types, the list of cell types to be considered (one for each class).
– **Test-time augmentation (TTA)** — we made use of Cellpose’s TTA methodology (rotations and flips) to improve the inference performance of Classpose.
– **Tissue detection** — to sample tiles from regions which have tissue, tissue detection using GrandQC [21] is performed and use this to i) select tiles for prediction and ii) exclude cells lying outside the tissue.
– **Artefact detection** — artefact detection is conducted using GrandQC [21] to enable post-hoc filtering by the user. This allows for users to manually inspect whether they would agree with the artefact predictions prior to removing cells contained within each artefact.
– **User-defined regions of interest (ROI)** — we allow users to define their own regions of interest. This is especially useful if users are interested in predictions contained exclusively within specific structures such as colonic crypts or glands. This comes with the added benefit of greatly accelerating inference when users are interested exclusively in smaller ROIs.
– **Output types** — we give users the option to select their preferred output format: CSV and SpatialData. Geo-JSON outputs are always included, and users could additionally choose to include CSV, SpatialData, both, or neither.
– **Device, GPU indices and data types** — users can select between different devices (CPU, MPS (i.e. Apple Silicon) and GPU) while also enabling the selection of multiple GPU indices for parallel inference. For users with multi-GPU setups, this could greatly speed up inference. Finally, we allow for bf16 specification, which could significantly speed up inference.
– **Advanced/compute specific parameters** — users can specify additional parameters such as the batch size, the tile size, the overlap between adjacent tiles, and minimum tissue area to be used to filter the detected tissue regions.

We exposed these parameters through our QuPath extension, and make detailed installation instructions available in the Classpose Github ^17^ under `qupath-extension-classpose`. .
.
Additionally, we also expose the tissue detection and artefact detection methods from GrandQC in case users require these simpler functionalities. Of note, tissue detection could be used as an initial step for inference and followed by the selection of relevant sections of tissue for user-defined ROI inference. The complete workflow, from parameter configuration to the resulting cell classification overlay within QuPath, is displayed in Figure S2.

### S9.2 Semantic classification sub-network

We allowed for Classpose to be extended using a U-Net-based [62] semantic segmentation head to the model. The input for this U-Net sub-network was the 256-channel feature maps as generated by the SAM-driven backbone of CP-SAM. It progressively transformed them through an encoder with channel expansions of 256 to 512 to 768, a 768-channel bottleneck, and a symmetric decoder that reduced channels back to 256 before outputting class-specific logits. Incorporating skip connections between encoder and decoder layers to preserve spatial details, this U-Net sub-network replaced the simpler 1 × 1 convolution as used in CP-SAM, allowing for more refined feature representations. This was introduced to allow Classpose to learn how to adequately transform features before using them for semantic classification.

### S9.3 Cross-dataset cell type mapping

The complete cell type mapping used for each dataset pair is as follows:

– CoNIC → ConSep
  - Neutrophil, Lymphocyte, Plasma cell, Eosinophil → Inflammatory
  - Connective → Stroma
  - Epithelial → Background
– CoNIC → GlySAC
  - Epithelial → Epithelial
  - Lymphocyte → Lymphocyte
  - Neutrophil, Plasma cell, Eosinophil, Connective → Background
– CoNIC → MoNuSAC
  - Neutrophil → Neutrophil
  - Epithelial → Epithelial
  - Lymphocyte → Lymphocyte
  - Plasma cell, Eosinophil, Connective → Background
– CoNIC → PUMA
  - Epithelial → Epithelial
  - Lymphocyte → Lymphocyte
  - Connective → Stroma
  - Neutrophil, Plasma cell, Eosinophil → Background
– ConSep → CoNIC
  - Healthy epithelial, Malignant epithelial → Epithelial
  - Other, Inflammatory, Stroma, Muscle → Background
– ConSep → GlySAC
  - Healthy epithelial, Malignant epithelial → Epithelial
  - Other, Inflammatory, Stroma, Muscle → Background
– ConSep → MoNuSAC
  - Healthy epithelial, Malignant epithelial → Epithelial
  - Other, Inflammatory, Stroma, Muscle → Background
– ConSep → PUMA
  - Healthy epithelial → Epithelial
  - Malignant epithelial → Tumor
  - Stroma → Stroma
  - Other, Inflammatory, Muscle → Background
– GlySAC → CoNIC
  - Lymphocyte → Lymphocyte
  - Epithelial → Epithelial
  - Other, Ambiguous → Background
– GlySAC → ConSep
  - Lymphocyte → Inflammatory
  - Other, Epithelial, Ambiguous → Background
– GlySAC → MoNuSAC
  - Lymphocyte → Lymphocyte
  - Epithelial → Epithelial
  - Other, Ambiguous → Background
– GlySAC → PUMA
  - Lymphocyte → Lymphocyte
  - Epithelial → Epithelial
  - Other, Ambiguous → Background
– MoNuSAC → CoNIC
  - Epithelial → Epithelial
  - Lymphocyte → Lymphocyte
  - Neutrophil → Neutrophil
  - Macrophage → Background
– MoNuSAC → ConSep
  - Lymphocyte, Macrophage, Neutrophil → Inflammatory
  - Epithelial → Background
– MoNuSAC → GlySAC
  - Epithelial → Epithelial
  - Lymphocyte → Lymphocyte
  - Macrophage, Neutrophil → Background
– MoNuSAC → PUMA
  - Epithelial → Epithelial
  - Lymphocyte → Lymphocyte
  - Macrophage, Neutrophil → Background
– PUMA → CoNIC
  - Lymphocyte → Lymphocyte
  - Epithelial → Epithelial
  - Apoptosis, Tumor, Endothelial, Stroma, Histocyte, Melanophage, Other → Background
– PUMA → ConSep
  - Tumor → Malignant epithelial
  - Stroma → Stroma
  - Lymphocyte, Histocyte → Inflammatory
  - Apoptosis, Endothelial, Epithelial, Melanophage, Other → Background
– PUMA → GlySAC
  - Lymphocyte → Lymphocyte
  - Epithelial → Epithelial
  - Apoptosis, Tumor, Endothelial, Stroma, Histocyte, Melanophage, Other → Background
– PUMA → MoNuSAC
  - Lymphocyte → Lymphocyte
  - Histocyte, Melanophage → Macrophage
  - Epithelial → Epithelial
  - Apoptosis, Tumor, Endothelial, Stroma, Other → Background

### S9.4 Annotation protocol

Regions of invasive colorectal cancer (CRC) from all publicly available haematoxylin-and-eosin (H&E)-stained slides (n=427) were expert-annotated by a board-certified pathologist (KB) using the brush tool in QuPath version 0.6. Adjacent colorectal mucosa and dysplasia of any grade were excluded from annotation if present on slide. Fat, smooth muscle, necrosis, and any technical artefacts were excluded wherever possible and when not tumour-infiltrated. Furthermore, lymph node metastases were not annotated. Mucinous cancers were annotated to include the cancer epithelium surrounding mucin extravasates (“pools”). It should be noted that the dataset also contained CRC that had infiltrated the appendix wall, and that the amount/area of invasive cancer varied across the cohort. Furthermore, some cancers exhibited a clearly expansile and nodular configuration, while others were more patchy and infiltrative, which necessitated multi-region annotation in some slides.

The SurGen dataset contains a variety of growth patterns and morphologies. For example, it includes cases of signet ring cell invasion (case 294), strong desmoplastic regions (case 189), and cases with a larger adjacent adenoma (case 309) or infiltration of the appendix wall (case 498). Some cases were excluded from annotation and subsequent analysis due to the absence of invasive cancer (cases 14 and 410), the presence of pure lymph node metastases (cases 81, 148, 164 and 406), or uncertainty regarding the diagnosis of colorectal adenocarcinoma (case 205). The limitations of the SurGen dataset include inconsistent staining quality and human annotations that are subject to interobserver variability.

### S9.5 Spatial features

We computed three distinct spatial feature families to characterise cell type relationships: the **pair correlation function (PCF), nearest-neighbour distance (NND)**, and *k***-nearest-neighbour (***k***-NN) neighbourhood composition**.

For each input sample, the Classpose output is reduced to a finite marked point pattern

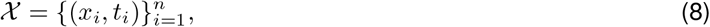

where 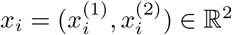 denotes the centroid of cell *i* in cartesian coordinates, *t*_*i*_ ∈ T is its cell-type label, and *n* is the total number of cells in the sample or case under consideration. In the SurGen results,

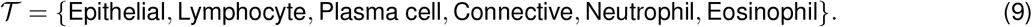

When a sample contained multiple disconnected region-of-interest, the analysis domain is defined as

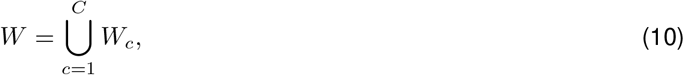

where each *W*_*c*_ denotes one connected component. All three feature families are computed separately within each *W*_*c*_. If *θ*_*c*_ denotes any component-level summary and area(*W*_*c*_) the corresponding component area, the sample-level summary is calculated by

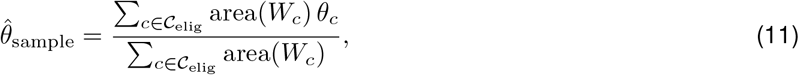

where *C*_elig_ denotes the set of components for which the summary is defined. A pair (*A, B*) was evaluated only in connected components containing at least 15 cells of each label.

#### Pair correlation function (PCF)

The pair correlation function is a second-order summary of a spatial point process. For two cell types *A, B* ∈ *T*, the corresponding type-specific subpatterns are *X*_*A*_ ={ *x*_*i*_ : (*x*_*i*_, *t*_*i*_) ∈ *X, t*_*i*_ = *A* and *X*_*B*_ = *x*_*i*_ : (*x*_*i*_, *t*_*i*_) ∈ *X, t*_*i*_ = *B*}. The PCF then asks whether cells of type *B* occur around cells of type *A* more often or less often than expected under complete spatial randomness (CSR) after conditioning on the marginal intensity of *B*. Values *ĝ*_*AB*_(*r*) *>* 1 indicated local enrichment at scale *r*, in contrast 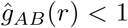 indicates depletion or exclusion.

In the SurGen analysis, PCF served as the principal scale-resolved statistic because it retains information across a range of biologically plausible interaction radii instead of collapsing the signal to a single number.

Within one component *W*_*c*_, the type-specific subpatterns are denoted by 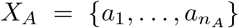 and 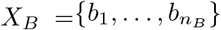, where *n*_*A*_ = |*X*_*A*_| and *n*_*B*_ = |*X*_*B*_|. We use annuli of fixed width *w* = 0.010 mm and

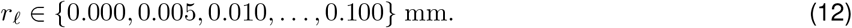

The *ℓ*^th^ annulus is [*r*_*ℓ*_, *r*_*ℓ*_ + *w*), where *ℓ* indexes the annulus grid. With *λ*_*B*_ the intensity of type *B* inside *W*_*c*_, the estimated PCF is defined as

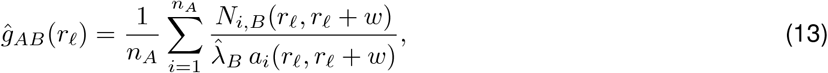

where

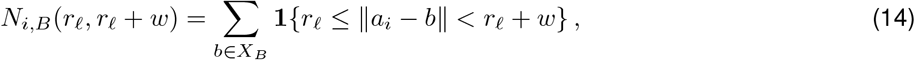

And

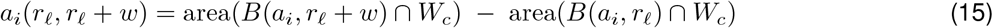

is the annulus area available within the observed component. Here *N*_*i,B*_(*r*_*ℓ*_, *r*_*ℓ*_ + *w*) is the number of type *B* cells lying in the annulus around the focal point *a*_*i*_, and *B*(*a, r*) denotes the disc of radius *r* centered at *a*.

For cross-PCF, we estimate

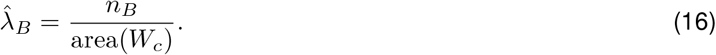

Here *n*_*B*_ = |*X*_*B*_ | is the number of type-*B* cells in the component and area(*W*_*c*_) is the area of the component. For auto-PCF (*A* = *B*), self-pairs must be excluded and the intensity becomes

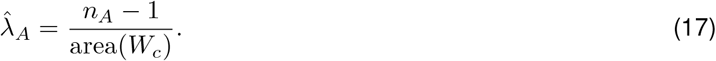

For descriptive summaries of cross-type organisation, it is convenient to represent the pair *A, B* by a single symmetric curve. However, Eq. (13) is directional because conditioning on type *A* and conditioning on type *B* involve different marginal intensities. A symmetric summary is therefore obtained from the pointwise average

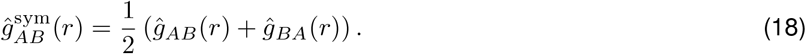

This yields a single symmetric curve for the unordered pair *A, B* .

As the PCF is a function of distance rather than a single scalar quantity, sample-level comparison is facilitated by band-specific signed areas relative to the CSR baseline as

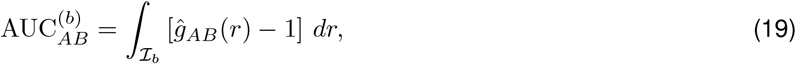

Subtracting 1 centers the integrand at CSR, so 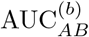 *b* indexes one of three prespecified distance bands and I_*b*_ denotes the corresponding interval

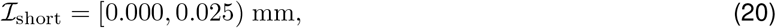

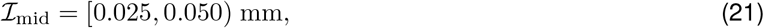

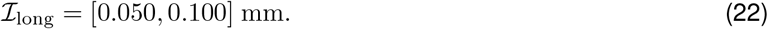

We approximate the integral by the trapezoidal rule over the discrete inner-radius grid. Positive values indicate clustering above CSR over that interval and negative values indicate exclusion.

#### Nearest-neighbour distance (NND)

NND measures the most immediate contact scale. For a focal type *A* and target type *B*, the directed NND distribution asks*‘how far must each A cell travel to encounter its nearest B cell?’*.

For each *a*_*i*_ ∈ *X*_*A*_, the directed nearest-neighbour distance to type *B* is

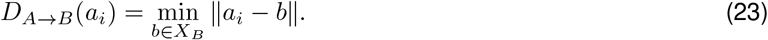

For auto-NND, the self-match must be excluded:

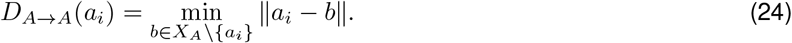

We summarise each NND pair by four robust scalar quantities

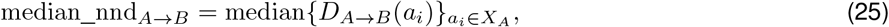

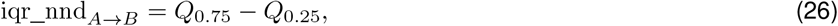

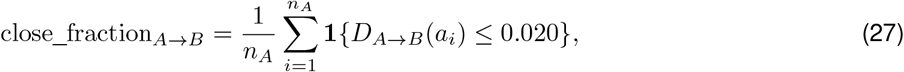

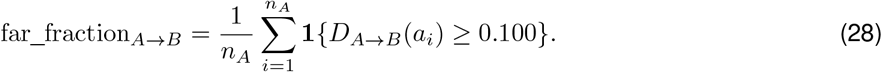

Here *Q*_0.25_ and *Q*_0.75_ denote the empirical first and third quartiles of the directed NND distribution. The close_fraction threshold serves as a close-contact proxy, while the far_fraction quantifies exclusion.

#### *k*-nearest neighbours (*k*-NN)

We estimated neighbourhood composition around each cell using *k* = 10 nearest neighbours. For a component containing *n* cells in total, *N*_*k*_(*x*_*i*_) denotes the set of the *k* = 10 nearest neighbours of cell *i*, excluding the cell itself. If *t*_*j*_ is the label of neighbour *x*_*j*_, the empirical local proportion of type *B* around cell *i* is

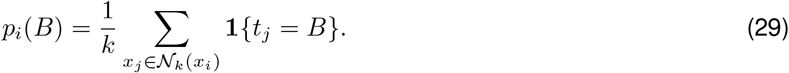

Thus *p*_*i*_(*B*) is the fraction of the *k* neighbours of cell *i* that belong to phenotype *B*. For a focal label *A*, the observed average fraction of *B* among the neighbours of *A* cells is

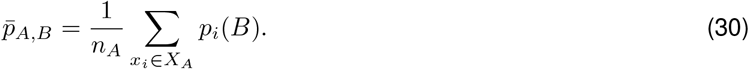

The expected background frequency of *B* within the component is taken as the global composition

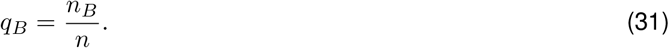

The resulting standardized enrichment is

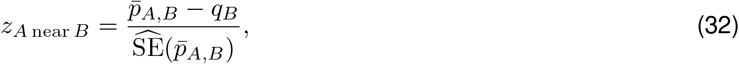

where 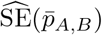 is the standard error of the per-cell neighbour fractions among focal *A* cells,

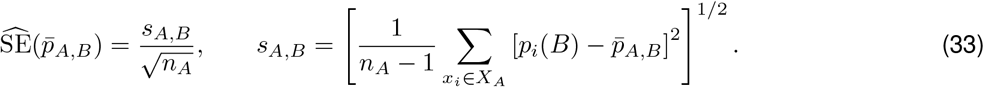

*s*_*A,B*_ is the sample standard deviation of the cell-wise neighbourhood fractions *p*_*i*_(*B*) among focal *A* cells. Positive values in Eq. (32) indicate that *B* is over-represented in the immediate neighbourhood of *A* beyond what would be expected from the component-wide class prevalence.

The per-cell neighbourhood proportions {*p*_*i*_(*t*)}_*t*∈T_ are also used to compute niche diversity. For a cell *x*_*i*_,

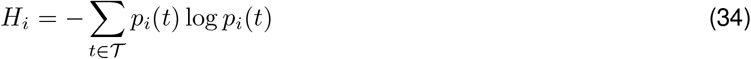

is the Shannon entropy of the 10-neighbour label distribution. This quantity is then averaged over all focal cells of type *A*:

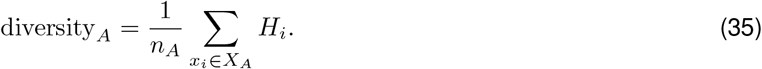

High values indicate mixed microenvironments and low values indicate homogeneous local niches.

We also include neighbourhood immune-fraction summaries around epithelial cells, asa epithelial–immune interaction is of high interest. The immune label set is *T*_imm_ = {Lymphocyte, Plasma cell, Eosinophil, Neutrophil} . For each epithelial cell *x*_*i*_,

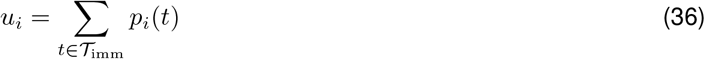

is the fraction of immune neighbours in its 10-NN niche. Both the mean and median of the epithelial-cell-specific values {*u*_*i*_} are then aggregated to the sample level.

## S10 Supplementary Results

### S10.1 Classpose discovers spatial and spatial-morphological signals in SurGen

#### MMR deficiency: immune enrichment, connective dispersal, and elevated niche diversity

MMR-deficient (dMMR) colorectal cancer carries a high mutational burden that renders tumours immunogenic, driving the dense lymphocytic infiltration that is well established [31,32,33]. Classpose resolves three quantitative spatial correlates of this biology at single-cell resolution (Figure 6).

##### Immune cells are found further from connective cells and nearer to epithelial cells

In dMMR tumours, lymphocytes and plasma cells lie further from connective cells and closer to epithelial cells, with fewer close-range immune-connective contacts, an elevated fraction of close-range immune-epithelial contacts confirmed at near and intermediate distances by pair-correlation analysis, and reduced depletion of all non-epithelial cell types from the epithelial neighbourhood. The proximity of lymphocytes to the cancer epithelium is broadly consistent with the dense intratumoral T cell infiltration established in dMMR CRC by immunohistochemistry and single-cell profiling [31,32,33]. As H&E cannot distinguish lymphocyte subtypes, the lymphocytes detected here likely include T cells among other subtypes. Plasma cells show an equivalent spatial redistribution toward epithelial cells and away from connective cells. Epithelial niche diversity, which is the Shannon entropy of the 10-nearest-neighbour label distribution, is correspondingly elevated, providing a single-summary measure of the overall increase in neighbourhood heterogeneity captured by the pairwise features above.

##### Connective cells show greater distances from one another and reduced distances from epithelial cells

Connective cells (a heterogeneous class spanning fibroblasts, muscle, and endothelial cells) show greater inter-cellular distances and fewer close-range connective-connective contacts in dMMR tumours, while lying closer to epithelial cells across all three pair-correlation distance bands. We interpret this as disruption of the organised connective-cell niche, an interpretation consistent with the stromal remodelling reported in dMMR tumours by spatial transcriptomics [32].

##### Cell shape changes at the cancer border

The perimeters of epithelial cells, plasma cells, and lymphocytes all decrease more steeply near the cancer ROI boundary in dMMR samples (Figure S12). Plasma-cell solidity (area divided by convex-hull area) declines more slowly near the cancer border in dMMR samples, indicating a less steep loss of convexity approaching the boundary. These per-cell shape gradients are, to our knowledge, novel. Prior work has quantified immune-cell density as a continuous function of distance from the cancer border [35]. Gradients in histological phenotype and protein marker expression across CRC tissue have also been described [36]. However, continuous per-cell shape gradients measured as a function of distance from the cancer border have not, to our knowledge, been reported.

#### BRAF ^MUT^: immune enrichment near cancer cells, connective-cell dispersal, and tighter epithelial–connective proximity

*BRAF* ^MUT^ CRC shares many spatial features with dMMR tumours. Connective cells are further from one another and immune cells are further from connective cells. Epithelial cells are in closer proximity to lymphocytes, plasma cells, and connective cells, with connective cells less depleted from the epithelial neighbourhood. Pair-correlation analysis confirms epithelial co-localisation with connective cells (near and intermediate distances) and with lymphocytes and plasma cells (near distances), alongside epithelial self-clustering at far distances. The lymphocyte perimeter decreases more steeply near the cancer border, and both the immune fraction in the epithelial niche and epithelial niche diversity are elevated.

We note that *BRAF* ^MUT^ and MMR deficiency co-occur in a substantial proportion of cases in this cohort, consistent with their known association [23]. Features specific to the combined condition are examined in the dMMRx*BRAF* ^MUT^ paragraph below. The shared features above may therefore partly reflect the common dMMR status of the same cases rather than an independent effect of *BRAF* ^MUT^, and the conditional-inference model accounts for this cooccurrence statistically but cannot fully disentangle the two contributions. One spatial pattern is specific to *BRAF* ^MUT^ and absent in dMMR alone: epithelial cells lie closer to connective cells, with lower variability in that distance.

#### *KRAS*^MUT^: connective-cell elongation at the cancer border

*KRAS*^MUT^ CRC shows a much narrower set of associations than dMMR or *BRAF* ^MUT^ tumours, concentrated almost entirely in connective-cell morphology near the cancer border. Connective cells near the cancer border show progressively more elongated outlines and greater perimeters while their solidity falls. This combination reflects a longer, more irregular outline rather than simple enlargement. A lower fraction of epithelial cells also lie close to connective cells in *KRAS*^MUT^ samples, consistent with reports that *KRAS* signalling in cancer cells modulates the activity of surrounding fibroblasts and depletes the extracellular matrix [34], though that study was conducted in MSS locally advanced rectal cancer and its generalisation to the mixed CRC cohort studied here is an extrapolation.

#### *NRAS*^MUT^: connective- and plasma-cell elongation at the cancer border

*NRAS* mutations are uncommon in CRC (≈ 4%) [23] and show the fewest associations in our data. No significant associations were recovered in the spatial feature classes (nearest-neighbour distances, neighbourhood composition, pair-correlation). The detectable signal is confined to morphological gradients at the border. Connective cells have more elongated and less convex outlines near the boundary, mirroring the pattern seen in *KRAS*^MUT^, while plasma cells elongate at a higher rate. These observations should be treated as exploratory given the small sample size (*n* = 15).

#### dMMRx*BRAF* ^MUT^ : a broadly shared programme with emergent immune-niche heterogeneity

The dMMRx*BRAF* ^MUT^ interaction comprises a large core of features shared with one or both parent conditions, plus a small set significant specifically in the combined dMMR+*BRAF* ^MUT^.

##### Shared features

Given their known co-occurrence [23], it is expected that the dMMRx*BRAF* ^MUT^ interaction group shares many of the features of both parent conditions. Immune enrichment near the cancer epithelium, elevated immune fraction, connective-cell dispersal, and the lymphocyte perimeter gradient at the cancer border are all present in dMMRx*BRAF* ^MUT^ . Notably, the bulk immune-connective dispersal metrics (median and IQR of lymphocyte- and plasma-cell-to-connective distances), significant in both parent conditions individually, do not reach significance in dMMRx*BRAF* ^MUT^, suggesting the interaction signal is concentrated at the close-range extreme rather than across the full distribution. Other features are inherited specifically from dMMR (connective cells less self-enriched, more lymphocytes and plasma cells in close proximity to epithelial cells) or from *BRAF* ^MUT^ (epithelial cells closer to connective cells, with lower variability).

##### Emergent features

Three features reach significance in dMMRx*BRAF* ^MUT^ but in neither parent alone (Figure 6). First, the fraction of epithelial cells far from their nearest connective neighbour is reduced. Combined with the elevated close-contact fraction shared with both parents, this compression of the epithelial-to-connective distance distribution toward shorter ranges suggests more thorough intermingling of the two compartments than either parent produces alone. Second and third, niche diversity is elevated not only for epithelial cells (shared with both parents) but independently for lymphocytes and plasma cells, indicating that immune cells in the combined context occupy more heterogeneous local microenvironments. These three features could reflect either a genuine emergent biological state specific to the combined dMMR+*BRAF* ^MUT^ context, or a statistical effect in which cases carrying both alterations are simply at the extreme end of distributions that do not individually reach significance in either parent condition alone. Resolving this would require experimental or longitudinal data beyond the scope of this study.

#### Classpose-driven spatial-morphological hallmarks

Together, the signatures recovered by Classpose reveal a hierarchical structure of molecular-phenotype associations in colorectal cancer. dMMRx*BRAF* ^MUT^ dominate, producing an extensive reorganisation of the spatial relationships between immune, epithelial, and connective cells, characterised by immune enrichment near cancer cells, connective dispersal, and elevated niche heterogeneity. We note that *BRAF* ^MUT^ and dMMR co-occur in a substantial proportion of cases in this cohort, consistent with their known association [23]. The contributions of dMMR and *BRAF* ^MUT^ therefore cannot be fully disentangled even with a conditional-inference model. *KRAS*^MUT^ and *NRAS*^MUT^ produce a narrower, morphology-confined signature, restricted to connective-cell shape changes near the cancer border. That these cell-type-resolved spatial signatures are recoverable directly from routine H&E illustrates the potential of Classpose for hypothesis generation and biological discovery at WSI scale.

## Footnotes

6 https://huggingface.co/classpose/classpose

7 available in https://conic-challenge.grand-challenge.org/Data/, accessed on May 20th, 2025

8 available in https://www.kaggle.com/datasets/karthikperupogu/consep, accessed on May 20th, 2025

9 available in https://sites.google.com/view/nucls/single-rater?authuser=0 as the “Corrected single-rater dataset”, accessed on May 20th, 2025

10 available in https://monusac-2020.grand-challenge.org/Data/, accessed on July 11th, 2025

11 availabe in https://github.com/MouseLand/cellpose/blob/main/paper/cpsam/semantic.py, accessed on July 11th, 2025

12 available in https://drive.google.com/file/d/1g1_xYFWgp3cRLKrlSwD2U5JDjooC0yHp/view, accessed on July 11th, 2025, and made available by the authors of DiffMix [44]

13 available in https://puma.grand-challenge.org/dataset/, accessed on July 15th, 2025

14 https://github.com/stardist/stardist/tree/conic-2022, accessed on July 15th, 2025

15 https://drive.google.com/drive/folders/1ujtMcxAr5kYYuvnbglfYZZnRH3ZOli79, accessed on September 2nd, 2025

16 https://github.com/MouseLand/cellpose/blob/main/paper/CP-SAM/semantic.py, accessed on October 1st 2025

17 https://github.com/sohmandal/classpose

